## Supplementary figures for "Analysis of diverse eukaryotes suggests the existence of an ancestral mitochondrial apparatus derived from the bacterial type II secretion system"

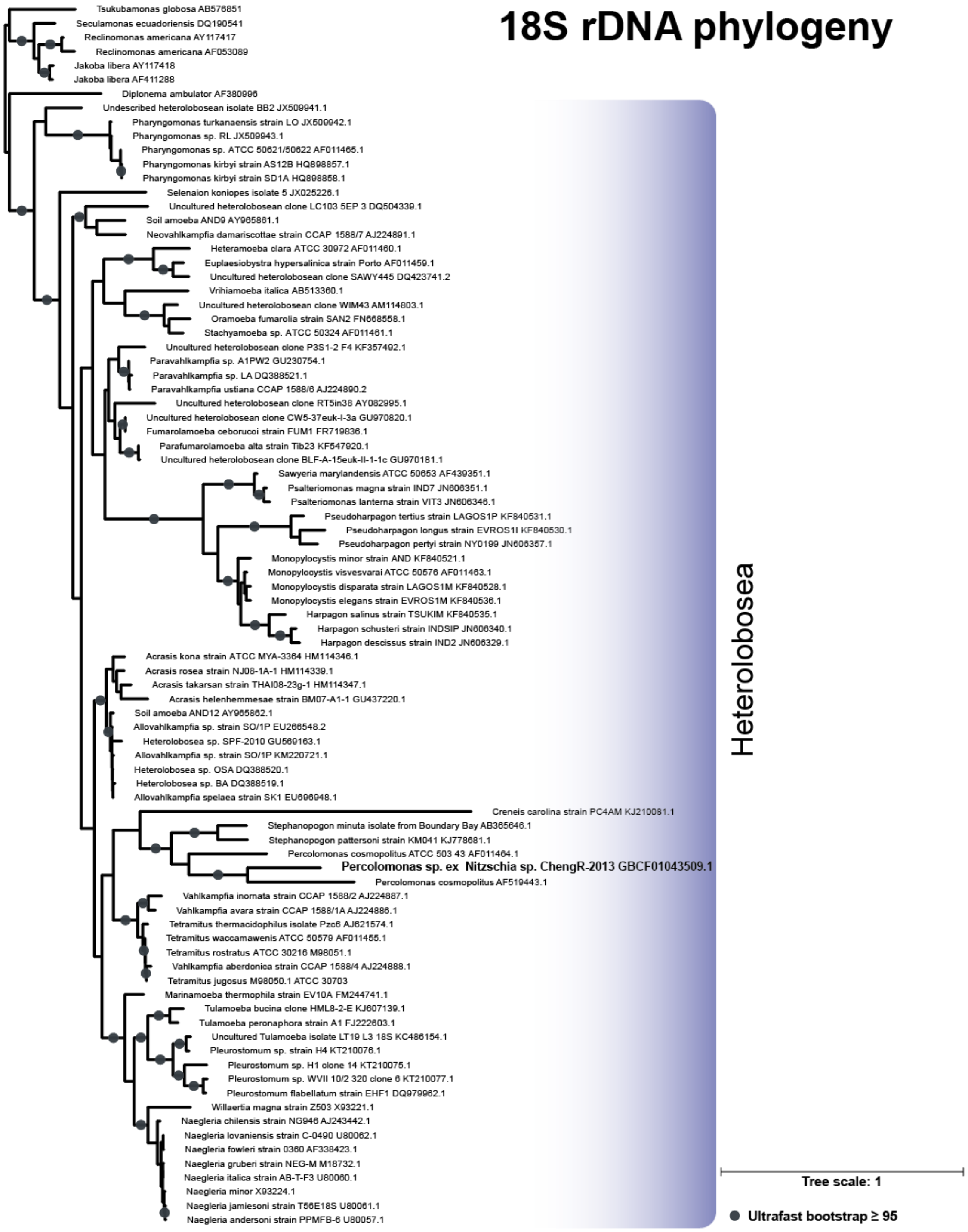


**Supplementary Fig. 1** Phylogenetic analysis of 18S rRNA gene from Heterolobosea showing the position of three *Percolomonas* species for which transcriptome data are available. The ML tree was calculated using IQ-TREE (substitution model TIM2+I+G4) with 1000 ultrafast bootstraps. The two taxa currently identified as two strains (AE-1 and WS) of the same species of *Percolomonas cosmopolitus* and used for generating transcriptome assemblies in the context of the MMETSP project (MMETSP0758 and MMETSP0759, respectively; https://www.imicrobe.us/#/projects/104) are in fact very different at the level of the 18S rRNA gene sequence and apparently represent two different species. In addition, we found out that the transcriptome assembly of the diatom *Nitzschia* sp. ChengR-2013 (NCBI BioProject PRJNA243394) is massively contaminated by sequences from a heterolobosean that according to the phylogenetic analysis of the 18S rRNA sequence present in the assembly is specifically related to *P. cosmopolitus* strain WS, yet again sufficiently different at the sequence level to be considered a separate species (provisionally denoted as *Percolomonas* sp. ex *Nitzschia* sp. ChengR-2013).

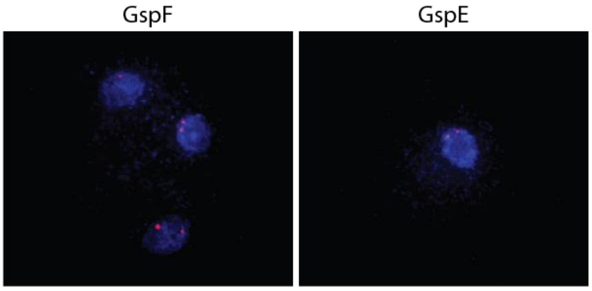


**Supplementary Fig. 2** Fluorescence *in situ* hybridization (FISH) of selected Gsp genes in *N. gruberi* nuclei. Probes specific for the *Ng*GspE and *Ng*GspF genes each label two loci (red) in diploid nuclei (blue) of *N. gruberi*. DNA was stained with DAPI, blue dots correspond to mitochondrial DNA.

A **GspD**

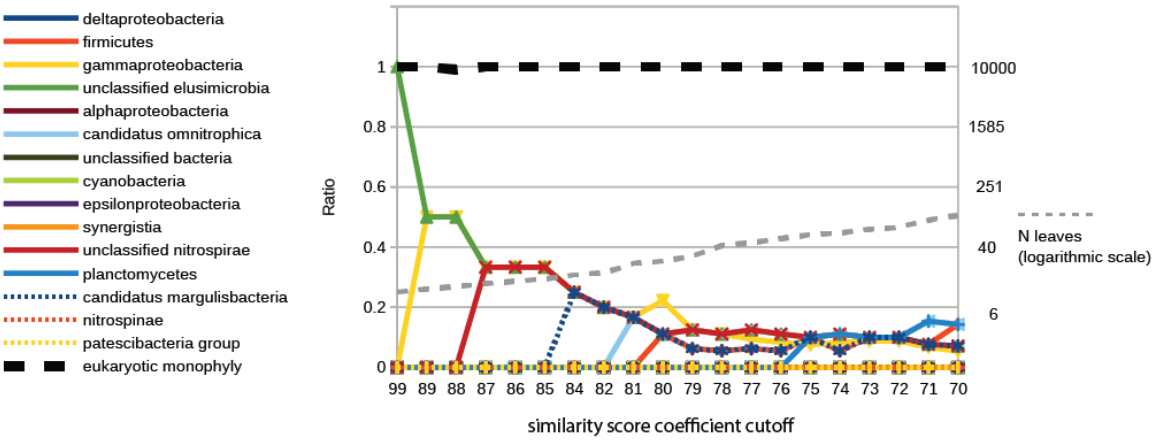


GspD - 75 AA positions

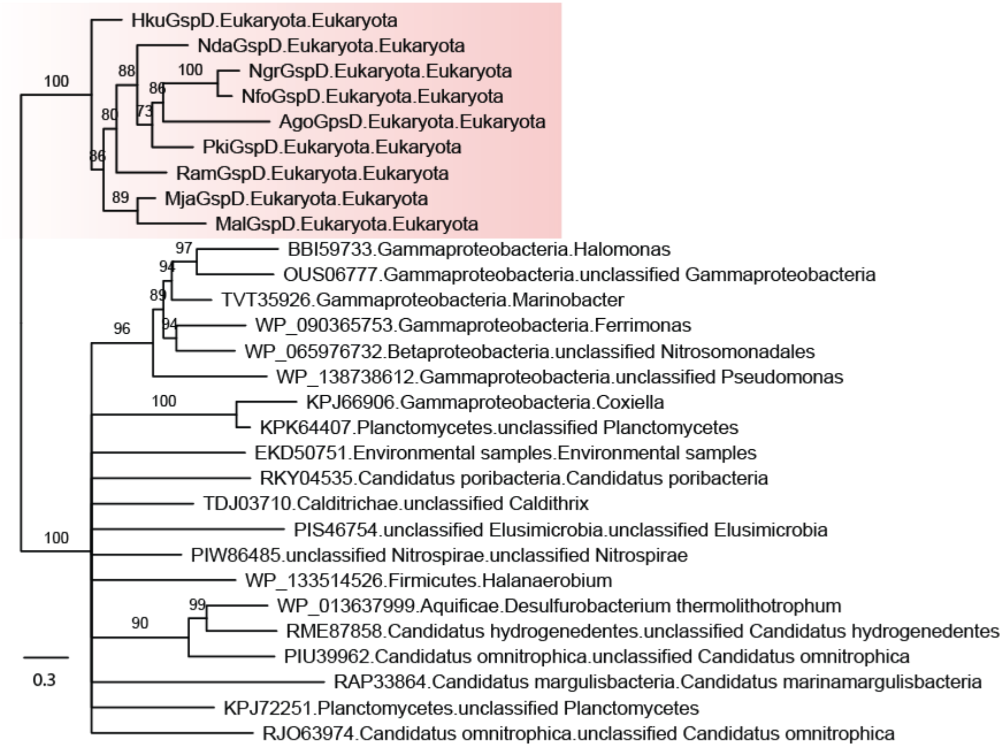


**Supplementary Fig. 3A** Phylogenetic relationship of the eukaryotic GspD proteins to prokaryotic homologues. For further explanations see page 7.

B **GspE**

GspE - 158 AA positions

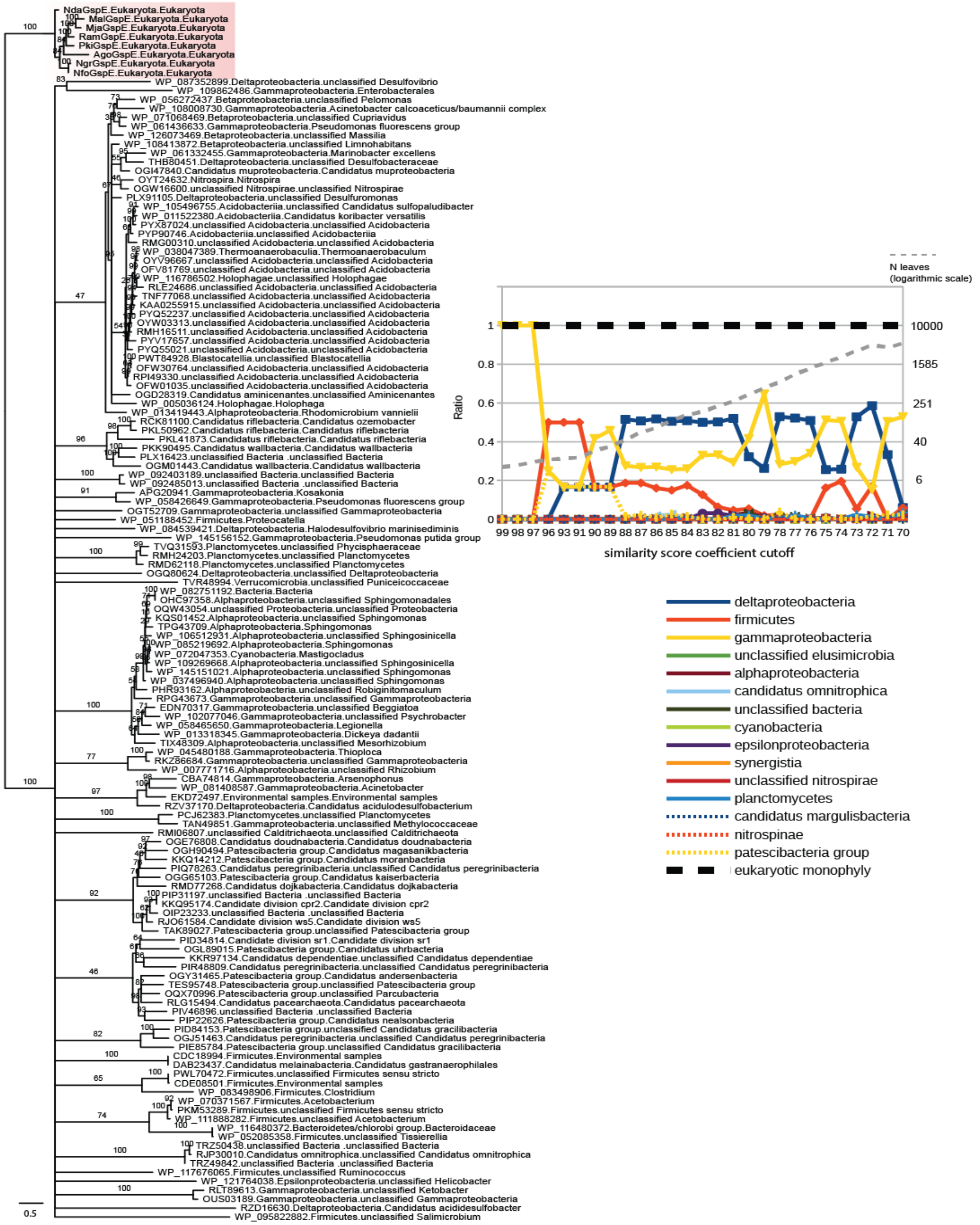


**Supplementary Fig. 3B** Phylogenetic relationship of the eukaryotic GspE proteins to prokaryotic homologues. For further explanations see page 7.

C GspF

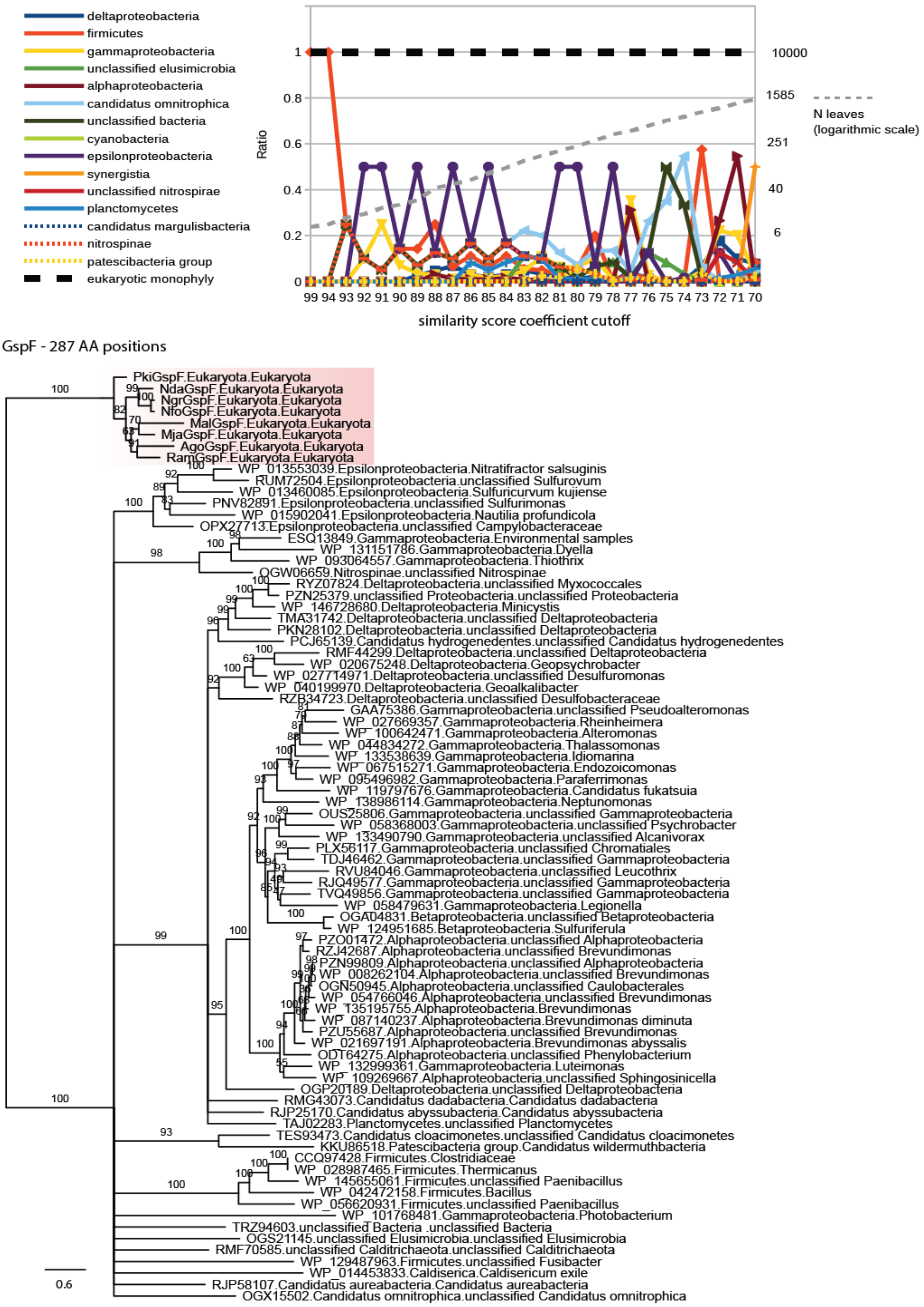


**Supplementary Fig. 3C** Phylogenetic relationship of the eukaryotic GspE proteins to prokaryotic homologues. For further explanations see page 7.

D **GspG**

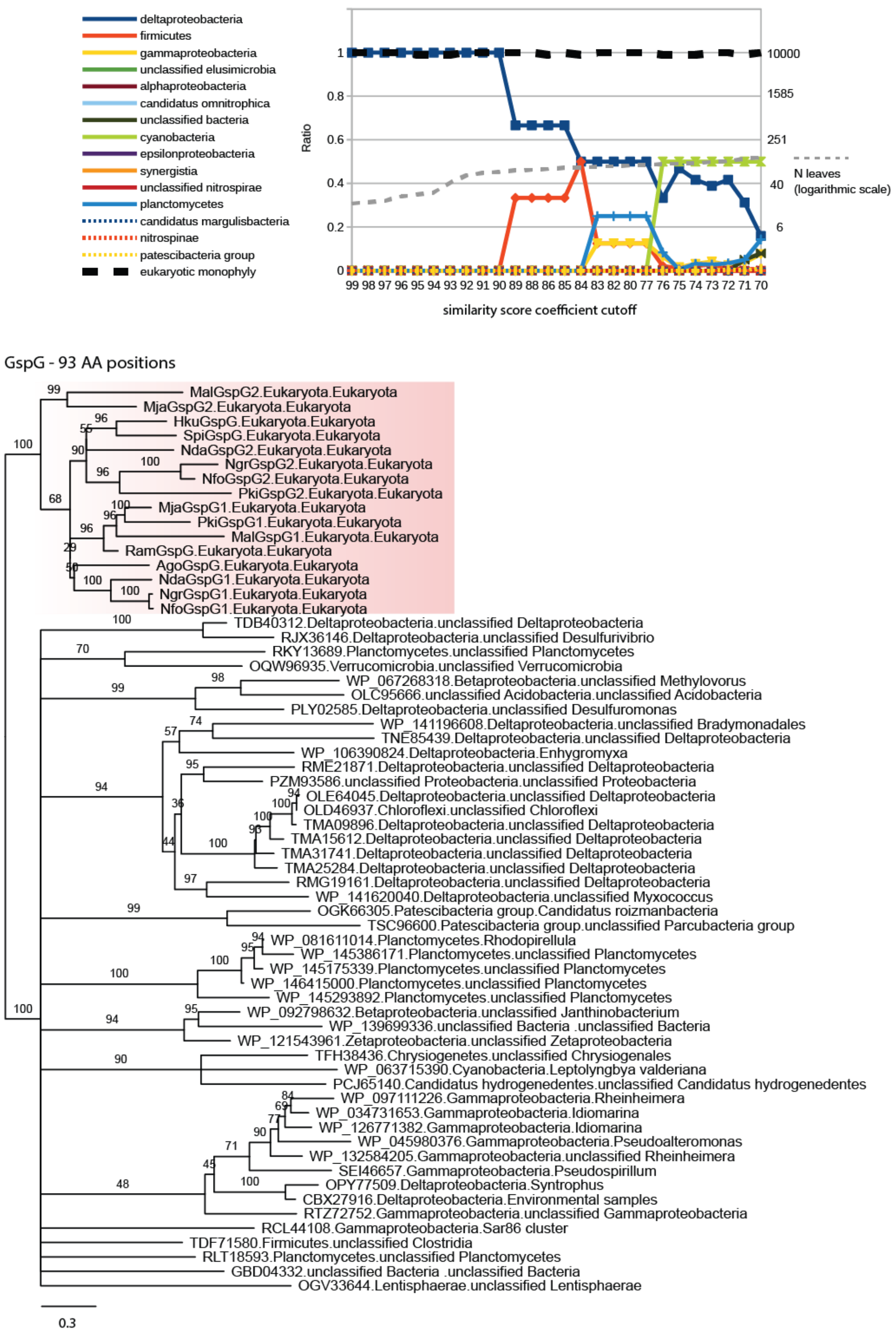


**Supplementary Fig. 3D** Phylogenetic relationship of the eukaryotic GspG proteins to prokaryotic homologues. For further explanations see page 7.

**Supplementary Fig. 3** (pg. 3-6) Phylogenetic relationship of the eukaryotic GspD (panel A), GspE (panel B, pg. 4), GspF (panel C, pg. 5), and GspG (panel D, pg. 6) proteins to prokaryotic homologues. **Top part of each panel:** The monophyly of eukaryotic Gsp sequences and their relation to the prokaryotic homologues was systematically analyzed by building phylogenetic trees (using IQ-TREE) from gradually expanding datasets. Specifically, the different sets of sequences were defined by varying the HMMER score cutoff (in searches using profile HMMs built for the eukaryotic Gsp families) based on the formula $score_{cutoff} = c*score_{best\ prokaryotic\ hit}$, with the coefficient c decreasing from 0.99 to 0.70 incrementally by 0.01 (X axis). The resulting trees were systematically analyzed for support of the monophyly of eukaryotic sequences and for the taxonomic assignment of the parental prokaryotic node of the eukaryotic subtree (details of the procedure are described in Materials and Methods). The Y axis of the plot simultaneously denotes three variables: (i) support of the monophyly of eukaryotic sequences (black dashed line; percentages are rescaled to values 0 to 1); (ii) the taxonomic affiliation of the parental prokaryotic node of the eukaryotic subtree plotted as a fraction (0-1) for each prokaryotic taxonomic group (lines and shapes of a different colour, see the graphical legend to the right); and (iii) number of leaves (individual sequences) in the respective tree (grey dashed line; the values rescaled to their thousandth). **Bottom part of each panel:** Exemplar phylogenetic trees based on datasets defined by particular HMMER score cutoffs (dotted arrow). The trees were arbitrarily rooted between the clade of eukaryotic Gsp homologues (highlighted by a light red background) and prokaryotic sequences. The leaves are described accordingly: “organism name”.”main taxon”.”sequence ID”.”system classification”. The “system classification” has been assigned using the models of subunits of T4P superfamily of molecular machines[1]. Ultra-fast bootstrap values are shown at internal branches.

### (A) GspD and GspDL

```
                                 10        20        30        40        50        60        70        80        90       100       110       120       130
                         ....|....|....|....|....|....|....|....|....|....|....|....|....|....|....|....|....|....|....|....|....|....|....|....|....|....|
A.godoyi D               ---------------------------------------------------------------------------------------------------------------------------------- 1
R.americana D            ---------------------------------------------------------------------------------------------------------------------------------- 1
M.jakobiformis D         MSQH------------------------------------------------------------------------------------------------------------------------------ 4
G.okellyi D              MS-------------------------------------------------------------------------------------------------------------------------------- 2
H.kukwesjijk D           ---------------------------------------------------------------------------------------------------------------------------------- 1
N.gruberi D              ---------------------------------------------------------------------------------------------------------------------------------- 1
N.fowleri D              MR-------------------------------------------------------------------------------------------------------------------------------- 2
Neo.damariscottae D      ---------------------------------------------------------------------------------------------------------------------------------- 1
P.cosmopolitus AE1 D     ---------------------------------------------------------------------------------------------------------------------------------- 1
P.kyrbyi D               ---------------------------------------------------------------------------------------------------------------------------------- 1
P.cosmopolitus WS D      -------------------------------------------------------------P---TCLKKSHPQKNY----------------------------------------------------- 13
E.coli                   MFWR--DMTLSIWRKKTTGL---------------------------------------------KTKKRLLPLM---LA--------------------AALCSSPV--------------WAEEATFT 46
V.cholerae               M---------KYWLKKSSWL----------------------------------------------------------LA--------------------GSLLSTPL---------------AMANEFS 28
A.salmonicida            MINK-----GKSWRLAT-------------------------------------------------------------VA--------------------AALMMAGS---------------AWATEYS 29
P.aeruginosa             MSQP---------LLRALFA---------------------------------------------PSSRSYVPAVLLSLAL--------GIQAAHAENSGGNAFVPAGN--------------QQEAHWT 54
N.meningtitis            MNTKLTKIISGLFVATAAFQT---------ASAGNITDIKVSSLPNKQKIVKVSFDKEIVNP---TGFVTSSPAR---IALDF---EQTGISMDQQVLEYADPLLSKISAAQNSSRARLVLNLNKPGQYN 112
Myxococcus               MLEE-SAVTRGKWMLAAAWAVVLVGARVHGAELNTLRGLDVSRTGSGAQVVVTG-----TRPPTFTVFRLSGPER---LVVDLSSADATGIKGHHEGSGPVSGVVASQFSDQRASVGRVLLALDKASQYD 121
S.enterica               -------------LYNKNY-----------------------------------------------------PLR------------------------------------------------------- 9
A.godoyi DL              -------------------------------------------------------------------------MRRTDLA-------------------------------------------------- 7
R.americana DL           ---------------------------------------------------------------------------------------------------------------------------------- 1
N.gruberi DL             ---------------------------------------------------------------------------------------------------------------------------------- 1
N.fowleri DL             ---------------------------------------------------------------------------------------------------------------------------------- 1
P.cosmopolitus AE1 DL    ---------------------------------------------------------------------------------------------------------------------------------- 1
Neo.damariscottae DL     ------------------------------------------------------------------------------MS-------------------------------------------------- 2
P.kyrbyi DL              ------------------------------------------------------------------------------MG-------------------------------------------------- 2
M.jakobiformis DL        ---------------------------------------------------------------------------------------------------------------------------------- 1
G.okellyi DL             ------------------------------------------------------------------------------MAL------------------------------------------------- 3
S.multiciliatum DL       ---------------------------------------------------------------------------------------------------------------------------------- 1
P.cosmopolitus WS DL     ---------------------------------------------------------------------------------------------------------------------------------- 1

                                140       150       160       170       180       190       200       210       220       230       240       250       260
                         ....|....|....|....|....|....|....|....|....|....|....|....|....|....|....|....|....|....|....|....|....|....|....|....|....|....|
A.godoyi D               ----------------------------------------------------------------------------------------------MFAV-------------------------------- 4
R.americana D            ----------------------------------------------------------------------------------------------MLPL-------------------------------- 4
M.jakobiformis D         ---------------------------------------------------------------------------------------------------------------------------------- 4
G.okellyi D              ----------------------------------------------------CSAEAEGSGQQSISDQLL-------RPFRI--SWLSIRHAVRLLTTSWSG---------------------------- 43
H.kukwesjijk D           ----------------------------------------------------------------------------------------------MLRL-------------------------------- 4
N.gruberi D              ----------------------------------------------------MENTLQPNSTNSLD-------------FKVLNNNVKERNNLSII---------------------------------- 31
N.fowleri D              ------------------------------------------------------------------------N-----IFSSSTSHSSHSSSWFMI---------------------------------- 21
Neo.damariscottae D      --------------------------------------------------------------------------------------------MKIL---------------------------------- 4
P.cosmopolitus AE1 D     -------------------------------------------------------------------------------------------KMKVMFNHHH----------------------------- 10
P.kyrbyi D               --------------------------------------------------------------------------------------------MNIV---------------------------------- 4
P.cosmopolitus WS D      ----------------------------------------PT---------------------------------------------NSHNAFGVLFFNY------------------------------ 28
E.coli                   ANFKDTDLKSFIETVGANLNK--------------TIIMGPG----------VQGKVSIRTMTPLNERQ------------------YYQLFLNLLEAQG----------------------YAV----- 107
V.cholerae               ASFKGTDIQEFINIVGRNLEK--------------TIIVDPS----------VRGKVDVRSFDTLNEEQ------------------YYSFFLSVLEVYG----------------------FAV----- 89
A.salmonicida            ASFKNADIEEFINTVGKNLSK--------------TIIIEPS----------VRGKINVRSYDLLNEEQ------------------YYQFFLSVLDVYG----------------------FAV----- 90
P.aeruginosa             INLKDADIREFIDQISEITGE--------------TFVVDPR----------VKGQVSVVSKAQLSLSE------------------VYQLFLSVMSTHG----------------------FTV----- 115
N.meningtitis            TEVRGNKVWIFINESDDTVSA-----PARP-----AVKAAPA-----APA--KQQAAAPSTKSAVSVSE---------PFTPAKQQAAAPFTESVVSVSAP---------------------FSPA---- 191
Myxococcus               VRADGNRVVISVDGAAQSVEAKRAEPPARTEGVTASVEVKPHSVSAAAPAKVVQAESAAVSKAALPENVVAAEADEREVSNPAQHITAMSFADDTLSIRADGDIARYEVLELADPPRLAVDLFGVGLATR 251
S.enterica               ---------------GDNRKGT-------------FYVSGPP----------VYVDMVVNAATMMDKQ-------------------------------------------------------------- 39
A.godoyi DL              ---------------------------------------------------------------------------------------------------------------------------------- 7
R.americana DL           ---------------------------------------------------------------------------------------------------------------------------------- 1
N.gruberi DL             ---------------------------------------------------------------------------------------------------------------------------------- 1
N.fowleri DL             ---------------------------------------------------------------------------------------------------------------------------------- 1
P.cosmopolitus AE1 DL    ---------------------------------------------------------------------------------------------------------------------------------- 1
Neo.damariscottae DL     ---------------------------------------------------------------------------------------------------------------------------------- 2
P.kyrbyi DL              ---------------------------------------------------------------------------------------------------------------------------------- 2
M.jakobiformis DL        ---------------------------------------------------------------------------------------------------------------------------------- 1
G.okellyi DL             ----------------------------------------PY---------------------------------------------------------------------------------------- 5
S.multiciliatum DL       ---------------------------------------------------------------------------------------------------------------------------------- 1
P.cosmopolitus WS DL     ---------------------------------------------------------------------------------------------------------------------------------- 1

                                270       280       290       300       310       320       330       340       350       360       370       380       390
                         ....|....|....|....|....|....|....|....|....|....|....|....|....|....|....|....|....|....|....|....|....|....|....|....|....|....|
A.godoyi D               -----------------------------------------LYSVL------------------------------------------------------------------------------------ 9
R.americana D            -----------------------------------------VQSLW------------------------------------------------------------------------------------ 9
M.jakobiformis D         -----------------------------------------LLGIW------------------------------------------------------------------------------------ 9
G.okellyi D              ---------------------------RGI-----------LVAVI------------------------------------------------------------------------------------ 51
H.kukwesjijk D           -----------------------------------------LLQVL------------------------------------------------------------------------------------ 9
N.gruberi D              -----------------------------------------IQRII------------------------------------------------------------------------------------ 36
N.fowleri D              -----------------------------------------LKDLI------------------------------------------------------------------------------------ 26
Neo.damariscottae D      -----------------------------------IEW---ILKII------------------------------------------------------------------------------------ 12
P.cosmopolitus AE1 D     -----------------------------------------LKKII------------------------------------------------------------------------------------ 15
P.kyrbyi D               -----------------------------------------FKKIW------------------------------------------------------------------------------------ 9
P.cosmopolitus WS D      -----------------------------------------FVNVFA----------------------------------------------------------------------------------- 34
E.coli                   -----------------------------------VPMENDVLKVV----------------------------KSSAAKVEPLPLVG----EGSDNYAGDEMVTKV----------------------- 147
V.cholerae               -----------------------------------VEMDNGVLKVI----------------------------KSKDAKTSAIPVL-----SGEERANGDEVITQV----------------------- 128
A.salmonicida            -----------------------------------VPMDNGVLKVV----------------------------RSKDAKTSAIPVV-----DETNPGIGDEMVTRV----------------------- 129
P.aeruginosa             -----------------------------------VAQGD-QARIV----------------------------PNAEAKTEA----------GGGQSAPDRLETRV----------------------- 148
N.meningtitis            ---------------------------------------------------KQQAAASAKQQ------------AAAPAKQQAAAPAKQTNIDFRKDGKNAGIIELAALGFAGQPDISQQHDHIIVTLKN 258
Myxococcus               APRVKSGALRDVRVGAHADKVRLVLDVRGTMPAYRVDRANRGLEVVLGRAVARKAAPAVETQAVVASVAEVEPLRQAPVKSEASPVVEVKDVRFEESSAGGRIVMKLSGTSAWKVDRPDPRSAVL-TLEN 380
S.enterica               ----NDG----------------------------IELGRQKIGVM----------------------------------------------RLNNTFVGDRTYN------------------------- 66
A.godoyi DL              ---------------------------------------------------------------------------------------------------------------------------------- 7
R.americana DL           ---------------------------------------------------------------------------------------------------------------------------------- 1
N.gruberi DL             ---------------------------------------------------------------------------------------------------------------------------------- 1
N.fowleri DL             ---------------------------------------------------------------------------------------------------------------------------------- 1
P.cosmopolitus AE1 DL    ---------------------------------------------------------------------------------------------------------------------------------- 1
Neo.damariscottae DL     ---------------------------------------------------------------------------------------------------------------------------------- 2
P.kyrbyi DL              ---------------------------------------------------------------------------------------------------------------------------------- 2
M.jakobiformis DL        ---------------------------------------------------------------------------------------------------------------------------------- 1
G.okellyi DL             ---------------------------------------------------------------------------------------------------------------------------------- 5
S.multiciliatum DL       ---------------------------------------------------------------------------------------------------------------------------------- 1
P.cosmopolitus WS DL     --------------------------------------------------------------------------MNHPFSSNSFP--------------------------------------------- 11

                                400       410       420       430       440       450       460       470       480       490       500       510       520
                         ....|....|....|....|....|....|....|....|....|....|....|....|....|....|....|....|....|....|....|....|....|....|....|....|....|....|
A.godoyi D               ----------------------------------------------------------------------------------------------------------------TWVLARGL---------- 17
R.americana D            ----------------------------------------------------------------------------------------------------------------RWLLERLSRL-------- 19
M.jakobiformis D         ----------------------------------------------------------------------------------------------------------------RFVLRIWPSLL------- 20
G.okellyi D              ---------------------------------------------------------------------------------------------------------------------------------- 51
H.kukwesjijk D           ----------------------------------------------------------------------------------------------------------------RFLIHGVSAIL------- 20
N.gruberi D              ----------------------------------------------------------------------------------------------------------------NWILSVLKGIFK------ 48
N.fowleri D              ----------------------------------------------------------------------------------------------------------------RFVWSILSQFIPSFWKRG 44
Neo.damariscottae D      ----------------------------------------------------------------------------------------------------------------RYILKIFNNLV------- 23
P.cosmopolitus AE1 D     ----------------------------------------------------------------------------------------------------------------KYVLLLFKKAPYTLYKLI 33
P.kyrbyi D               ----------------------------------------------------------------------------------------------------------------IFLIQIIPLTL------- 20
P.cosmopolitus WS D      ----------------------------------------------------------------------------------------------------------------CWITHSTTSFFSC---KR 49
E.coli                   -----------------VPVRNVSVRELAPI---LRQMIDSAGSGNVVNYDPS-----------------------------------------------NVIMLTGRASVVERLTEVIQRV-------- 202
V.cholerae               -----------------VAVKNVSVRELSPL---LRQLIDNAGAGNVVHYDPA-----------------------------------------------NIILITGRAAVVNRLAEIIRRV-------- 183
A.salmonicida            -----------------VPVRNVSVRELAPL---LRQLNDNAGGGNVVHYDPS-----------------------------------------------NVLLITGRAAVVNRLVEVVRRV-------- 184
P.aeruginosa             -----------------IQVQQSPVSELIPL---IRPLVPQYG--HLAAVPSA-----------------------------------------------NALIISDRSANIARIEDVIRQL-------- 201
N.meningtitis            HTLPTTLQRSLDVADFKTPVQKVTLKRLNND---TQLIITTAGNWELVNKSAAPGYFTFQVLPKKQNLESGGVNNAPKTFT------------------GRKISLDFQDVEIRTILQILAKESGMNIVAS 367
Myxococcus               ARLPKKFERSLDTSALDTPVKMISAFSVPGEGGKVRLVVAADGAIEEKVSQSAG-TLSWRLDVKGVKTEEVAVAQRTAGFTTEAPAYAAEGAPQQARYRGKRVSFEFKDIDIQNLLRVIAEISKKNIVVA 509
S.enterica               ----------------------------------LR----------------D-----------------------------------------------QKMVIPGIATAIERLLQ------------- 86
A.godoyi DL              ---------------------------------------------------------------------------------------------------------------------------------- 7
R.americana DL           ---------------------------------------------------------------------------------------------------------------------------------- 1
N.gruberi DL             ---------------------------------------------------------------------------------------------------------------------------------- 1
N.fowleri DL             ---------------------------------------------------------------------------------------------------------------------------------- 1
P.cosmopolitus AE1 DL    ---------------------------------------------------------------------------------------------------------------------------------- 1
Neo.damariscottae DL     ---------------------------------------------------------------------------------------------------------------------------------- 2
P.kyrbyi DL              ---------------------------------------------------------------------------------------------------------------------------------- 2
M.jakobiformis DL        ---------------------------------------------------------------------------------------------------------------------------------- 1
G.okellyi DL             ---------------------------------------------------------------------------------------------------------------------------------- 5
S.multiciliatum DL       ---------------------------------------------------------------------------------------------------------------------------------- 1
P.cosmopolitus WS DL     ---------------------------------------------------------------------------------------------------------------------------------- 11
```

```
                        ....|....|....|....|....|....|....|....|....|....|....|....|....|....|....|....|....|....|....|....|....|....|....|....|....|....|
A.godoyi D              RLRRGT-----------------------------------------VCINERSFL----------CPVAAHGVAG-------------QAVESAAGLQTV-----TCTALE------------------ 60
R.americana D           -------------------------------------------WRWLLWKSGRR------------PERAAE----------------SASTQSVRTVSAL-----EMQALEWSTS-------------- 59
M.jakobiformis D        -------SGSIAAL-----------------------------ASWYIWRINRR------------APVVQPSE--------------EMPLTKAGQISAL-----NMAANAFADP-------------- 69
G.okellyi D             -------------------------------------------AVVYFLLSRRRGH----------SPAALH-----------------SVQAVEGGVSLL-----NAMANEFTGN-------------- 92
H.kukwesjijk D          -------------------------------------------AQWRSIVSRLL------------FPVASHLSSSP------------SHLVCSSSVTAL-----NLQANAFFSQ-------------- 64
N.gruberi D             SNTSNNTNNNMFK-----------------------------------------------------SPISNSGDDDS-----------NNTASSSFECSQL-----SIYSDLFNQQ-------------- 95
N.fowleri D             RRVSNTTTITT-------------------------------------------------------TTTDFHNTSNTNST-----TSSSLTTKQQQQHSQL-----EINSYLFSND-------------- 95
Neo.damariscottae D     ------------------------------------------------------------------FPQQKQTETQE-----------NETIIQRFEFSEL-----EYLSNQYSNR-------------- 57
P.cosmopolitus AE1 D    NKVLDTCILKIFNIKTIQF----------------------------------------------IPHSLHPFVDQTMKHPKKNIMIPNIIVPSKRNITNL-----EMFSQAFRAPEQ------------ 100
P.kyrbyi D              -------IYALFSL-----I----------------------GSILPVLSSTRKNH----------FPTKPH-----------------NSSIQSKKVTGL-----EMAANQWNTG-------------- 70
P.cosmopolitus WS D     RGVAGAGNYSVWFI--------------------------FCFSFFTWLRERRKSHGVAVKRNGVSLPR-------------------DSTRSNERDISEL-----ELLSFQYQDE-------------- 115
E.coli                  -DHAGNRTEEVIPLDNASASEIARVLESLTKN---SGENQPATLKSQIVADERTNSVI------VSGDPATRDKMRRLIR------RLDSEMERSGNS--------QVFYLKYSKAEDLVDVLKQVSGT 307
V.cholerae              -DQAGDKEIEVVELNNASAAEMVRIVEALNKT--TDAQNTPEFLKPKFVADERTNSIL------ISGDPKVRERLKRLIK------QLDVEMAAKGNN--------RVVYLKYAKAEDLVEVLKGVSEN 289
A.salmonicida           -DKAGDQEVDIIKLKYASAGEMVRLVTNLNKDGNTQGGNTSLLLAPKVVADERTNSVV------VSGEPKARARIIQMVR------QLDRDLQSQGNT--------RVFYLKYGKAKDMVEVLKGVSTS 292
P.aeruginosa            -DQKGSHDYSVINLRYGWVMDAAEVLNNAMSRGQAKGA-----AGAQVIADARTNRLI------ILGPPQARAKLVQLAQ------SLDTPTARSANT--------RVIRLRHNDAKTLAETLGQISEG 304
N.meningtitis           DSVNGKMTLSLKDVPWDQALDLV--------------------MQARNLDMRQQGNIV------NIAPRDELLAKDK------------AFLQAEKDIADLGALYSQNFQLKYKNVEEFRSILR------ 453
Myxococcus              DDVSGKVTIRLRNVPWDQALDLV--------------------LRTKALGKEEFGNII------RIAPLKTLEEEARLRQ------ERKASLQRQEDLMV------NLIPVNYAVAADMAARVK------ 595
S.enterica              ------------------------------------GEEQPL------------GNIV------SSEPPAMPA---------------------FSANGE------KGKAANYAGGMSLQEALK------ 129
A.godoyi DL             ---------------------------------------------------------------------------------------------------------------------------------- 7
R.americana DL          ----------------------------------------------------------------------------------------------MAWDRRN-----TRNPSLY----------------- 14
N.gruberi DL            -----------------------------------------------MFNQQRNNY-------------------------------------MMGGGSGS----GQIQPS-Y--M-------------- 25
N.fowleri DL            ---------------------------------------------MSFQQQQRTNR-------------------------------------MMLPSSYQ-----DMSPS-Y--L-------------- 26
P.cosmopolitus AE1 DL   ----------------------------------------------MYQQQQPPGY-------------------------------------SYSMNPYN-----QTAPS-Y----------------- 24
Neo.damariscottae DL    -------------------------------------------FYSQYGQNEPTNYGS-------------------GYG-------NNYGGSSYSGGGYG-----GNAVG-YNRN-------------- 43
P.kyrbyi DL             -------------------------------------------RLSRYGYDDI-----------------------------------------------------ATLPS-Y----------------- 18
M.jakobiformis DL       -----------------------------------------------------------------------------------------------MDNVY------------------------------ 5
G.okellyi DL            -DRAG--------------------------------------AYGGYGGFGGPGYG---------GPGYGGAYGATGSG------MGYGAGTYGTNGYGA----GGYGPSAYNTG-------------- 63
S.multiciliatum DL      ------------------------------------------------------------------------------------------------------------MSS-Y----------------- 4
P.cosmopolitus WS DL    ----------------------------------------------------------------------------------------------------------PHQQSQYFPA-------------- 21

                             660       670       680       690       700       710       720       730       740       750       760       770       780
                        ....|....|....|....|....|....|....|....|....|....|....|....|....|....|....|....|....|....|....|....|....|....|....|....|....|....|
A.godoyi D              ---------------------------------------------------------------------------------------------------------------------------------- 60
R.americana D           ---------------------------------------------------------------------------------------------------------------------------------- 59
M.jakobiformis D        ---------------------------------------------------------------------------------------------------------------------------------- 69
G.okellyi D             ---------------------------------------------------------------------------------------------------------------------------------- 92
H.kukwesjijk D          ---------------------------------------------------------------------------------------------------------------------------------- 64
N.gruberi D             ---------------------------------------------------------------------------------------------------------------------------------- 95
N.fowleri D             ---------------------------------------------------------------------------------------------------------------------------------- 95
Neo.damariscottae D     ---------------------------------------------------------------------------------------------------------------------------------- 57
P.cosmopolitus AE1 D    ---------------------------------------------------------------------------------------------------------------------------------- 100
P.kyrbyi D              ---------------------------------------------------------------------------------------------------------------------------------- 70
P.cosmopolitus WS D     ---------------------------------------------------------------------------------------------------------------------------------- 115
E.coli                  LTAAKEEAEGTVGSGRE-----VVSIAASKHSNALIVTAPQDIMQSLQSVIEQLDIRRAQVHVEALIVEVAEGSNINFGVQWGSK-----DAG----LMQFANGTQIPIGTLGAAISAAKPQKGSTVISE 423
V.cholerae              LQAEKGTGQPTTSKRN------EVMIAAHADTNSLVLTAPQDIMNAMLEVIGQLDIRRAQVLIEALIVEMAEGDGINLGVQWGSL-----ESG---SVIQYGN-TGASIGNVMIGLEEAKDTTQTKAVYD 404
A.salmonicida           IEADKKGGGATAGGGNASIGGGKLAISADETTNALVITAQPDVMAELEQVVAKLDIRRAQVLVEAIIVEIADGDGLNLGVQWANT-----NGG----GTQFTN-AGPGIGSVAIAAKDYKDN-------- 404
P.aeruginosa            MKNNGGQGGEQTGGGRPS----NILIRADESTNALVLLADPDTVNALEDIVRQLDVPRAQVLVEAAIVEISGDIQDAVGVQWAIN-----KGGMGGTKTNFAN-TGLSIGTLLQSLESNKAP-------- 416
N.meningtitis           ------LDNADTTGNRNTLVSGRGSVLIDPATNTLIVTDTRSVIEKFRKLIDELDVPAQQVMIEARIVEAADGFSRDLGVKFG-------ATGKK----KLKNDTSAFGWGVNSGFGGDDKW-------- 558
Myxococcus              -----------------DVLSERGSVTVDQRTNVLIVKDVRSNTERARSLVRSLDTQTPQVLIESRIVEANTSFSRSLGVQWGGQARAGQATGNS-TGLIFPNNLAVTGGVTGTGAGLPDN--------- 698
S.enterica              ----------------QNAAAGNIKIVAYPDTNSLLVKGTAEQVHFIEMLVKALDVAKRHVELSLWIVDLNKSDLERLGTSWSGSITIGDKLG------------------------------------- 206
A.godoyi DL             ---------------------------------------------------------------------------------------------------------------------------------- 7
R.americana DL          -----------------------------------------------------------------------------------------------------------DTR-------------------- 17
N.gruberi DL            ----------------------------------------------------------------------------------------------------YNDMISPSSS-------------------- 35
N.fowleri DL            ----------------------------------------------------------------------------------------------------YNDAPYGGYG-------------------- 36
P.cosmopolitus AE1 DL   ---------------------------------------------------------------------------------------------------------------------------------- 24
Neo.damariscottae DL    ----------------------------------------------------------------------------------------------------YMN--------------------------- 46
P.kyrbyi DL             ---------------------------------------------------------------------------------------------------------------------------------- 18
M.jakobiformis DL       ---------------------------------------------------------------------------------------------------------------------------------- 5
G.okellyi DL            ----------------------------------------------------------------------------------------------------YGSGGYGTTGGYGAGGYGPSGY-------- 85
S.multiciliatum DL      ---------------------------------------------------------------------------------------------------------------------------------- 4
P.cosmopolitus WS DL    ---------------------------------------------------------------------------------------------------------------------------------- 21

                             790       800       810       820       830       840       850       860       870       880       890       900       910
                        ....|....|....|....|....|....|....|....|....|....|....|....|....|....|....|....|....|....|....|....|....|....|....|....|....|....|
A.godoyi D              ---------------------------------------------------------------------------------------------------------------------------------- 60
R.americana D           -------------------------------------------------AVATVVDSDII-TLEEPE---------------------------------------------------EEPESSTAVALA 88
M.jakobiformis D        ---------------------------------------------------------------------------------------------------------------------------------- 69
G.okellyi D             --------------------------------------------------------------------------------------------------------------------AGTANLPALRLNLG 106
H.kukwesjijk D          ----------------------------------------------------QTAHHGHQQQLQPQT--AGLFSQLFHDSC---------------TSAQDLTTSREL---------------------- 103
N.gruberi D             ---------------------------------------------------VNIDDNHQIQSIKPNH----------------------------------INHDDKP---------QLMNQIDI----- 126
N.fowleri D             -------------------------------------------------HHPDIISSPSLFNTQPNV--VPLFSNHHQNHTTSVVGDDPHQLLTTFLIPMSNNVEFHK---------CDASSS----YLL 161
Neo.damariscottae D     ------------------------------------------------------------------------------------------------------------------------NENKSQLKMI 67
P.cosmopolitus AE1 D    ----------------------------------------------DHSQTADRFYQEVYETLLNPS------------------HKIKSSTDTTFKFPVDKNISLKD---------SLATQHPTIGEMS 157
P.kyrbyi D              ----------------------------------------------------SKTKQTINEDLENPK--LSSTNN---------------------THNNDLNLSSHP---------NEEVNV----QII 112
P.cosmopolitus WS D     ----------------------------------------------------NCVGEPHILTDNGNN--ATVVRNANFITAFVGVRDKVHTQGNSKVLMNHTNASNDPTSFNG----------------- 174
E.coli                  NGATTINPDTN---GDLSTLAQLLSGFSGTAVGVVKGDWMALVQAVKNDSSSNVLSTPSITTLDNQE--AFFMVG----------QDVPVLTGS--TVGSNNSNPFNT---------VERKKVGIMLKVT 527
V.cholerae              TNNNFLRNETTTTKGDYTKLASALSSIQGAAVSIAMGDWTALINAVSNDSSSNILSSPSITVMDNGE--ASFIVG----------EEVPVITGS--TAGSNNDNPFQT---------VDRKEVGIKLKVV 511
A.salmonicida           --------------GTTTGIAKLAESFNGMAAGFYQGNWAMLVTALSTNTKSDILSTPSIVTMDNKE--ASFNVG----------QEVPVQTGK--QNSTSGDTTFST---------IERKTVGTKLVVT 497
P.aeruginosa            -----------------------ESIPDGAIVGIGSSSFGALVTALSANTKSNLLSTPSLLTLDNQK--AEILVG----------QNVPFQTGSYTTNSEGSSNPFTT---------VERKDIGVSLKVT 502
N.meningtitis           GAETKINLPIT-----------AAANSISLVRAISSGALNLELSASESLSKTKTLANPRVLTQNRKE--AKIESG----------YEIPFTVTSIANGGSSTNTELKK--------------AVLGLTVT 651
Myxococcus              -PNFAVNLPTG------TGQGVGGAMGFTFGSAGGALQLNLRLSAAENEGSVKTISAPKVTTLDNNT--ARISQG----------VSIPFSQ----TSAQGVNTTFVE--------------ARLSLEVT 791
S.enterica              -----------------------VSLNQSSISTLDGSRFIAAVNALEEKKQATVVSRPVLLTQENVP--AIF----------------------------DNNRTFYTKLIGERNVALEHVTYGTMIRVL 283
A.godoyi DL             ---------------------------------------------VAERRSVESLLRDPLYNRSGRRWVSPYEDGRP-SLRRLDYAD----------------LPFE---------------------DR 54
R.americana DL          ---------------------------------------------ALMDPQAPYVAQ---------Q--RDRYTNMF-DPRSRYFED----------------FPYR---------------------ER 53
N.gruberi DL            ---------------------------------------------SMMRNNSPYMNSDPYYNPMSNR--RGYYDNYMIGNEPLYHED----------------KLFDEDLLMY---------------DR 87
N.fowleri DL            ---------------------------------------------GLSTPSPPYNYSDPYYNSMQNR--RGYYDNYMIGNEPLYHED----------------KLFDEDLLMY---------------DR 88
P.cosmopolitus AE1 DL   ----------------------------------------------FNRPQTRYAYQA--QPEYGGV---PYGSQYY-QQQPLYMED----------------RLFDDETAHY---------------AR 71
Neo.damariscottae DL    ---------------------------------------------RRNMPTSSYRRSP--YNQEGNM--SSYYDEYN-NNSPQYYED----------------RIYDDEAREY---------------ER 95
P.kyrbyi DL             ----------------------------------------------FRQP-TSYDY-----------------NRGR-NNTGRYTR------------------------------------------TR 41
M.jakobiformis DL       -------------------------------------------------------------PPPYFG--GDYYN----GFQPQWY--------------------------------------------- 23
G.okellyi DL            VPRYGGAGPMP----------------------------------GFVGP-QMYGPGPGQMPPLAGA--PGFFP----GFAPGW---------------------------------------------- 128
S.multiciliatum DL      ------------------------------------------------------------RDPPSHH--PGY------SNHPRYSSAGP-----------AAATPVSDSA------------------EW 37
P.cosmopolitus WS DL    ------------------------------------------------NSSNPYSWQSQYQNLPSHASPYSFYGGNAAANNYASPQD----------------QIFSNDEMQYI--------------NR 73

                             920       930       940       950       960       970       980       990      1000      1010      1020      1030      1040
                        ....|....|....|....|....|....|....|....|....|....|....|....|....|....|....|....|....|....|....|....|....|....|....|....|....|
A.godoyi D              ----------------------AASVAGVRRSRWQGMRIKKHFVTQTAVRVRP-SDSQFLGSFALSTELRTVQSGP-------LALIP----FL--KHWF--SRTEFVLVKSAILVFLTLHVEPYRPVLA 152
R.americana D           P------------------------QPTMGASRYSAIRIKKNFTLQTAMLLKE-NVPSYAGSFALATELRRVRRSP-------LSFIP----FL--ERFF--SRHEFMLVKSAILVFITVRTEAAGRPQV 178
M.jakobiformis D        ----------------------AVSMLASRRSRLVSMRIRKNFALQTSVMLKE-NQATYVGAFAISNELKTIRRGP-------LSYIP----LI--GDLF--RRDEFMRVKSAILVFVTCRAERTGSGPR 161
G.okellyi D             P--------------------------LALRSRLTSMRLRKNFTLQTSLMLKE-GQTTYAGSFALASELKTLKQGP-------LSFLP----VV--GQLF--SRDEFMLIKSAVLVFVTFHKERSQAAAA 194
H.kukwesjijk D          P---LVNAADCA---------FGDTFGGGGASRIESLRARKNFSLQTTMMLKE-NQTTYVGALAVGTELRTVKRSP-------LSYIP----FL--SRLF--TREEFMRVKSAILVFITARLQKLQPHVL 205
N.gruberi D             ----QFNKCI-----------SGTALPSMNSSKYEAMRIKKHFTTSTNLLIKEFDKPHFVGSFSNSTELKTIRKSP-------LSFIP----FL--SDLF--TRNEYMIVKSAILVFITASLKND----- 221
N.fowleri D             P----------------------SSLSPSSSSKYQAMRIKKHFTASTNLLIRELNTPHFVGSFANSTELKAIRKSP-------LSFLPF--QFL--SQLF--SRNEYMLVKSAILVFITASLDQQQQHDH 256
Neo.damariscottae D     P------------------------VSDELKSRWESMRLKKHFTFQTNVLLKP-NQPTYVGAFSSSTELQTVKRSP-------FSYLP----FL--RDLF--SRKEFILIKSAILVFITAKYEDKKTNQL 157
P.cosmopolitus AE1 D    PYGGLFN--------------PFTGFSELVKSSWNAMRLKKHFTVQTNMLLTP-NEPTFVGAFAVSQQLKTIKRSP-------LSFLP----FL--KWIF--ARTEYMLIKSAVLVFITARFSES----- 252
P.kyrbyi D              P-----------------------SNSNNGRSRWDAMRLKKHFAVQTTVMLKE-NEPSFAGAFAASTELRTVKKSP-------FSFLP----FKWGKALF--SRTEFMLVKSAILVFIKCIPQRAFNVTE 205
P.cosmopolitus WS D     PVGDFFSDPA-----------NPFSPVAYGKSSIQAIRLRKHFTCQTQLMVQP-NEPTYVGAFAMSTDIRTVRQSP-------LRALW--------GWLW--DRTEYLKIKSALLVFVTCRILKPDET-- 273
E.coli                  P---QINEGNAVQMVIEQEV-SKV----EGQTSLD--VVFGERKLKTTVLAND-GELIVLGGLMDDQAGESVAKVP------LLGDIP----LI--GNLF--KSTADKKEKRNLMVFIRPTILRDGMAAD 632
V.cholerae              P---QINEGNSVQLNIEQEV-SNV---LGANGAVD--VRFAKRQLNTSVMVQD-GQMLVLGGLIDERALESESKVP------LLGDIP----LL--GQLF--RSTSSQVEKKNLMVFIKPTIIRDGVTAD 617
A.salmonicida           P---QINEGDSVLLTIEQEV-SSVGKQASGTEGLG--PTFDTRTVKNAVLVKS-GETVVLGGLMDEQTKEEVSKVP------LLGDIP----VL--GYLF--RSTSNTTSKRNLMVFIRPTILRDANVYS 606
P.aeruginosa            P---HINDGAALRLEIEQEI-SALLPNAQQRNNTD--LITSKRSIKSTILAEN-GQVIVIGGLIQDDVSQAESKVP------LLGDIP----LL--GRLF--RSTKDTHTKRNLMVFLRPTVVRDSAGLA 611
N.meningtitis           P---NITPDGQIIMTVK----ITKDSPAQCASGNQTILCISTKNLNTQAMVEN-GGTLIVGGIYEEDNGNTLTKVP------LLGDIP----VI--GNLF--KTRGKKTDRRELLIFITPRIMGTAG--- 756
Myxococcus              P---HITQDGSVLMSIN----ASNNQPDPSSTGANGQPSIQRKEANTQVLVKD-GDTTVIGGIYVRRGATQVNSVP------FLSRIP----VL--GLLF--KNNSETDTRQELLIFITPRILNRQTIAQ 899
S.enterica              P---RFSADGQIEMSLDIEDGNDKTPQSDTTTSVDALPEVGRTLISTIARVPH-GKSLLVGGYTRDANTDTVQSIP------FLGKLP----LI--GSLF--RYSSKNKSNVVRVFMIEPKEIVDPLTPD 395
A.godoyi DL             P--------------------TYVPSQLLANPDFGNFRIRRSFT--TELWLQD-GETVAVLGDGKEREFRT-HYSPMYRRPFFSGWIGRLLQFLALDRLFGLHIDSHKKSRTRSVLLVTAKLLPLNEQRT 160
R.americana DL          A--------------------TYSRGSALGNPDFGNFRFRKGYS--AELTLRD-GETAMLSGDSRSKSYRT-HYSPEYRSSFFYSWLGKLMRLLWLDRLFGLHVDTVKHVRTRSYVFVTARILPDAATAT 159
N.gruberi DL            P--------------------TYRPSRMIANPDFGNFNYKKGYT--AEVWLND-GETVVLSGDSRKKTYRT-HYSPEYLRSFFFGWVGRLIRYLWMDRLFGLHVDTVKEVKTKNYIFVTAQLLSE----- 188
N.fowleri DL            P--------------------TYRPSRMIANPDFGNFNYKKGFT--AEVWLND-GETVVLNGDSRKKTYRT-HYSPEYLRSFFFGWVGRLIRYLWLDRLFGLHVDTVKEVKTRNYIFVTTQIIPE----- 189
P.cosmopolitus AE1 DL   P--------------------SYRRSAMLGNPDFGNYHFKKGYA--TEVWLND-GETVVLAADTRKKKYRT-HYSPQFLRSFFFGWIGRLLRFLMLDRLFGLHVDTTKTVTTRNYIFVTAHVLPS----- 172
Neo.damariscottae DL    P--------------------TYRRSKTIANPDFGNFHFKKGYT--AEVWLND-GETVVLSSDSRKKSYRT-HYSPEYMRSFFFGWIGRLLRFFYLDRLFGLHVDTVKNVTIKNFIFVTAHSLPI----- 196
P.kyrbyi DL             P--------------------TYVRSNLAASPDFGNFGFRKGYT--ASVWMND-GETVVLSADRKQRTYRT-HYSPEFLRSFFYGWLGRLLRFFMLDRLFGFHMDTVKTVNKRSFVFVTAHVLDT----- 142
M.jakobiformis DL       ----------------------YRPSRLLANPDFGNFRYKKGFT--AEVTLED-GETVVLAADSRVKTYRT-HYSP--FNRYSFSWLDRILRFLWLDRLLGLNIDSVKRIRTKSFIFVTAHLLPH----- 120
G.okellyi DL            ---------------------AYRPASLAANPDFGTYRYKKGYT--AEVTLQD-GETVVLAADSRTKSYRT-QYSP--FNRYSFSWLDRLLRFLWLDRLLGLHVDSVKHVRTKSFICVTAHLLPS----- 226
S.multiciliatum DL      P--------------------QPRTSVLQGSPDFGNLHFRKGFT--SEVMLND-GETAILSAHSRNRSYRL-RYSPIF--SFVYGWIRRFLAFFWLDRLLGPQVDSVKTVKTKSFVMVSAHILPAV---- 137
P.cosmopolitus WS DL    P--------------------TMHRSSLLASPSFPNLSLKHAIS--TNVTLES-GRTVALQSQEREKVFRG-RYWSGWKWYRMSGWLKVLVYAF--RWLIGLNIDTVKKVKVSNMIYLTGYVLPT----- 172
```

```
                              1050      1060      1070      1080      1090      1100      1110
                          ....|....|....|....|....|....|....|....|....|....|....|....|....|....|....|...
A.godoyi D                LNDIAEQ-----IED--------------------------------------------------------------- 162
R.americana D             TQS--------------------------------------------------------------------------- 181
M.jakobiformis D          LTQQQQ---PMLQTDARY------------------------------------------------------------ 177
G.okellyi D               PVQTAL------------------------------------------------------------------------ 200
H.kukwesjijk D            AQ---------------------------------------------------------------------------- 207
N.gruberi D               ------------------------------------------------------------------------------ 221
N.fowleri D               NNMNNTSIGSSSISGGHGSGG--------------------------------------------------------- 277
Neo.damariscottae D       KGNVIKK----------------------------------------------------------------------- 164
P.cosmopolitus AE1 D      ------------------------------------------------------------------------------ 252
P.kyrbyi D                TTM-------PAIENEPQYFGEQ-----------------Y------------------------------------- 222
P.cosmopolitus WS D       ------------------------------------------------------------------------------ 273
E.coli                    GVSQRKY---NYMRAEQIYRDEQ-GL-SLMPHTAQPILP--AQNQA-LPPEVRAFLNAGRTR---------------- 686
V.cholerae                GITQRKY---NYIRAEQLFRAEK-GL-RLLDDASVPVLPKFGDDRR-HSPEIQAFIEQMEAKQ--------------- 674
A.salmonicida             GISSNKY---TLFRAQQLEAAAQKGY-ATSPD--RQVLPEYGQDVV-QSPEIQKQIELMKARQQATADGAQPFVQGNN 677
P.aeruginosa              ALSGKKY---SDIRVIDGTRGPE-GRPSILPTNANQLFDGQAVDLRELMTE--------------------------- 658
N.meningtitis             ----------NSLRY--------------------------------------------------------------- 761
Myxococcus                TL---------------------------------------------------------------------------- 901
S.enterica                ASESVN----NILKQSGAWSGDD-----------------------KLQKWVRVYLDRGQEAI--------------K 432
A.godoyi DL               ATDKSN----DDVSVSES------------------------------------------------------------ 174
R.americana DL            AAPSQA----SEVS---------------------------------------------------------------- 169
N.gruberi DL              ------------------------------------------------------------------------------ 188
N.fowleri DL              ------------------------------------------------------------------------------ 189
P.cosmopolitus AE1 DL     ------------------------------------------------------------------------------ 172
Neo.damariscottae DL      ------------------------------------------------------------------------------ 196
P.kyrbyi DL               ------------------------------------------------------------------------------ 142
M.jakobiformis DL         ------------------------------------------------------------------------------ 120
G.okellyi DL              ------------------------------------------------------------------------------ 226
S.multiciliatum DL        ------------------------------------------------------------------------------ 137
P.cosmopolitus WS DL      ------------------------------------------------------------------------------ 172
```

### (B) GspDN1

```
AgoGspDN1       ---------------------------------------------------------------------------------------------------------------------------------- 1
RamGspDN1       ---------------------------------------------------------------------------------------------------------------------------------- 1
NgrGspDN1       ---------------------------------------------------------------------------------------------------------------------------------- 1
NfoGspDN1       ---------------------------------------------------------------------------------------------------------------------------------- 1
NdaGspDN1       ---------------------------------------------------------------------------------------------------------------------------------- 1
PcoAEGspDN1     ---------------------------------------------------------------------------------------------------------------------------------- 1
PkiGspDN1       ---------------------------------------------------------------------------------------------------------------------------------- 1
GokGspDN1       ---------------------------------------------------------------------------------------------------------------------------------- 1
SpiGspDN1       ---------------------------------------------------------------------------------------------------------------------------------- 1
E.coli          MFWR--------DMT----------------------------------------------LSIWRKKTTGLKTKKR-LLPLMLAA-------------------------ALCSSPVWAEEATFANFKDT 52
V.cholerae      -------------------------------------------------------------MKYWLKKSSWL-----------LAG-----------------------SLLSTPL-AMANEFSASFKGT 34
A.salmonicida   ---------------------------------------------------------------MINKGKSWR-------LAT-VAA-----------------------ALMMAGS-AWATEYSASFKNA 35
P.aeruginosa    MSQP--------LLR----------------------------------------------ALFAPSSRSYVPAVLL-SLALGIQAAH--------------AENSGGNAFVPAGN-QQEAHWTINLKDA 60
N.meningtitis   MNTK--------LTKIISGLFVATAAFQTASAGNITDIKVSSLPNKQKIVKVSFDKEIVNPTGFVTSSPARIALDF---EQTGISMDQQVLEYADPLLSKISAAQNSSRARLVLNL-NKPGQYNTEVRGN 118
Myxococcus      MLEESAVTRGKWMLAAAWAVVLVGARVHGAELNTLRGLDVSRTGSGAQVVVTGTRPPTF--TVFRLSGPERLVVDLSSADATGIKGHHEGSGPVSGVVASQFSDQRASVGRVLLAL-DKASQYDVRADGN 127
S.enterica      LYNKNYPLRGD----------------------------------------------------------------------------------------------------------NRKGTF------- 17

                       140       150       160       170       180       190       200       210       220       230       240       250       260
                ....|....|....|....|....|....|....|....|....|....|....|....|....|....|....|....|....|....|....|....|....|....|....|....|....|....|
AgoGspDN1       ---------------------------------------------------------------------------------------------------------------------------------- 1
RamGspDN1       ---------------------------------------------------------------------------------------------------------------------------------- 1
NgrGspDN1       ---------------------------------------------------------------------------------------------------------------------------------- 1
NfoGspDN1       ---------------------------------------------------------------------------------------------------------------------------------- 1
NdaGspDN1       ---------------------------------------------------------------------------------------------------------------------------------- 1
PcoAEGspDN1     ---------------------------------------------------------------------------------------------------------------------------------- 1
PkiGspDN1       ---------------------------------------------------------------------------------------------------------------------------------- 1
GokGspDN1       ---------------------------------------------------------------------------------------------------------------------------------- 1
SpiGspDN1       ---------------------------------------------------------------------------------------------------------------------------------- 1
E.coli          DLKSFIETVGANLNK--------------TIIMGPG----------VQGKVSIRTMTPLNER------------------QYYQLFLNLLEAQG----------------------YAV----------- 107
V.cholerae      DIQEFINIVGRNLEK--------------TIIVDPS----------VRGKVDVRSFDTLNEE------------------QYYSFFLSVLEVYG----------------------FAV----------- 89
A.salmonicida   DIEEFINTVGKNLSK--------------TIIIEPS----------VRGKINVRSYDLLNEE------------------QYYQFFLSVLDVYG----------------------FAV----------- 90
P.aeruginosa    DIREFIDQISEITGE--------------TFVVDPR----------VKGQVSVVSKAQLSLS------------------EVYQLFLSVMSTHG----------------------FTV----------- 115
N.meningtitis   KVWIFINESDDTVSA-----PAR------PAVKAAPAA------PAKQQAAAPSTKSAVSVSE---------PFTPAKQQAAAPFTESVVSVSAP---------------------FSP----------- 190
Myxococcus      RVVISVDGAAQSVEAKRAEPPARTEGVTASVEVKPHSVSAAAPAKVVQAESAAVSKAALPENVVAAEADEREVSNPAQHITAMSFADDTLSIRADGDIARYEVLELADPPRLAVDLFGVGLATRAPRVKS 257
S.enterica      ---------------------------------------------------------------------------------------------------------------------------------- 17

                       270       280       290       300       310       320       330       340       350       360       370       380       390
                ....|....|....|....|....|....|....|....|....|....|....|....|....|....|....|....|....|....|....|....|....|....|....|....|....|....|
AgoGspDN1       ---------------------------------------------------------------------------------------------------------------------------------- 1
RamGspDN1       ---------------------------------------------------------------------------------------------------------------------------------- 1
NgrGspDN1       ---------------------------------------------------------------------------------------------------------------------------------- 1
NfoGspDN1       ---------------------------------------------------------------------------------------------------------------------------------- 1
NdaGspDN1       ---------------------------------------------------------------------------------------------------------------------------------- 1
PcoAEGspDN1     ---------------------------------------------------------------------------------------------------------------------------------- 1
PkiGspDN1       ---------------------------------------------------------------------------------------------------------------------------------- 1
GokGspDN1       ---------------------------------------------------------------------------------------------------------------------------------- 1
SpiGspDN1       ---------------------------------------------------------------------------------------------------------------------------------- 1
E.coli          -----------------------------VPMENDVLKVV----------------------------KSSAAKVEPLPLVG----EGSDNYAGDEMVTKV----------------------------- 147
V.cholerae      -----------------------------VEMDNGVLKVI----------------------------KSKDAKTSAIPVL-----SGEERANGDEVITQV----------------------------- 128
A.salmonicida   -----------------------------VPMDNGVLKVV----------------------------RSKDAKTSAIPVV-----DETNPGIGDEMVTRV----------------------------- 129
P.aeruginosa    -----------------------------VAQGDQA-RIV----------------------------PNAEAKTEA----------GGGQSAPDRLETRV----------------------------- 148
N.meningtitis   ----------------------------------------A----KQQAAASAKQQ------------AAAPAKQQAAAPAKQTNIDFRKDGKNAGIIELAALGFAGQPDISQQHDHIIVTLKNHTLPTT 264
Myxococcus      GALRDVRVGAHADKVRLVLDVRGTMPAYRVDRANRGLEVVLGRAVARKAAPAVETQAVVASVAEVEPLRQAPVKSEASPVVEVKDVRFEESSAGGRIVMKLSGTSAWKVDRPDPRSAVL-TLENARLPKK 386
S.enterica      ---------------------------------------------------------------------------------------------------------------------------------- 17

                       400       410       420       430       440       450       460       470       480       490       500       510       520
                ....|....|....|....|....|....|....|....|....|....|....|....|....|....|....|....|....|....|....|....|....|....|....|....|....|....|
AgoGspDN1       ---------------------------------------------------------------------------------------------------------------------------------- 1
RamGspDN1       ---------------------------------------------------------------------------------------------------------------------------------- 1
NgrGspDN1       ---------------------------------------------------------------------------------------------------------------------------------- 1
NfoGspDN1       ---------------------------------------------------------------------------------------------------------------------------------- 1
NdaGspDN1       ---------------------------------------------------------------------------------------------------------------------------------- 1
PcoAEGspDN1     ---------------------------------------------------------------------------------------------------------------------------------- 1
PkiGspDN1       ---------------------------------------------------------------------------------------------------------------------------------- 1
GokGspDN1       ---------------------------------------------------------------------------------------------------------------------------------- 1
SpiGspDN1       ---------------------------------------------------------------------------------------------------------------------------------- 1
E.coli          -----------VPVRNVSVRELAPI---LRQMIDSAGSGN-VVNYDP-----------------------------------------------SNVIMLTGRASVVERLTEVIQRV---------DHAG 206
V.cholerae      -----------VAVKNVSVRELSPL---LRQLIDNAGAGN-VVHYDP-----------------------------------------------ANIILITGRAAVVNRLAEIIRRV---------DQAG 187
A.salmonicida   -----------VPVRNVSVRELAPL---LRQLNDNAGGGN-VVHYDP-----------------------------------------------SNVLLITGRAAVVNRLVEVVRRV---------DKAG 188
P.aeruginosa    -----------IQVQQSPVSELIPL---IRPLVPQ--YGH-LAAVPS-----------------------------------------------ANALIISDRSANIARIEDVIRQL---------DQKG 205
N.meningtitis   LQRSLDVADFKTPVQKVTLKRLNND---TQLIITTAGNWE-LVNKSAAPGYFTFQVLPKKQNLESGGVNNAPKTF------------------TGRKISLDFQDVEIRTILQILAKESGMNIVASDSVNG 372
Myxococcus      FERSLDTSALDTPVKMISAFSVPGEGGKVRLVVAADGAIEEKVSQSA--GTLSWRLDVKGVKTEEVAVAQRTAGFTTEAPAYAAEGAPQQARYRGKRVSFEFKDIDIQNLLRVIAEISKKNIVVADDVSG 514
S.enterica      -----------------------------------------YVSGPP-----------------------------------------------VYVDMVVNAATMMDKQNDGIE-------------LG 46

                       530       540       550       560       570       580       590       600       610       620       630       640       650
                ....|....|....|....|....|....|....|....|....|....|....|....|....|....|....|....|....|....|....|....|....|....|....|....|....|....|
AgoGspDN1       -------------------------------------------------------------------------------------------MSNGPHYLFLQHSRLTDEVQKYVSSFVQYIASIVEPD-- 37
RamGspDN1       ---------------------------------------M-STPPAAAAAPPSVSPPPPLVTSTEAPPPDEDSAPASAPPLRRPVGDAAEGDTREPYYIFLQNTTLTRAVQRYIAAFVQHVAGVLEPEPP 90
NgrGspDN1       ----------------------------------------------MSKQQTTTNTLPT-----TST--------------------SEDDNIREPFYIYLQNSELNTNLQKYISAFVQHIASILEQD-- 57
NfoGspDN1       ----------------------------------------------MSTNQEDLTQRP------NKT--------------------DHDDDIREPFYIYLQNSELNTNLQKYISAFVQHIASILEQD-- 56
NdaGspDN1       --------------------------------------------------------------MNE----------------------GQQKQEKEPFYIYLENSQLNEKLQKYISDFIQHIASITEKE-- 44
PcoAEGspDN1     ----------------------------------------------------MTDSSPNPTQQEENE---------------------SKTQSREPYYIYLQNSELTEALQKYIAAFVQHIASILEKD-- 55
PkiGspDN1       -----------------------------------------MNSSSSSSESQFENNIPPPPNVNENALKD----------------IQLDNESREPFYIYLQNSILTNDLQRYIGAFVQHIASILEKN-- 71
GokGspDN1       -----------------------------------------------------------------------------------------MASAPEPYYIYLTNSVLDGNLQKYISAYVEHIAKTLEGP-- 39
SpiGspDN1       --------------------------------------------------------MLMDPLQDRSDQPQP----------------LGSEQPSAEPFCVHLSNAVMDPQLQQFISSFVDYIASVLEPP- 57
E.coli          NRTEEVIPLDNASASEIARVLESLTKNS---GENQPATLKSQIVADERTNSVIVSGDPATRDKMRRLIRRLDSEMERSGNS----------QVFYLKYSKAEDLVDVLKQVSGTLTAAKEEAEGTVGSGR 323
V.cholerae      DKEIEVVELNNASAAEMVRIVEALNKTTD--AQNTPEFLKPKFVADERTNSILISGDPKVRERLKRLIKQLDVEMAAKGNN----------RVVYLKYAKAEDLVEVLKGVSENLQAEKGTGQPTT-SKR 304
A.salmonicida   DQEVDIIKLKYASAGEMVRLVTNLNKDGNTQGGNTSLLLAPKVVADERTNSVVVSGEPKARARIIQMVRQLDRDLQSQGNT----------RVFYLKYGKAKDMVEVLKGVSTSIEADKKGGGATAGGGN 308
P.aeruginosa    SHDYSVINLRYGWVMDAAEVLNNAMSRGQAKGA-----AGAQVIADARTNRLIILGPPQARAKLVQLAQSLDTPTARSANT----------RVIRLRHNDAKTLAETLGQISEGMKNNGGQGGEQTGGGR 320
N.meningtitis   KMTLSLKDVPWDQALDLV--------------------MQARNLDMRQQGNIVNIAP------RDELLAKDKAFLQAEKDIA-D-LGALYSQNFQLKYKNVEEFRSILRLDNADTTGNRN----TLVSGR 470
Myxococcus      KVTIRLRNVPWDQALDLV--------------------LRTKALGKEEFGNIIRIAPLKTLEEEARLRQERKASLRQREDLMV--------NLIPVNYAVAADMAARVK---------------DVLSER 601
S.enterica      RQKIGVMRLNNTFVGDRT--------------------------YNLRDQKMVIPGIATAIERLLQGEEQPLGNIVSSEPPAMPAFSANGEKGKAANYAGGMSLQEALKQ--------------NAAAGN 136

                       660       670       680       690       700       710       720       730       740       750       760       770       780
                ....|....|....|....|....|....|....|....|....|....|....|....|....|....|....|....|....|....|....|....|....|....|....|....|....|....|
AgoGspDN1       ---TEPSRRIISFLPDRNGVVLVGFQDSTVRVIHRFLEKMDEEKRTLKFTVHLVLAESHRMTSNFSF--------------------------------------------------------------- 101
RamGspDN1       GT--PSERRIVSFLPERNGVLFLGFRRSTVRVIYKFLTRMDTEKRNLHFEVNLVLVESHRMTSQFSF--------------------------------------------------------------- 155
NgrGspDN1       ---APNIRRIVSFLPERNGILLFGFRRSTIKVLYSFLKSLDKKKNNINFEINLVLVESHKMTNNFNF--------------------------------------------------------------- 121
NfoGspDN1       ---APNVRRIVSFLPERNGVLLFGFRRSTIKVLYSFLKSLDQKKHNINFEINLVLVESHKMTHDFNF--------------------------------------------------------------- 120
NdaGspDN1       ---ETTTKRVISFLPERNGILLLGFKSSTVKVIYHFLKKLDEEKRNINFEVNLVLVESHKMTSGFKF--------------------------------------------------------------- 108
PcoAEGspDN1     ---APSVKRIVSFLPERNGILLFGFQSTTIETIYRFLKGLDEKKYNIDFEVNLVLVESHKMTSGFKF--------------------------------------------------------------- 119
PkiGspDN1       ---DKNSRRVVSFLPERNGILLFGFKVSTVKVIYKFLRRLDEEKHNLHFEVNLVLVESHKMTGGFQF--------------------------------------------------------------- 135
GokGspDN1       ----AHANRVVTYIPERNGILLFGFQPETIRVIYRLLQKLDEETFNLRFKVDLVLVESHRMTSNFEF--------------------------------------------------------------- 102
SpiGspDN1       ---ESAEMRHVAFMPERGGILLFGFRPSTVQVIYRFLAKIDEERHNIRFEINLVYVESHKMTNAFEF--------------------------------------------------------------- 121
E.coli          -EVV-----SIAASKHSNALIVTA-PQDIMQSLQSVIEQLDIRRAQVHVEALIVEVAEGSNINFGVQWGSK-----DAG----LMQFANGTQIPIGTLGAAISAAKPQKGSTVISENGATTINPDTN--- 434
V.cholerae      -NEV-----MIAAHADTNSLVLTA-PQDIMNAMLEVIGQLDIRRAQVLIEALIVEMAEGDGINLGVQWGSL-----ESG---SVIQYGN-TGASIGNVMIGLEEAKDTTQTKAVYDTNNNFLRNETTTTK 418
A.salmonicida   -ASIGGGKLAISADETTNALVITA-QPDVMAELEQVVAKLDIRRAQVLVEAIIVEIADGDGLNLGVQWANT-----NGG----GTQFTN-AGPGIGSVAIAAKDYKDN---------------------- 404
P.aeruginosa    PSNI-----LIRADESTNALVLLA-DPDTVNALEDIVRQLDVPRAQVLVEAAIVEISGDIQDAVGVQWAIN-----KGGMGGTKTNFAN-TGLSIGTLLQSLESNKAP---------------------- 416
N.meningtitis   ------G--SVLIDPATNTLIVTD-TRSVIEKFRKLIDELDVPAQQVMIEARIVEAADGFSRDLGVKFG-------ATG----KKKLKN-DTSAFGWGVNSGFGGDDK--------WGAETKINLPI--- 568
Myxococcus      ------G--SVTVDQRTNVLIVKD-VRSNTERARSLVRSLDTQTPQVLIESRIVEANTSFSRSLGVQWGGQARAGQATG-NSTGLIFPN-NLAVTGGVTGTGAGLPDN----------PNFAVNLPT--- 707
S.enterica      ---I-----KIVAYPDTNSLLVKG-TAEQVHFIEMLVKALDVAKRHVELSLWIVDLNKSDLERLGTSWSGSITIGDKLG--------------------------------------------------- 206
```

```
                        790       800       810       820       830       840       850       860       870       880       890       900       910
               ....|....|....|....|....|....|....|....|....|....|....|....|....|....|....|....|....|....|....|....|....|....|....|....|....|....|
AgoGspDN1      ---------------------------------------------------------------------------------------------------------------------------------- 101
RamGspDN1      ---------------------------------------------------------------------------------------------------------------------------------- 155
NgrGspDN1      ---------------------------------------------------------------------------------------------------------------------------------- 121
NfoGspDN1      ---------------------------------------------------------------------------------------------------------------------------------- 120
NdaGspDN1      ---------------------------------------------------------------------------------------------------------------------------------- 108
PcoAEGspDN1    ---------------------------------------------------------------------------------------------------------------------------------- 119
PkiGspDN1      ---------------------------------------------------------------------------------------------------------------------------------- 135
GokGspDN1      ---------------------------------------------------------------------------------------------------------------------------------- 102
SpiGspDN1      ---------------------------------------------------------------------------------------------------------------------------------- 121
E.coli         GDLSTLAQLLSGFSGTAVGVVKGDWMALVQAVKNDSSSNVLSTPSITTLDNQEAFFMVGQDVPVLTGS--TVGSNNSNPFNTVERKKVGIMLKVTPQINEGNAVQMVIEQEV-SKV---EGQT-SLD--V 555
V.cholerae     GDYTKLASALSSIQGAAVSIAMGDWTALINAVSNDSSSNILSSPSITVMDNGEASFIVGEEVPVITGS--TAGSNNDNPFQTVDRKEVGIKLKVVPQINEGNSVQLNIEQEV-SNV---LGANGAVD--V 540
A.salmonicida  GTTTGIAKLAESFNGMAAGFYQGNWAMLVTALSTNTKSDILSTPSIVTMDNKEASFNVGQEVPVQTGK--QNSTSGDTTFSTIERKTVGTKLVVTPQINEGDSVLLTIEQEV-SSVGKQASGTEGLG--P 529
P.aeruginosa   ---------ESIPDGAIVGIGSSSFGALVTALSANTKSNLLSTPSLLTLDNQKAEILVGQNVPFQTGSYTTNSEGSSNPFTTVERKDIGVSLKVTPHINDGAALRLEIEQEI-SALLPNAQQRNNTD--L 534
N.meningtitis  -----TAAANSISLVRAIS--SGALNLELSASESLSKTKTLANPRVLTQNRKEAKIESGYEIPFTVTSIANGGSSTNTEL-----KKAVLGLTVTPNITPDGQIIMTVKITK-DSPAQCASGN-QTI--L 682
Myxococcus     GTGQGVGGAMGFTFGSAGG--ALQLNLRLSAAENEGSVKTISAPKVTTLDNNTARISQGVSIPFSQTS----AQGVNTTF-----VEARLSLEVTPHITQDGSVLMSINASN-NQPDPSSTGA-NGQ--P 822
S.enterica     ---------VSLNQSSISTLDGSRFIAAVNALEEKKQATVVSRPVLLTQENVPAIFDNNRTFYTKLIG------ERNVAL---EHVTYGTMIRVLPRFSADGQIEMSLDIEDGNDKTPQSDTTTSVDALP 318

                        920       930       940       950       960       970       980       990      1000      1010      1020      1030      1040
               ....|....|....|....|....|....|....|....|....|....|....|....|....|....|....|....|....|....|....|....|....|....|....|....|....|....|
AgoGspDN1      ---------------------------------------------------------------------------------------------------------------------------------- 101
RamGspDN1      ---------------------------------------------------------------------------------------------------------------------------------- 155
NgrGspDN1      ---------------------------------------------------------------------------------------------------------------------------------- 121
NfoGspDN1      ---------------------------------------------------------------------------------------------------------------------------------- 120
NdaGspDN1      ---------------------------------------------------------------------------------------------------------------------------------- 108
PcoAEGspDN1    ---------------------------------------------------------------------------------------------------------------------------------- 119
PkiGspDN1      ---------------------------------------------------------------------------------------------------------------------------------- 135
GokGspDN1      ---------------------------------------------------------------------------------------------------------------------------------- 102
SpiGspDN1      ---------------------------------------------------------------------------------------------------------------------------------- 121
E.coli         VFGERKLKTTVLANDGELIVLGGLMDDQAGESVAKVPLLGDIPLIGNLFKSTADKKEKRNLMVFIRPTILRDGMAADGVSQRKYNYMRAEQIYRDEQ-GLSLMPHTAQPILP--AQNQALPPEVRAFLNA 682
V.cholerae     RFAKRQLNTSVMVQDGQMLVLGGLIDERALESESKVPLLGDIPLLGQLFRSTSSQVEKKNLMVFIKPTIIRDGVTADGITQRKYNYIRAEQLFRAEK-GLRLLDDASVPVLPKFGDDRRHSPEIQAFIEQ 669
A.salmonicida  TFDTRTVKNAVLVKSGETVVLGGLMDEQTKEEVSKVPLLGDIPVLGYLFRSTSNTTSKRNLMVFIRPTILRDANVYSGISSNKYTLFRAQQLEAAAQKGYATSPDRQ--VLPEYGQDVVQSPEIQKQIEL 657
P.aeruginosa   ITSKRSIKSTILAENGQVIVIGGLIQDDVSQAESKVPLLGDIPLLGRLFRSTKDTHTKRNLMVFLRPTVVRDSAGLAALSGKKYSDIRVIDGTRGPE-GRPS-------ILPTNANQLFDGQAVDLR-EL 655
N.meningtitis  CISTKNLNTQAMVENGGTLIVGGIYEEDNGNTLTKVPLLGDIPVIGNLFKTRGKKTDRRELLIFITPRIMGTAG----------NSLRY----------------------------------------- 761
Myxococcus     SIQRKEANTQVLVKDGDTTVIGGIYVRRGATQVNSVPFLSRIPVLGLLFKNNSETDTRQELLIFITPRILNRQTIAQTL--------------------------------------------------- 901
S.enterica     EVGRTLISTIARVPHGKSLLVGGYTRDANTDTVQSIPFLGKLPLIGSLFRYSSKNKSNVVRVFMIEPKEIVDPLTPDA-SESVNNILKQSGAWSGDD---------------------KLQKWVRVYLDR 426

                       1050      1060
               ....|....|....|....|
AgoGspDN1      -------------------- 101
RamGspDN1      -------------------- 155
NgrGspDN1      -------------------- 121
NfoGspDN1      -------------------- 120
NdaGspDN1      -------------------- 108
PcoAEGspDN1    -------------------- 119
PkiGspDN1      -------------------- 135
GokGspDN1      -------------------- 102
SpiGspDN1      -------------------- 121
E.coli         GRTR---------------- 686
V.cholerae     MEAKQ--------------- 674
A.salmonicida  MKARQQATADGAQPFVQGNN 677
P.aeruginosa   MTE----------------- 658
N.meningtitis  -------------------- 761
Myxococcus     -------------------- 901
S.enterica     GQEAIK-------------- 432
```

### (C) GspDN2

```
                       10        20        30        40        50        60        70        80        90       100       110       120       130
               ....|....|....|....|....|....|....|....|....|....|....|....|....|....|....|....|....|....|....|....|....|....|....|....|....|....|
AgoGspDN2      ---------------------------------------------------------------------------------------------------------------------------------- 1
RamGspDN2      ---------------------------------------------------------------------------------------------------------------------------------- 1
NgrGspDN2      ---------------------------------------------------------------------------------------------------------------------------------- 1
NfoGspDN2      ---------------------------------------------------------------------------------------------------------------------------------- 1
NdaGspDN2      ---------------------------------------------------------------------------------------------------------------------------------- 1
PcoWSGspDN2    ---------------------------------------------------------------------------------------------------------------------------------- 1
PcoAEGspDN2    ---------------------------------------------------------------------------------------------------------------------------------- 1
PkiGspDN2      ---------------------------------------------------------------------------------------------------------------------------------- 1
MjaGspDN2      ---------------------------------------------------------------------------------------------------------------------------------- 1
GokGspDN2      ---------------------------------------------------------------------------------------------------------------------------------- 1
SpiGspDN2      ---------------------------------------------------------------------------------------------------------------------------------- 1
E.coli         MFWR--------DMT----------------------------------------------LSIWRKKTTGLKTKKR-LLPLMLAA-----------------------ALCSSPVWAEEATFTANFKDT 52
V.cholerae     -------------------------------------------------------------MKYWLKKSSWL-----------LAG-----------------------SLLSTPL-AMANEFSASFKGT 34
A.salmonicida  ---------------------------------------------------------------MINKGKSWR-------LAT-VAA-----------------------ALMMAGS-AWATEYSASFKNA 35
P.aeruginosa   MSQP--------LLR----------------------------------------------ALFAPSSRSYVPAVLL-SLALGIQAAH--------------AENSGGNAFVPAGN-QQEAHWTINLKDA 60
N.meningtitis  MNTK--------LTKIISGLFVATAAFQTASAGNITDIKVSSLPNKQKIVKVSFDKEIVNPTGFVTSSPARIALDF---EQTGISMDQQVLEYADPLLSKISAAQNSSRARLVLNL-NKPGQYNTEVRGN 118
M.xanthus      MLEESAVTRGKWMLAAWAVVLVGARVHGAELNTLRGLDVSRTGSGAQVVVTGTRPPTF--TVFRLSGPERLVVDLSSADATGIKGHHEGSGPVSGVVASQFSDQRASVGRVLLAL-DKASQYDVRADGN 127
S.enterica     LYNKNYPLRGD-----------------------------------------------------------------------------------------------------------NRKGTF------ 17

                      140       150       160       170       180       190       200       210       220       230       240       250       260
               ....|....|....|....|....|....|....|....|....|....|....|....|....|....|....|....|....|....|....|....|....|....|....|....|....|....|
AgoGspDN2      ---------------------------------------------------------------------------------------------------------------------------------- 1
RamGspDN2      ---------------------------------------------------------------------------------------------------------------------------------- 1
NgrGspDN2      ---------------------------------------------------------------------------------------------------------------------------------- 1
NfoGspDN2      ---------------------------------------------------------------------------------------------------------------------------------- 1
NdaGspDN2      ---------------------------------------------------------------------------------------------------------------------------------- 1
PcoWSGspDN2    ---------------------------------------------------------------------------------------------------------------------------------- 1
PcoAEGspDN2    ---------------------------------------------------------------------------------------------------------------------------------- 1
PkiGspDN2      ---------------------------------------------------------------------------------------------------------------------------------- 1
MjaGspDN2      ---------------------------------------------------------------------------------------------------------------------------------- 1
GokGspDN2      ---------------------------------------------------------------------------------------------------------------------------------- 1
SpiGspDN2      ---------------------------------------------------------------------------------------------------------------------------------- 1
E.coli         DLKSFIETVGANLNK--------------TIIMGPG----------VQGKVSIRTMTPLNER------------------QYYQLFLNLLEAQG----------------------YAV----------- 107
V.cholerae     DIQEFINIVGRNLEK--------------TIIVDPS----------VRGKVDVRSFDTLNEE------------------QYYSFFLSVLEVYG----------------------FAV----------- 89
A.salmonicida  DIEEFINTVGKNLSK--------------TIIIEPS----------VRGKINVRSYDLLNEE------------------QYYQFFLSVLDVYG----------------------FAV----------- 90
P.aeruginosa   DIREFIDQISEITGE--------------TFVVDPR----------VKGQVSVVSKAQLSLS------------------EVYQLFLSVMSTHG----------------------FTV----------- 115
N.meningtitis  KVWIFINESDDTVSA-----PAR-----PAVKAAPAA-------PAKQQAAAPSTKSAVSVSE---------PFTPAKQQAAAPFTESVVSVSAP---------------------FSP----------- 190
M.xanthus      RVVISVDGAAQSVEAKRAEPPARTEGVTASVEVKPHSVSAAAPAKVVQAESAAVSKAALPENVVAAEADEREVSNPAQHITAMSFADDTLSIRADGDIARYEVLELADPPRLAVDLFGVGLATRAPRVKS 257
S.enterica     ---------------------------------------------------------------------------------------------------------------------------------- 17

                      270       280       290       300       310       320       330       340       350       360       370       380       390
               ....|....|....|....|....|....|....|....|....|....|....|....|....|....|....|....|....|....|....|....|....|....|....|....|....|....|
AgoGspDN2      ---------------------------------------------------------------------------------------------------------------------------------- 1
RamGspDN2      ---------------------------------------------------------------------------------------------------------------------------------- 1
NgrGspDN2      ---------------------------------------------------------------------------------------------------------------------------------- 1
NfoGspDN2      ---------------------------------------------------------------------------------------------------------------------------------- 1
NdaGspDN2      ---------------------------------------------------------------------------------------------------------------------------------- 1
PcoWSGspDN2    ---------------------------------------------------------------------------------------------------------------------------------- 1
PcoAEGspDN2    ---------------------------------------------------------------------------------------------------------------------------------- 1
PkiGspDN2      ---------------------------------------------------------------------------------------------------------------------------------- 1
MjaGspDN2      ---------------------------------------------------------------------------------------------------------------------------------- 1
GokGspDN2      ---------------------------------------------------------------------------------------------------------------------------------- 1
SpiGspDN2      ---------------------------------------------------------------------------------------------------------------------------------- 1
E.coli         -----------------------------VPMENDVLKVV----------------------------KSSAAKVEPLPLVG----EGSDNYAGDEMVTKV----------------------------- 147
V.cholerae     -----------------------------VEMDNGVLKVI----------------------------KSKDAKTSAIPVL-----SGEERANGDEVITQV----------------------------- 128
A.salmonicida  -----------------------------VPMDNGVLKVV----------------------------RSKDAKTSAIPVV-----DETNPGIGDEMVTRV----------------------------- 129
P.aeruginosa   -----------------------------VAQGDQA-RIV----------------------------PNAEAKTEA----------GGGQSAPDRLETRV----------------------------- 148
N.meningtitis  ----------------------------------------A----KQQAAASAKQQ------------AAAPAKQQAAAPAKQTNIDFRKDGKNAGIIELAALGFAGQPDISQQHDHIIVTLKNHTLPTT 264
M.xanthus      GALRDVRVGAHADKVRLVLDVRGTMPAYRVDRANRGLEVVLGRAVARKAAPAVETQAVVASVAEVEPLRQAPVKSEASPVVEVKDVRFEESSAGGRIVMKLSGTSAWKVDRPDPRSAVL-TLENARLPKK 386
S.enterica     ---------------------------------------------------------------------------------------------------------------------------------- 17

                      400       410       420       430       440       450       460       470       480       490       500       510       520
               ....|....|....|....|....|....|....|....|....|....|....|....|....|....|....|....|....|....|....|....|....|....|....|....|....|....|
AgoGspDN2      ---------------------------------------------------------------------------------------------------------------------------------- 1
RamGspDN2      ---------------------------------------------------------------------------------------------------------------------------------- 1
NgrGspDN2      ---------------------------------------------------------------------------------------------------------------------------------- 1
NfoGspDN2      ---------------------------------------------------------------------------------------------------------------------------------- 1
NdaGspDN2      ---------------------------------------------------------------------------------------------------------------------------------- 1
PcoWSGspDN2    ---------------------------------------------------------------------------------------------------------------------------------- 1
PcoAEGspDN2    ---------------------------------------------------------------------------------------------------------------------------------- 1
PkiGspDN2      ---------------------------------------------------------------------------------------------------------------------------------- 1
MjaGspDN2      ---------------------------------------------------------------------------------------------------------------------------------- 1
GokGspDN2      ---------------------------------------------------------------------------------------------------------------------------------- 1
SpiGspDN2      ---------------------------------------------------------------------------------------------------------------------------------- 1
E.coli         -----------VPVRNVSVRELAPI---LRQMIDSAGSGN-VVNYDP-----------------------------------------------SNVIMLTGRASVVERLTEVIQRV---------DHAG 206
V.cholerae     -----------VAVKNVSVRELSPL---LRQLIDNAGAGN-VVHYDP-----------------------------------------------ANIILITGRAAVVNRLAEIIRRV---------DQAG 187
A.salmonicida  -----------VPVRNVSVRELAPL---LRQLNDNAGGGN-VVHYDP-----------------------------------------------SNVLLITGRAAVVNRLVEVVRRV---------DKAG 188
P.aeruginosa   -----------IQVQQSPVSELIPL---IRPLVPQ--YGH-LAAVPS-----------------------------------------------ANALIISDRSANIARIEDVIRQL---------DQKG 205
N.meningtitis  LQRSLDVADFKTPVQKVTLKRLNND---TQLIITTAGNWE-LVNKSAAPGYFTFQVLPKKQNLESGGVNNAPKTF------------------TGRKISLDFQDVEIRTILQILAKESGMNIVASDSVNG 372
M.xanthus      FERSLDTSALDTPVKMISAFSVPGEGGKVRLVVAADGAIEEKVSQSA--GTLSWRLDVKGVKTEEVAVAQRTAGFTTEAPAYAAEGAPQQARYRGKRVSFEFKDIDIQNLLRVIAEISKKNIVVADDVSG 514
S.enterica     -----------------------------------------YVSGPP-----------------------------------------------VYVDMVVNAATMMDKQNDGIE-------------LG 46

                      530       540       550       560       570       580       590       600       610       620       630       640       650
               ....|....|....|....|....|....|....|....|....|....|....|....|....|....|....|....|....|....|....|....|....|....|....|....|....|....|
AgoGspDN2      --------------------------------------------------------------------------------------------------MTSNGSSSGGHEALSTPLPLLQSSVKAYVPPS 32
RamGspDN2      ----------------------------------------------------------------------------------------------------------------MDDGERFRSTVFSQRAPA 18
NgrGspDN2      ------------------------------------------MQESKQTNNGSVLNRQQDNKNEKQFLSETNTHETDNDKQINN-------KIEDGEQNMNTTTTAQPTLNNYQ-QVHHRSQVKTYTPPV 80
NfoGspDN2      ---------------------------------------------------------------------------------------------------MNNSS--------------LRSQVKTYTPPP 17
NdaGspDN2      ---------------------------------------------------------------------------------------------------MNKK------EETPKTSVDEKSTVKTYRPP- 24
PcoWSGspDN2    ----------------------------------------------------------------------------------------MSPSQPAAPTKPHANRRSKNPRDNKPKNTSMKSSVKTFRPPP 42
PcoAEGspDN2    ------------------------------------------------------------------------------------------------------------------------STVKTYRPP- 9
PkiGspDN2      ------------------------------------------MPSSKEENESQPSPQSPPHEEHKNPSLPRKKSKRSSSKQKKNRSSKQKQKQQEQKQKQNQKQQKQPPETTHRPSSSTKSTVKTYRPP- 87
MjaGspDN2      -----------------------------------------------------------------------------------------------------------------------MSTVSSFRPPE 11
GokGspDN2      ----------------------------------------------------------------------------------------------------------MVDPPPAAAIDDNGSSVACYRTPQ 24
SpiGspDN2      ---------------------------------------------------------------------------------------------------------------------------------- 1
E.coli         NRTEEVIPLDNASASEIARVLESLTKNS---GENQPATLKSQIVADERTNSVIVSGDPATRDKMRRLIRRLDSEMERSGNS----------QVFYLKYSKAEDLVDVLKQVSGTLTAAKEEAEGTVGSGR 323
V.cholerae     DKEIEVVELNNASAAEMVRIVEALNKTTD--AQNTPEFLKPKFVADERTNSILISGDPKVRERLKRLIKQLDVEMAAKGNN----------RVVYLKYAKAEDLVEVLKGVSENLQAEKGTGQPTT-SKR 304
A.salmonicida  DQEVDIIKLKYASAGEMVRLVTNLNKDGNTQGGNTSLLLAPKVVADERTNSVVVSGEPKARARIIQMVRQLDRDLQSQGNT----------RVFYLKYGKAKDMVEVLKGVSTSIEADKKGGGATAGGGN 308
P.aeruginosa   SHDYSVINLRYGWVMDAAEVLNNAMSRGQAKGA-----AGAQVIADARTNRLIILGPPQARAKLVQLAQSLDTPTARSANT----------RVIRLRHNDAKTLAETLGQISEGMKNNGGQGGEQTGGGR 320
N.meningtitis  KMTLSLKDVPWDQALDLV--------------------MQARNLDMRQQGNIVNIAP------RDELLAKDKAFLQAEKDIA-D-LGALYSQNFQLKYKNVEEFRSILRLDNADTTGNRN----TLVSGR 470
M.xanthus      KVTIRLRNVPWDQALDLV--------------------LRTKALGKEEFGNIIRIAPLKTLEEEARLRQERKASLQRQEDLMV--------NLIPVNYAVAADMAARVK---------------DVLSER 601
S.enterica     RQKIGVMRLNNTFVGDRT--------------------------YNLRDQKMVIPGIATAIERLLQGEEQPLGNIVSSEPPAMPAFSANGEKGKAANYAGGMSLQEALKQ--------------NAAAGN 136

                      660       670       680       690       700       710       720       730       740       750       760       770       780
               ....|....|....|....|....|....|....|....|....|....|....|....|....|....|....|....|....|....|....|....|....|....|....|....|....|....|
AgoGspDN2      MTDGG----SGGDEPGRPLLCIHTRPSYMKRISEVLLRLDTPPQMLQLEFQLVEVAEDDLTQLRHRINHPVLNRIAGESAMQDDVFSRRAYSPMDIRAANTSGKCNVLQPQMQVQSRL------------ 146
RamGspDN2      ELAGE--SGGGGSGQAAAVVCVHTYPHLMRRLVAFLRHFDKPPKNLSLEFTFVEVADEDVSQFRYHLQHPLLRSALRAPAPP-E----------MRQPPEPRGLPSQLSANL------------------ 117
NgrGspDN2      LGSP---SGSSGGIGTTPVVSIYTKPHLMKRLKQFLDHFDRAPQQMFIDFAFVEVEENDVSQFRYKLQHPILQSIYGPTVDEQL------PLYQNESTSSSARLPSSLTPRL------------------ 183
NfoGspDN2      MSDTPHSSPSSSVASTTPVVSIYTKPHLMRRLKQFLDHFDRPPQQLFIDFAFVEVEENDVSQFRYKLQHPILQSIYGPTVDEQL------PLYQNESTSSSARLPTSLTPRL------------------ 123
NdaGspDN2      -------------DGSTPVVCIHTKPSLMKKLMSFLQHFDRPPQQLFLEFVFVEVEENDAAQFRYRLNHPMLNSITGGDIDSDLLNMQ--PFQQTS-NQPSTRLPSELNPRM------------------ 120
PcoWSGspDN2    GDDG--------ATLQSSMLLIHTYPSLHQRLLQFLSHFDKPPQQMYFELLFTEMRDVDVKQMRWKFGDNVLSG-----AQMFD------QMVNGMNQMENVRLPQELQGKM------------------ 135
PcoAEGspDN2    --DG-----------SKPMICIHTKPTLHKRLLKFLEHFDKPPQQLYLEFSFVEVAEDDVAQFRYKLDHPVLNEIMG-TDGVYS------SMIN-AESTP-THLPSSLQPKM------------------ 99
PkiGspDN2      -------------DGSPPVLCIHTRPHLMKKLRAFFEHFDRPPQQLFLEFVFVEVAEEDAAQFRYRLEHPVIDSVYGRAADADV------PLSQRE-RRRHPELPDELLPRL------------------ 179
MjaGspDN2      ----------GGQ----PVVAIHTTPALMKKLVAFLKHFDRPPQQVYLEFVFLEVAEDDLAEFRYNVQHPLLYRLLGR-APP------------VNAPMAGSDAAAAPPPSL------------------ 96
GokGspDN2      ----------GGQ----PVIAVHTTPALMRKLVAFLRHFDRPPQQLFLEFVFLEVSEDELNEFRYNLQHPLLYRLLGR-TPP------------ATAPAPSAASTSTAAPV------------------- 108
SpiGspDN2      ----------GGQ----PVVCIHTTPALMARLTEFLRHFDRPPQQLFLEFLFVEMASEELEQLRYHLRHPLLSQMLGAPVPPSA----------VTETP----LPEQEP--------------------- 81
E.coli         -EVV-----SIAASKHSNALIVTAPQDIMQSLQSVIEQLDIRRAQVHVEALIVEVAEGSNINFGVQWGSK-----DAG----LMQFANGTQIPIGTLGAAISAAKPQKGSTVISENGATTINPDTN---G 435
V.cholerae     -NEV-----MIAAHADTNSLVLTAPQDIMNAMLEVIGQLDIRRAQVLIEALIVEMAEGDGINLGVQWGSL-----ESG---SVIQYGN-TGASIGNVMIGLEEAKDTTQTKAVYDTNNNFLRNETTTTKG 419
A.salmonicida  -ASIGGGKLAISADETTNALVITAQPDVMAELEQVVAKLDIRRAQVLVEAIIVEIADGDGLNLGVQWANT-----NGG----GTQFTN-AGPGIGSVAIAAKDYKDN----------------------G 405
P.aeruginosa   PSNI-----LIRADESTNALVLLADPDTVNALEDIVRQLDVPRAQVLVEAAIVEISGDIQDAVGVQWAIN-----KGGMGGTKTNFAN-TGLSIGTLLQSLESNKAP----------------------- 416
N.meningtitis  ------G--SVLIDPATNTLIVTDTRSVIEKFRKLIDELDVPAQQVMIEARIVEAADGFSRDLGVKFG-------ATG----KKKLKN-DTSAFGWGVNSGFGGDDK--------WGAETKINLPI---- 568
M.xanthus      ------G--SVTVDQRTNVLIVKDVRSNTERARSLVRSLDTQTPQVLIESRIVEANTSFSRSLGVQWGGQARAGQATG-NSTGLIFPN-NLAVTGGVTGTGAGLPDN----------PNFAVNLPT---G 708
S.enterica     ---I-----KIVAYPDTNSLLVKGTAEQVHFIEMLVKALDVAKRHVELSLWIVDLNKSDLERLGTSWSGSITIGDKLG---------------------------------------------------- 206
```

```
                790       800       810       820       830       840       850       860       870       880       890       900       910
        ....|....|....|....|....|....|....|....|....|....|....|....|....|....|....|....|....|....|....|....|....|....|....|....|....|....|
AgoGspDN2      ---------------------------------------------------------------------------------------------------------------------------------- 146
RamGspDN2      ---------------------------------------------------------------------------------------------------------------------------------- 117
NgrGspDN2      ---------------------------------------------------------------------------------------------------------------------------------- 183
NfoGspDN2      ---------------------------------------------------------------------------------------------------------------------------------- 123
NdaGspDN2      ---------------------------------------------------------------------------------------------------------------------------------- 120
PcoWSGspDN2    ---------------------------------------------------------------------------------------------------------------------------------- 135
PcoAEGspDN2    ---------------------------------------------------------------------------------------------------------------------------------- 99
PkiGspDN2      ---------------------------------------------------------------------------------------------------------------------------------- 179
MjaGspDN2      ---------------------------------------------------------------------------------------------------------------------------------- 96
GokGspDN2      ---------------------------------------------------------------------------------------------------------------------------------- 108
SpiGspDN2      ---------------------------------------------------------------------------------------------------------------------------------- 81
E.coli         DLSTLAQLLSGFSGTAVGVVKGDWMALVQAVKNDSSSNVLSTPSITTLDNQEAFFMVGQDVPVLTGS--TVGSNNSNPFNTVERKKVGIMLKVTPQINEGNAVQMVIEQEV-SKV---EGQT-SLD--VV 556
V.cholerae     DYTKLASALSSIQGAAVSIAMGDWTALINAVSNDSSSNILSSPSITVMDNGEASFIVGEEVPVITGS--TAGSNNDNPFQTVDRKEVGIKLKVVPQINEGNSVQLNIEQEV-SNV---LGANGAVD--VR 541
A.salmonicida  TTTGIAKLAESFNGMAAGFYQGNWAMLVTALSTNTKSDILSTPSIVTMDNKEASFNVGQEVPVQTGK--QNSTSGDTTFSTIERKTVGTKLVVTPQINEGDSVLLTIEQEV-SSVGKQASGTEGLG--PT 530
P.aeruginosa   --------ESIPDGAIVGIGSSSFGALVTALSANTKSNLLSTPSLLTLDNQKAEILVGQNVPFQTGSYTTNSEGSSNPFTTVERKDIGVSLKVTPHINDGAALRLEIEQEI-SALLPNAQQRNNTD--LI 535
N.meningtitis  ----TAAANSISLVRAIS--SGALNLELSASESLSKTKTLANPRVLTQNRKEAKIESGYEIPFTVTSIANGGSSTNTEL-----KKAVLGLTVTPNITPDGQIIMTVKITK-DSPAQCASGN-QTI--LC 683
M.xanthus      TGQGVGGAMGFTFGSAGG--ALQLNLRLSAAENEGSVKTISAPKVTTLDNNTARISQGVSIPFSQTS----AQGVNTTF-----VEARLSLEVTPHITQDGSVLMSINASN-NQPDPSSTGA-NGQ--PS 823
S.enterica     --------VSLNQSSISTLDGSRFIAAVNALEEKKQATVVSRPVLLTQENVPAIFDNNRTFYTKLIG------ERNVAL---EHVTYGTMIRVLPRFSADGQIEMSLDIEDGNDKTPQSDTTTSVDALPE 319

                920       930       940       950       960       970       980       990      1000      1010      1020      1030      1040
        ....|....|....|....|....|....|....|....|....|....|....|....|....|....|....|....|....|....|....|....|....|....|....|....|....|
AgoGspDN2      ---------------------------------------------------------------------------------------------------------------------------------- 146
RamGspDN2      ---------------------------------------------------------------------------------------------------------------------------------- 117
NgrGspDN2      ---------------------------------------------------------------------------------------------------------------------------------- 183
NfoGspDN2      ---------------------------------------------------------------------------------------------------------------------------------- 123
NdaGspDN2      ---------------------------------------------------------------------------------------------------------------------------------- 120
PcoWSGspDN2    ---------------------------------------------------------------------------------------------------------------------------------- 135
PcoAEGspDN2    ---------------------------------------------------------------------------------------------------------------------------------- 99
PkiGspDN2      ---------------------------------------------------------------------------------------------------------------------------------- 179
MjaGspDN2      ---------------------------------------------------------------------------------------------------------------------------------- 96
GokGspDN2      ---------------------------------------------------------------------------------------------------------------------------------- 108
SpiGspDN2      ---------------------------------------------------------------------------------------------------------------------------------- 81
E.coli         FGERKLKTTVLANDGELIVLGGLMDDQAGESVAKVPLLGDIPLIGNLFKSTADKKEKRNLMVFIRPTILRDGMAADGVSQRKYNYMRAEQIYRDEQ-GLSLMPHTAQPILP--AQNQALPPEVRAFLNAG 683
V.cholerae     FAKRQLNTSVMVQDGQMLVLGGLIDERALESESKVPLLGDIPLLGQLFRSTSSQVEKKNLMVFIKPTIIRDGVTADGITQRKYNYIRAEQLFRAEK-GLRLLDDASVPVLPKFGDDRRHSPEIQAFIEQM 670
A.salmonicida  FDTRTVKNAVLVKSGETVVLGGLMDEQTKEEVSKVPLLGDIPVLGYLFRSTSNTTSKRNLMVFIRPTILRDANVYSGISSNKYTLFRAQQLEAAAQKGYATSPDRQ--VLPEYGQDVVQSPEIQKQIELM 658
P.aeruginosa   TSKRSIKSTILAENGQVIVIGGLIQDDVSQAESKVPLLGDIPLLGRLFRSTKDTHTKRNLMVFLRPTVVRDSAGLAALSGKKYSDIRVIDGTRGPE-GRPS-------ILPTNANQLFDGQAVDLR-ELM 656
N.meningtitis  ISTKNLNTQAMVENGGTLIVGGIYEEDNGNTLTKVPLLGDIPVIGNLFKTRGKKTDRRELLIFITPRIMGTAG----------NSLRY----------------------------------------- 761
M.xanthus      IQRKEANTQVLVKDGDTTVIGGIYVRRGATQVNSVPFLSRIPVLGLLFKNNSETDTRQELLIFITPRILNRQTIAQTL---------------------------------------------------- 901
S.enterica     VGRTLISTIARVPHGKSLLVGGYTRDANTDTVQSIPFLGKLPLIGSLFRYSSKNKSNVVRVFMIEPKEIVDPLTPDA-SESVNNILKQSGAWSGDD--------------------KLQKWVRVYLDRG 427

                1050
        ....|....|....|....
AgoGspDN2      ------------------- 146
RamGspDN2      ------------------- 117
NgrGspDN2      ------------------- 183
NfoGspDN2      ------------------- 123
NdaGspDN2      ------------------- 120
PcoWSGspDN2    ------------------- 135
PcoAEGspDN2    ------------------- 99
PkiGspDN2      ------------------- 179
MjaGspDN2      ------------------- 96
GokGspDN2      ------------------- 108
SpiGspDN2      ------------------- 81
E.coli         RTR---------------- 686
V.cholerae     EAKQ--------------- 674
A.salmonicida  KARQQATADGAQPFVQGNN 677
P.aeruginosa   TE----------------- 658
N.meningtitis  ------------------- 761
M.xanthus      ------------------- 901
S.enterica     QEAIK-------------- 432
```

### (D) GspDN3

```
                        10        20        30        40        50        60        70        80        90       100       110       120       130
                ....|....|....|....|....|....|....|....|....|....|....|....|....|....|....|....|....|....|....|....|....|....|....|....|....|....|
AgoDN3          ---------------------------------------------------------------------------------------------------------------------------------- 1
RamDN3          ---------------------------------------------------------------------------------------------------------------------------------- 1
NgrDN3          ---------------------------------------------------------------------------------------------------------------------------------- 1
NfoDN3          ---------------------------------------------------------------------------------------------------------------------------------- 1
PcoAEDN3        ---------------------------------------------------------------------------------------------------------------------------------- 1
PcoWSDN3        ---------------------------------------------------------------------------------------------------------------------------------- 1
NdaDN3          ---------------------------------------------------------------------------------------------------------------------------------- 1
PkiDN3          ---------------------------------------------------------------------------------------------------------------------------------- 1
MjaDN3          ---------------------------------------------------------------------------------------------------------------------------------- 1
GokDN3          ---------------------------------------------------------------------------------------------------------------------------------- 1
E.coli          MFWR--------DMT----------------------------------------------LSIWRKKTTGLKTKKR-LLPLMLAA-----------------------ALCSSPVWAEEATFTANFKDT 52
V.cholerae      -------------------------------------------------------------MKYWLKKSSWL-----------LAG-----------------------SLLSTPL-AMANEFSASFKGT 34
A.salmonicida   ---------------------------------------------------------------MINKGKSWR-------LAT-VAA-----------------------ALMMAGS-AWATEYSASFKNA 35
P.aeruginosa    MSQP--------LLR----------------------------------------------ALFAPSSRSYVPAVLL-SLALGIQAAH--------------AENSGGNAFVPAGN-QQEAHWTINLKDA 60
N.meningtitis   MNTK--------LTKIISGLFVATAAFQTASAGNITDIKVSSLPNKQKIVKVSFDKEIVNPTGFVTSSPARIALDF---EQTGISMDQQVLEYADPLLSKISAAQNSSRARLVLNL-NKPGQYNTEVRGN 118
M.xanthus       MLEESAVTRGKWMLAAAWAVVLVGARVHGAELNTLRGLDVSRTGSGAQVVVTGTRPPTF--TVFRLSGPERLVVDLSSADATGIKGHHEGSGPVSGVVASQFSDQRASVGRVLLAL-DKASQYDVRADGN 127
S.enterica      LYNKNYPLRGD----------------------------------------------------------------------------------------------------------NRKGTF------- 17

                       140       150       160       170       180       190       200       210       220       230       240       250       260
                ....|....|....|....|....|....|....|....|....|....|....|....|....|....|....|....|....|....|....|....|....|....|....|....|....|....|
AgoDN3          ---------------------------------------------------------------------------------------------------------------------------------- 1
RamDN3          ---------------------------------------------------------------------------------------------------------------------------------- 1
NgrDN3          ---------------------------------------------------------------------------------------------------------------------------------- 1
NfoDN3          ---------------------------------------------------------------------------------------------------------------------------------- 1
PcoAEDN3        ---------------------------------------------------------------------------------------------------------------------------------- 1
PcoWSDN3        ---------------------------------------------------------------------------------------------------------------------------------- 1
NdaDN3          ---------------------------------------------------------------------------------------------------------------------------------- 1
PkiDN3          ---------------------------------------------------------------------------------------------------------------------------------- 1
MjaDN3          ---------------------------------------------------------------------------------------------------------------------------------- 1
GokDN3          ---------------------------------------------------------------------------------------------------------------------------------- 1
E.coli          DLKSFIETVGANLNK--------------TIIMGPG----------VQGKVSIRTMTPLNER------------------QYYQLFLNLLEAQG----------------------YAV----------- 107
V.cholerae      DIQEFINIVGRNLEK--------------TIIVDPS----------VRGKVDVRSFDTLNEE------------------QYYSFFLSVLEVYG----------------------FAV----------- 89
A.salmonicida   DIEEFINTVGKNLSK--------------TIIIEPS----------VRGKINVRSYDLLNEE------------------QYYQFFLSVLDVYG----------------------FAV----------- 90
P.aeruginosa    DIREFIDQISEITGE--------------TFVVDPR----------VKGQVSVVSKAQLSLS------------------EVYQLFLSVMSTHG----------------------FTV----------- 115
N.meningtitis   KVWIFINESDDTVSA-----PAR-----PAVKAAPAA-------PAKQQAAAPSTKSAVSVSE---------PFTPAKQQAAAPFTESVVSVSAP---------------------FSP----------- 190
M.xanthus       RVVISVDGAAQSVEAKRAEPPARTEGVTASVEVKPHSVSAAAPAKVVQAESAAVSKAALPENVVAAEADEREVSNPAQHITAMSFADDTLSIRADGDIARYEVLELADPPRLAVDLFGVGLATRAPRVKS 257
S.enterica      ---------------------------------------------------------------------------------------------------------------------------------- 17

                       270       280       290       300       310       320       330       340       350       360       370       380       390
                ....|....|....|....|....|....|....|....|....|....|....|....|....|....|....|....|....|....|....|....|....|....|....|....|....|....|
AgoDN3          ---------------------------------------------------------------------------------------------------------------------------------- 1
RamDN3          ---------------------------------------------------------------------------------------------------------------------------------- 1
NgrDN3          ---------------------------------------------------------------------------------------------------------------------------------- 1
NfoDN3          ---------------------------------------------------------------------------------------------------------------------------------- 1
PcoAEDN3        ---------------------------------------------------------------------------------------------------------------------------------- 1
PcoWSDN3        ---------------------------------------------------------------------------------------------------------------------------------- 1
NdaDN3          ---------------------------------------------------------------------------------------------------------------------------------- 1
PkiDN3          ---------------------------------------------------------------------------------------------------------------------------------- 1
MjaDN3          ---------------------------------------------------------------------------------------------------------------------------------- 1
GokDN3          ---------------------------------------------------------------------------------------------------------------------------------- 1
E.coli          -----------------------------VPMENDVLKVV----------------------------KSSAAKVEPLPLVG----EGSDNYAGDEMVTKV----------------------------- 147
V.cholerae      -----------------------------VEMDNGVLKVI----------------------------KSKDAKTSAIPVL-----SGEERANGDEVITQV----------------------------- 128
A.salmonicida   -----------------------------VPMDNGVLKVV----------------------------RSKDAKTSAIPVV-----DETNPGIGDEMVTRV----------------------------- 129
P.aeruginosa    -----------------------------VAQGDQA-RIV----------------------------PNAEAKTEA----------GGGQSAPDRLETRV----------------------------- 148
N.meningtitis   ----------------------------------------A----KQQAAASAKQQ------------AAAPAKQQAAAPAKQTNIDFRKDGKNAGIIELAALGFAGQPDISQQHDHIIVTLKNHTLPTT 264
M.xanthus       GALRDVRVGAHADKVRLVLDVRGTMPAYRVDRANRGLEVVLGRAVARKAAPAVETQAVVASVAEVEPLRQAPVKSEASPVVEVKDVRFEESSAGGRIVMKLSGTSAWKVDRPDPRSAVL-TLENARLPKK 386
S.enterica      ---------------------------------------------------------------------------------------------------------------------------------- 17

                       400       410       420       430       440       450       460       470       480       490       500       510       520
                ....|....|....|....|....|....|....|....|....|....|....|....|....|....|....|....|....|....|....|....|....|....|....|....|....|....|
AgoDN3          ----------------------------------------------------------------------------------------------------------------MSNRNELVDRVVRHR--R 16
RamDN3          -------------------------------------------------------------------------------------------------------------MSRHLERVSYAPGGG------ 15
NgrDN3          -----------------------------------------------------------------------------------------------------------------MNK------LNFR--SN 9
NfoDN3          -----------------------------------------------------------------------------------------------------------------MNKAEIRKPLSFRNASQ 17
PcoAEDN3        ---------------------------------------------------------------------------------------------------------------------------------- 1
PcoWSDN3        ---------------------------------------------------------------------------------------------------------------------------------- 1
NdaDN3          -----------------------------------------------------------------------------------------------------------------MNEEEN-EVLSFR--QK 14
PkiDN3          --------------------------------------------------------------------------------------------------------------MTDTPNNNNSEKLEFR--PM 18
MjaDN3          --------------------------------------------------------------------------------------------------------------MDRPARRDNTDGSRSEHPQG 20
GokDN3          ----------------------------------------------------------------------------------------------------------------MSALPSLADGSR------ 12
E.coli          -----------VPVRNVSVRELAPI---LRQMIDSAGSGN-VVNYDP-----------------------------------------------SNVIMLTGRASVVERLTEVIQRV---------DHAG 206
V.cholerae      -----------VAVKNVSVRELSPL---LRQLIDNAGAGN-VVHYDP-----------------------------------------------ANIILITGRAAVVNRLAEIIRRV---------DQAG 187
A.salmonicida   -----------VPVRNVSVRELAPL---LRQLNDNAGGGN-VVHYDP-----------------------------------------------SNVLLITGRAAVVNRLVEVVRRV---------DKAG 188
P.aeruginosa    -----------IQVQQSPVSELIPL---IRPLVPQ--YGH-LAAVPS-----------------------------------------------ANALIISDRSANIARIEDVIRQL---------DQKG 205
N.meningtitis   LQRSLDVADFKTPVQKVTLKRLNND---TQLIITTAGNWE-LVNKSAAPGYFTFQVLPKKQNLESGGVNNAPKTF------------------TGRKISLDFQDVEIRTILQILAKESGMNIVASDSVNG 372
M.xanthus       FERSLDTSALDTPVKMISAFSVPGEGGKVRLVVAADGAIEEKVSQSA--GTLSWRLDVKGVKTEEVAVAQRTAGFTTEAPAYAAEGAPQQARYRGKRVSFEFKDIDIQNLLRVIAEISKKNIVVADDVSG 514
S.enterica      -----------------------------------------YVSGPP-----------------------------------------------VYVDMVVNAATMMDKQNDGIE-------------LG 46

                       530       540       550       560       570       580       590       600       610       620       630       640       650
                ....|....|....|....|....|....|....|....|....|....|....|....|....|....|....|....|....|....|....|....|....|....|....|....|....|....|
AgoDN3          AAPFIPGRPRFSVTDIAEPVVHRDD--------------------EMIRYYARKMKPGSLDARAPTIKE---------EPMKM--------YFYYPKYVEPENAAQV-----------ELLRTWEENDGI 98
RamDN3          ------TGVYITAEDMLRAEAEEDS----SHTPSSNQQ---------QVVNIVPRDPLAPSREEREAER-----------PQM--------FFYYPKYLDVDAVTLI---------AQNLLDTWEKEE-- 96
NgrDN3          ISDYSLE------PEENNSSTSNKT----NNNDITNEEE------SSSE---KALKIEENIRKEKERN----------EPYQM--------FFYYPRYLHPEAVSLI---------AQNLLNQWEDEEVS 93
NfoDN3          LLDTSEDKVVNETPLDNAHEGSDRTLLSATGTDQTTLDSSFESSLSSSDQISDELLLAEQIRREKERN----------EPYQM--------FFYYPKYLHPEAVALI---------AQNLLNEWEEE--- 117
PcoAEDN3        -------------------------------------------------LRAAAQEAENEKIRGEKRE--------------L--------FFYYPKYLHPEAVTTI---------LQNLLNDWE----- 45
PcoWSDN3        ----------------------------------------------------RGIEAIN-ILQNETRQ--------------I--------FFYKPKYLHPEALKMI---------AENLLAEWEREI-- 44
NdaDN3          APNFINIE-------ESEPQQSTKP--------------------------MEEIKKEETKETKPLKD----------EPMKM--------FFYYPKYLHPEAVALI---------SQNLLNEWEEK--- 81
PkiDN3          SADFISEESLVENKDNHQLQQKEQKQQNQSNSNSHYNQEKLQKHIEASQKAAEQRKEKNQTYLNNNNNNNNSDSDNLNDSMHM--------YFYYPKYLDVESVSVI---------AQNLLSEWEEE--- 128
MjaDN3          RVSFRQPAPTFITPIQLKEG-HGRR----SEVPATDLP------SAPSSAPLPPLPPVETQQSEGPEE------------RQM--------FFYYPKYLDPDAVVLISRTLLQQWEASE----------- 108
GokDN3          -VTFRSPAPVVVTSDTLRTDPQQPR----HHLRTDDLDPTLFLKSPPAIEDGQDAPPVDLNSTDAEKES-----------KEM--------FFYYPKYLDPDAVVLIARTLLKNWEEHP----------- 107
E.coli          NRTEEVIPLDNASASEIARVLESLTKNS---GENQPATLKSQIVADERTNSVIVSGDPATRDKMRRLIRRLDSEMERSGNS----------QVFYLKYSKAEDLVDVLKQVSGTLTAAKEEAEGTVGSGR 323
V.cholerae      DKEIEVVELNNASAAEMVRIVEALNKTTD--AQNTPEFLKPKFVADERTNSILISGDPKVRERLKRLIKQLDVEMAAKGNN----------RVVYLKYAKAEDLVEVLKGVSENLQAEKGTGQPTT-SKR 304
A.salmonicida   DQEVDIIKLKYASAGEMVRLVTNLNKDGNTQGGNTSLLLAPKVVADERTNSVVVSGEPKARARIIQMVRQLDRDLQSQGNT----------RVFYLKYGKAKDMVEVLKGVSTSIEADKKGGGATAGGGN 308
P.aeruginosa    SHDYSVINLRYGWVMDAAEVLNNAMSRGQAKGA-----AGAQVIADARTNRLIILGPPQARAKLVQLAQSLDTPTARSANT----------RVIRLRHNDAKTLAETLGQISEGMKNNGGQGGEQTGGGR 320
N.meningtitis   KMTLSLKDVPWDQALDLV--------------------MQARNLDMRQQGNIVNIAP------RDELLAKDKAFLQAEKDIA-D-LGALYSQNFQLKYKNVEEFRSILRLDNADTTGNRN----TLVSGR 470
M.xanthus       KVTIRLRNVPWDQALDLV--------------------LRTKALGKEEFGNIIRIAPLKTLEEEARLRQERKASLQRQEDLMV--------NLIPVNYAVAADMAARVK---------------DVLSER 601
S.enterica      RQKIGVMRLNNTFVGDRT--------------------------YNLRDQKMVIPGIATAIERLLQGEEQPLGNIVSSEPPAMPAFSANGEKGKAANYAGGMSLQEALKQ--------------NAAAGN 136

                       660       670       680       690       700       710       720       730       740       750       760       770       780
                ....|....|....|....|....|....|....|....|....|....|....|....|....|....|....|....|....|....|....|....|....|....|....|....|....|....|
AgoDN3          RLASGASSPPSSQPQPQPQPQSEQEHERDEVVEVPSIAANDASKKKTIAPVSGEKVVFKRMVP------------------------------------------------------------------- 161
RamDN3          ---------QEQETEERRRQRTVQPPPL------------------------------------------------------------------------------------------------------ 115
NgrDN3          NDDNSNNNNQENTTSRPQK--------------------------------------------------------------------------------------------------------------- 112
NfoDN3          ----------ENK--------------------------------------------------------------------------------------------------------------------- 120
PcoAEDN3        ----------EKIEPQN---DKTNE----------EEAEKGKTLG----K-------------------------------------------------------------------------------- 68
PcoWSDN3        -----GIGMEEEVADENESVDGSNKKGDGKDAGQGDAAPKGRTLGGDSLKKTSGDRNHIAKR-------------------------------------------------------------------- 101
NdaDN3          ----------EDTK-------------------------------------------------------------------------------------------------------------------- 85
PkiDN3          ----------ER---------------------------------------------------------------------------------------------------------------------- 130
MjaDN3          SARKESNDGDAGKQ-------------------------------------------------------------------------------------------------------------------- 122
GokDN3          DVPVTMPTSLAAMIPGAPTAAPAPAPSQPQP--------------------------------------------------------------------------------------------------- 138
E.coli          -EVV-----SIAASKHSNALIVTAPQDIMQSLQSVIEQLDIRRAQVHVEALIVEVAEGSNINFGVQWGSK-----DAG----LMQFANGTQIPIGTLGAAISAAKPQKGSTVISENGATTINPDTN---G 435
V.cholerae      -NEV-----MIAAHADTNSLVLTAPQDIMNAMLEVIGQLDIRRAQVLIEALIVEMAEGDGINLGVQWGSL-----ESG---SVIQYGN-TGASIGNVMIGLEEAKDTTQTKAVYDTNNNFLRNETTTTKG 419
A.salmonicida   -ASIGGGKLAISADETTNALVITAQPDVMAELEQVVAKLDIRRAQVLVEAIIVEIADGDGLNLGVQWANT-----NGG----GTQFTN-AGPGIGSVAIAAKDYKDN----------------------G 405
P.aeruginosa    PSNI-----LIRADESTNALVLLADPDTVNALEDIVRQLDVPRAQVLVEAAIVEISGDIQDAVGVQWAIN-----KGGMGGTKTNFAN-TGLSIGTLLQSLESNKAP----------------------- 416
N.meningtitis   ------G--SVLIDPATNTLIVTDTRSVIEKFRKLIDELDVPAQQVMIEARIVEAADGFSRDLGVKFG-------ATG----KKKLKN-DTSAFGWGVNSGFGGDDK--------WGAETKINLPI---- 568
M.xanthus       ------G--SVTVDQRTNVLIVKDVRSNTERARSLVRSLDTQTPQVLIESRIVEANTSFSRSLGVQWGGQARAGQATG-NSTGLIFPN-NLAVTGGVTGTGAGLPDN----------PNFAVNLPT---G 708
S.enterica      ---I-----KIVAYPDTNSLLVKGTAEQVHFIEMLVKALDVAKRHVELSLWIVDLNKSDLERLGTSWSGSITIGDKLG---------------------------------------------------- 206
```

```
                      790       800       810       820       830       840       850       860       870       880       890       900       910
               ....|....|....|....|....|....|....|....|....|....|....|....|....|....|....|....|....|....|....|....|....|....|....|....|....|....|
AgoDN3         ---------------------------------------------------------------------------------------------------------------------------------- 161
RamDN3         ---------------------------------------------------------------------------------------------------------------------------------- 115
NgrDN3         ---------------------------------------------------------------------------------------------------------------------------------- 112
NfoDN3         ---------------------------------------------------------------------------------------------------------------------------------- 120
PcoAEDN3       ---------------------------------------------------------------------------------------------------------------------------------- 68
PcoWSDN3       ---------------------------------------------------------------------------------------------------------------------------------- 101
NdaDN3         ---------------------------------------------------------------------------------------------------------------------------------- 85
PkiDN3         ---------------------------------------------------------------------------------------------------------------------------------- 130
MjaDN3         ---------------------------------------------------------------------------------------------------------------------------------- 122
GokDN3         ---------------------------------------------------------------------------------------------------------------------------------- 138
E.coli         DLSTLAQLLSGFSGTAVGVVKGDWMALVQAVKNDSSSNVLSTPSITTLDNQEAFFMVGQDVPVLTGS--TVGSNNSNPFNTVERKKVGIMLKVTPQINEGNAVQMVIEQEV-SKV---EGQT-SLD--VV 556
V.cholerae     DYTKLASALSSIQGAAVSIAMGDWTALINAVSNDSSSNILSSPSITVMDNGEASFIVGEEVPVITGS--TAGSNNDNPFQTVDRKEVGIKLKVVPQINEGNSVQLNIEQEV-SNV---LGANGAVD--VR 541
A.salmonicida  TTTGIAKLAESFNGMAAGFYQGNWAMLVTALSTNTKSDILSTPSIVTMDNKEASFNVGQEVPVQTGK--QNSTSGDTTFSTIERKTVGTKLVVTPQINEGDSVLLTIEQEV-SSVGKQASGTEGLG--PT 530
P.aeruginosa   --------ESIPDGAIVGIGSSSFGALVTALSANTKSNLLSTPSLLTLDNQKAEILVGQNVPFQTGSYTTNSEGSSNPFTTVERKDIGVSLKVTPHINDGAALRLEIEQEI-SALLPNAQQRNNTD--LI 535
N.meningtitis  ----TAAANSISLVRAIS--SGALNLELSASESLSKTKTLANPRVLTQNRKEAKIESGYEIPFTVTSIANGGSSTNTEL-----KKAVLGLTVTPNITPDGQIIMTVKITK-DSPAQCASGN-QTI--LC 683
M.xanthus      TGQGVGGAMGFTFGSAGG--ALQLNLRLSAAENEGSVKTISAPKVTTLDNNTARISQGVSIPFSQTS----AQGVNTTF-----VEARLSLEVTPHITQDGSVLMSINASN-NQPDPSSTGA-NGQ--PS 823
S.enterica     --------VSLNQSSISTLDGSRFIAAVNALEEKKQATVVSRPVLLTQENVPAIFDNNRTFYTKLIG------ERNVAL---EHVTYGTMIRVLPRFSADGQIEMSLDIEDGNDKTPQSDTTTSVDALPE 319

                      920       930       940       950       960       970       980       990      1000      1010      1020      1030      1040
               ....|....|....|....|....|....|....|....|....|....|....|....|....|....|....|....|....|....|....|....|....|....|....|....|....|....|
AgoDN3         ---------------------------------------------------------------------------------------------------------------------------------- 161
RamDN3         ---------------------------------------------------------------------------------------------------------------------------------- 115
NgrDN3         ---------------------------------------------------------------------------------------------------------------------------------- 112
NfoDN3         ---------------------------------------------------------------------------------------------------------------------------------- 120
PcoAEDN3       ---------------------------------------------------------------------------------------------------------------------------------- 68
PcoWSDN3       ---------------------------------------------------------------------------------------------------------------------------------- 101
NdaDN3         ---------------------------------------------------------------------------------------------------------------------------------- 85
PkiDN3         ---------------------------------------------------------------------------------------------------------------------------------- 130
MjaDN3         ---------------------------------------------------------------------------------------------------------------------------------- 122
GokDN3         ---------------------------------------------------------------------------------------------------------------------------------- 138
E.coli         FGERKLKTTVLANDGELIVLGGLMDDQAGESVAKVPLLGDIPLIGNLFKSTADKKEKRNLMVFIRPTILRDGMAADGVSQRKYNYMRAEQIYRDEQ-GLSLMPHTAQPILP--AQNQALPPEVRAFLNAG 683
V.cholerae     FAKRQLNTSVMVQDGQMLVLGGLIDERALESESKVPLLGDIPLLGQLFRSTSSQVEKKNLMVFIKPTIIRDGVTADGITQRKYNYIRAEQLFRAEK-GLRLLDDASVPVLPKFGDDRRHSPEIQAFIEQM 670
A.salmonicida  FDTRTVKNAVLVKSGETVVLGGLMDEQTKEEVSKVPLLGDIPVLGYLFRSTSNTTSKRNLMVFIRPTILRDANVYSGISSNKYTLFRAQQLEAAAQKGYATSPDRQ--VLPEYGQDVVQSPEIQKQIELM 658
P.aeruginosa   TSKRSIKSTILAENGQVIVIGGLIQDDVSQAESKVPLLGDIPLLGRLFRSTKDTHTKRNLMVFLRPTVVRDSAGLAALSGKKYSDIRVIDGTRGPE-GRPS-------ILPTNANQLFDGQAVDLR-ELM 656
N.meningtitis  ISTKNLNTQAMVENGGTLIVGGIYEEDNGNTLTKVPLLGDIPVIGNLFKTRGKKTDRRELLIFITPRIMGTAG----------NSLRY------------------------------------------ 761
M.xanthus      IQRKEANTQVLVKDGDTTVIGGIYVRRGATQVNSVPFLSRIPVLGLLFKNNSETDTRQELLIFITPRILNRQTIAQTL---------------------------------------------------- 901
S.enterica     VGRTLISTIARVPHGKSLLVGGYTRDANTDTVQSIPFLGKLPLIGSLFRYSSKNKSNVVRVFMIEPKEIVDPLTPDA-SESVNNILKQSGAWSGDD---------------------KLQKWVRVYLDRG 427

                     1050
               ....|....|....|....
AgoDN3         ------------------- 161
RamDN3         ------------------- 115
NgrDN3         ------------------- 112
NfoDN3         ------------------- 120
PcoAEDN3       ------------------- 68
PcoWSDN3       ------------------- 101
NdaDN3         ------------------- 85
PkiDN3         ------------------- 130
MjaDN3         ------------------- 122
GokDN3         ------------------- 138
E.coli         RTR---------------- 686
V.cholerae     EAKQ--------------- 674
A.salmonicida  KARQQATADGAQPFVQGNN 677
P.aeruginosa   TE----------------- 658
N.meningtitis  ------------------- 761
M.xanthus      ------------------- 901
S.enterica     QEAIK-------------- 432
```

**Supplementary Fig. 4A-D** Multiple sequence alignments of eukaryotic GspD, GspDN1, GspDN2 and GspDN3 with bacterial GspD proteins. For further details see page 23.

(E) GspE

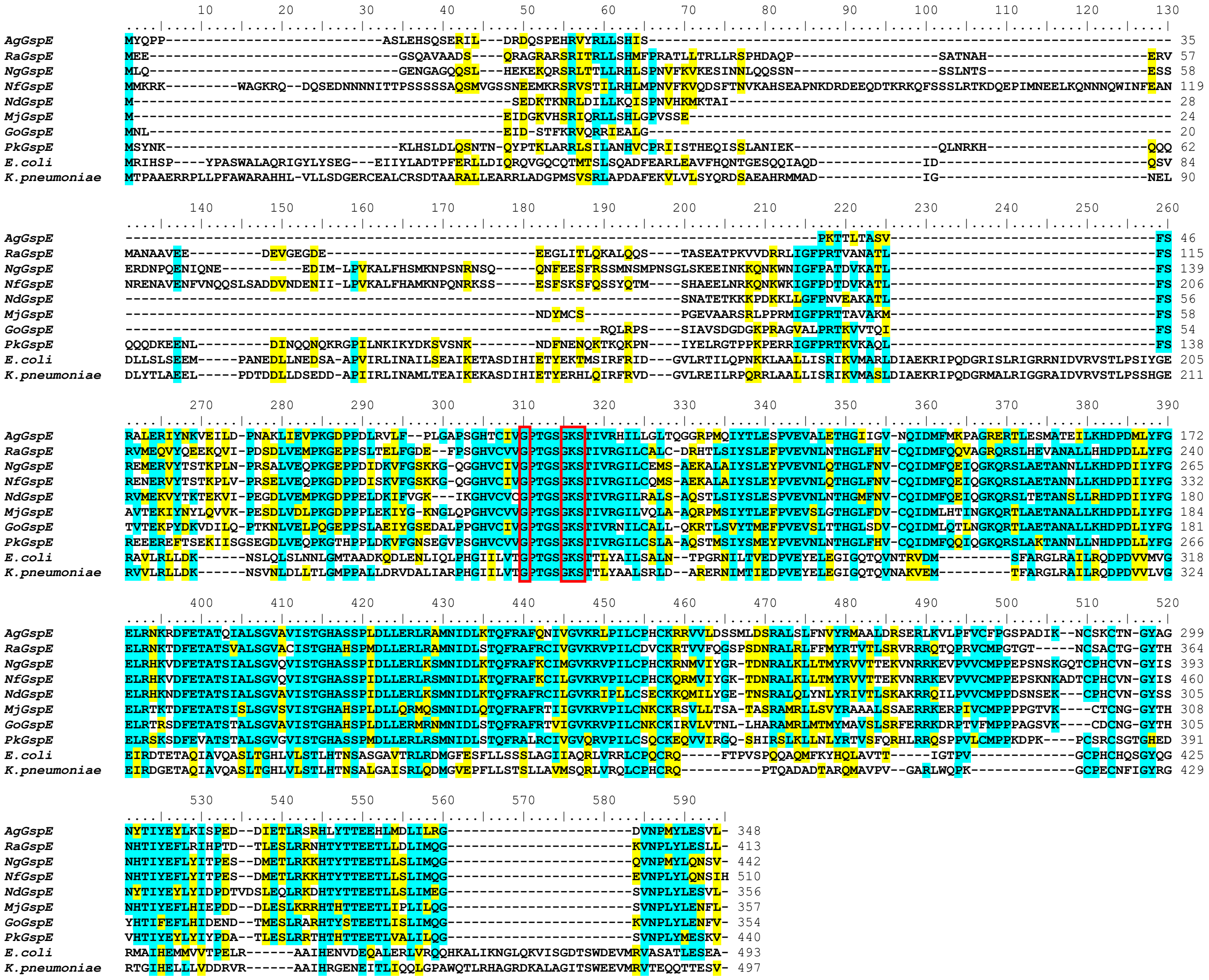
AgGspE
RaGspE
NgGspE
NfGspE
NdGspE
MjGspE
GoGspE
PkGspE
E.coli
K.pneumoniae

(F) GspEL

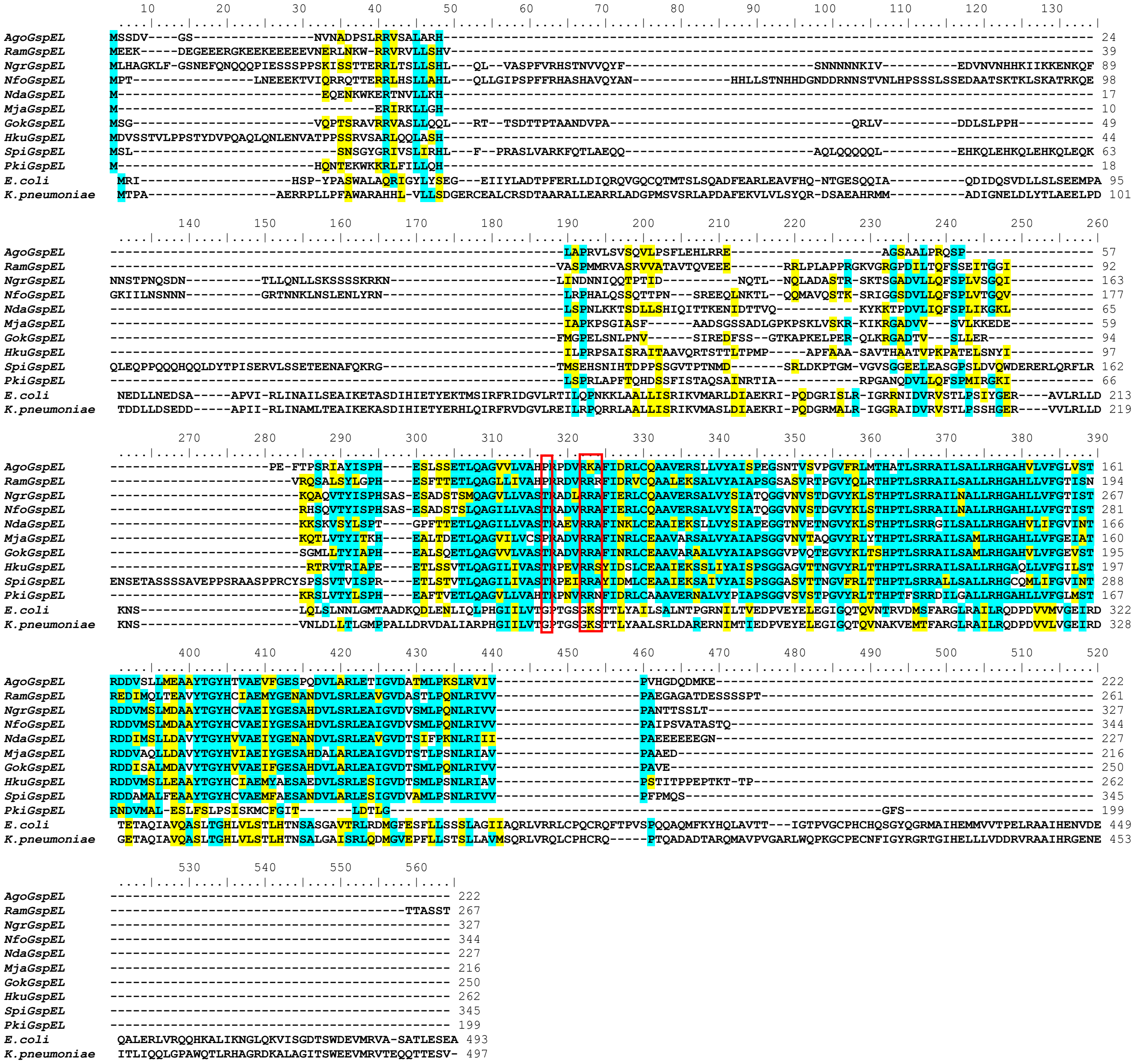
AgoGspEL
RamGspEL
NgrGspEL
NfoGspEL
NdaGspEL
MjaGspEL
GokGspEL
HkuGspEL
SpiGspEL
PkiGspEL
E.coli
K.pneumoniae

(G) GspEN2A

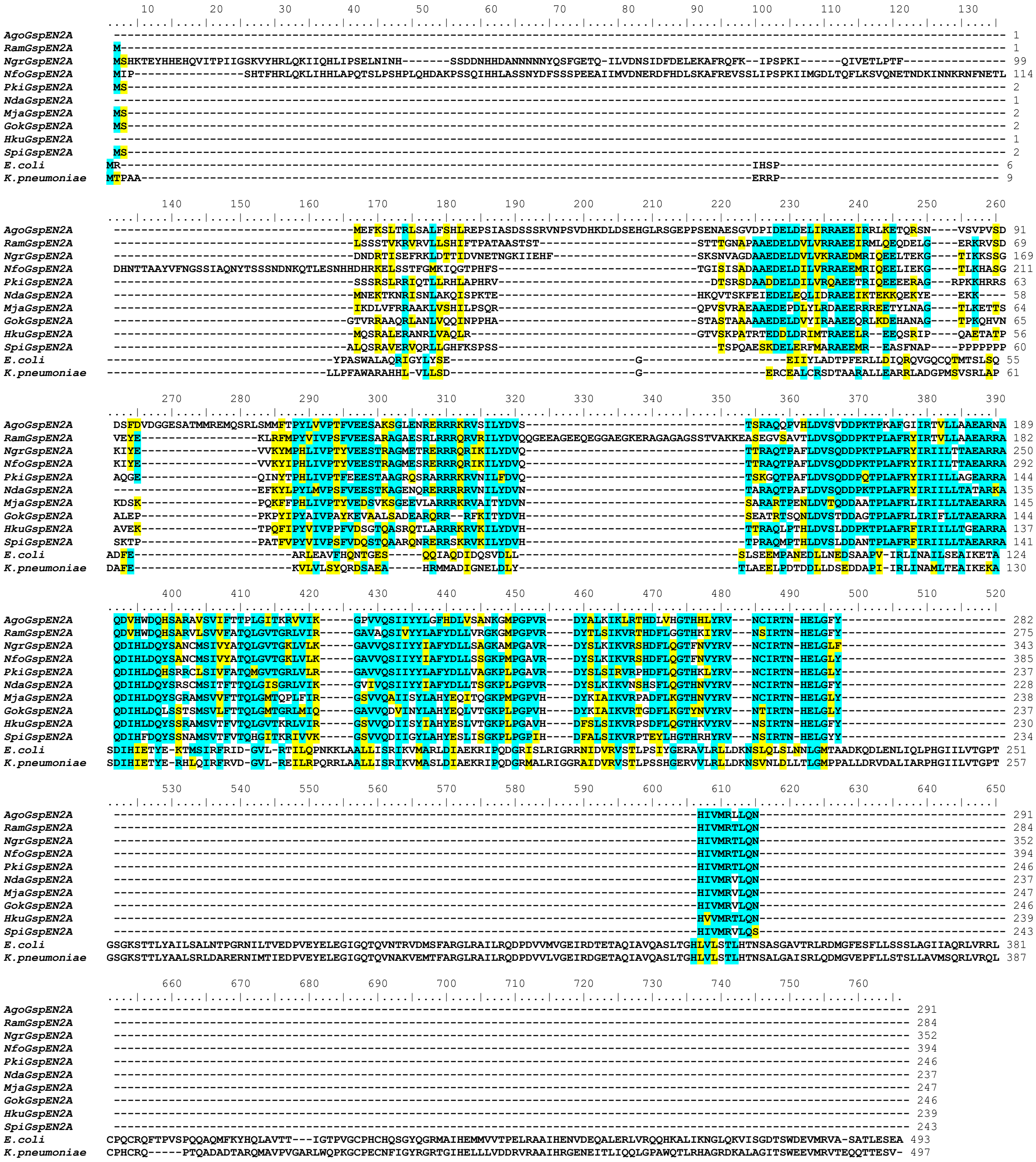

(H) GspEN2B

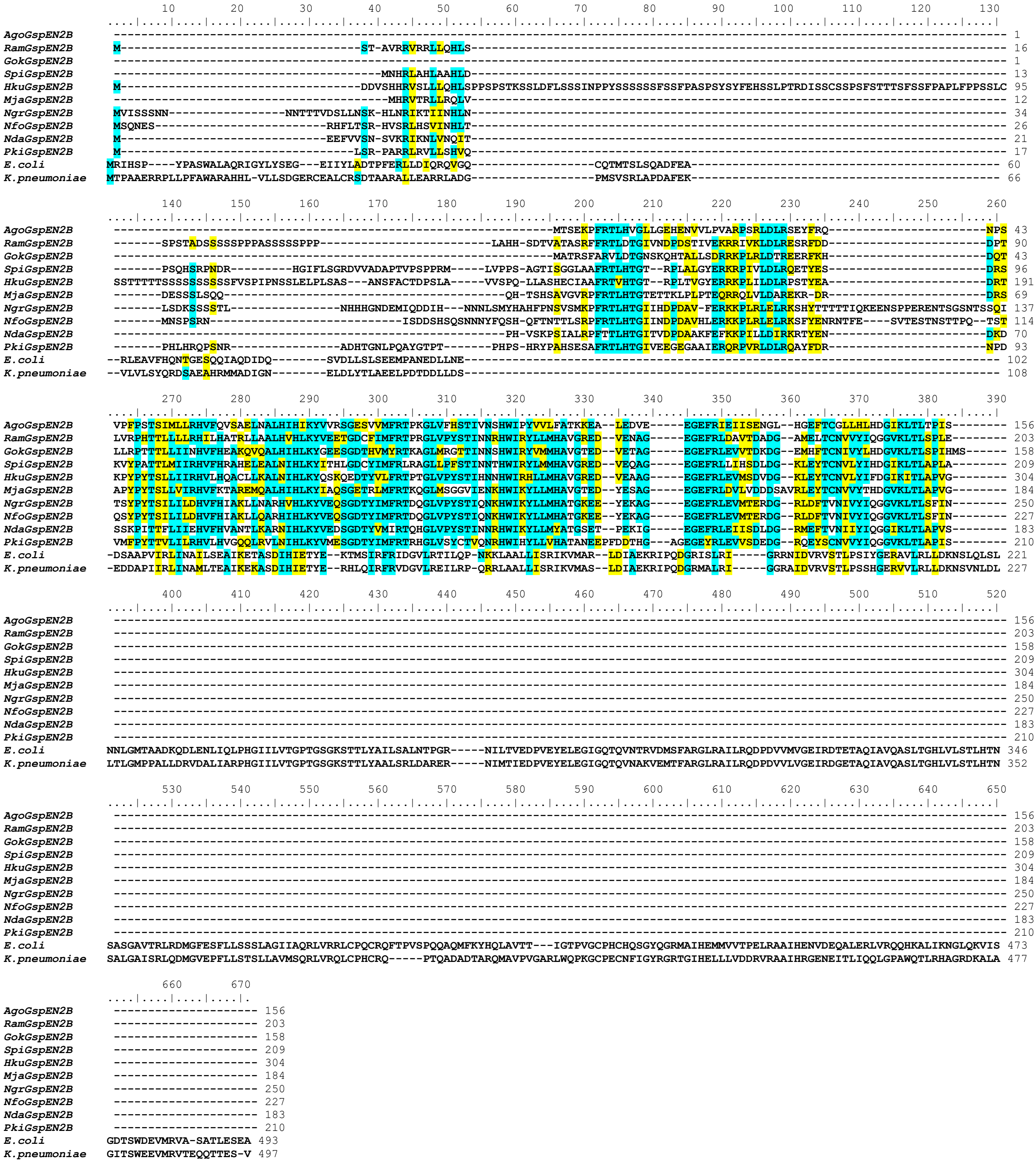

**Supplementary Fig. 4E-H** Protein sequence alignments of eukaryotic GspE, GspEL, GspEN2A and GspEN2B with reference bacterial GspE proteins. For further details see page 23.

(I) GspF

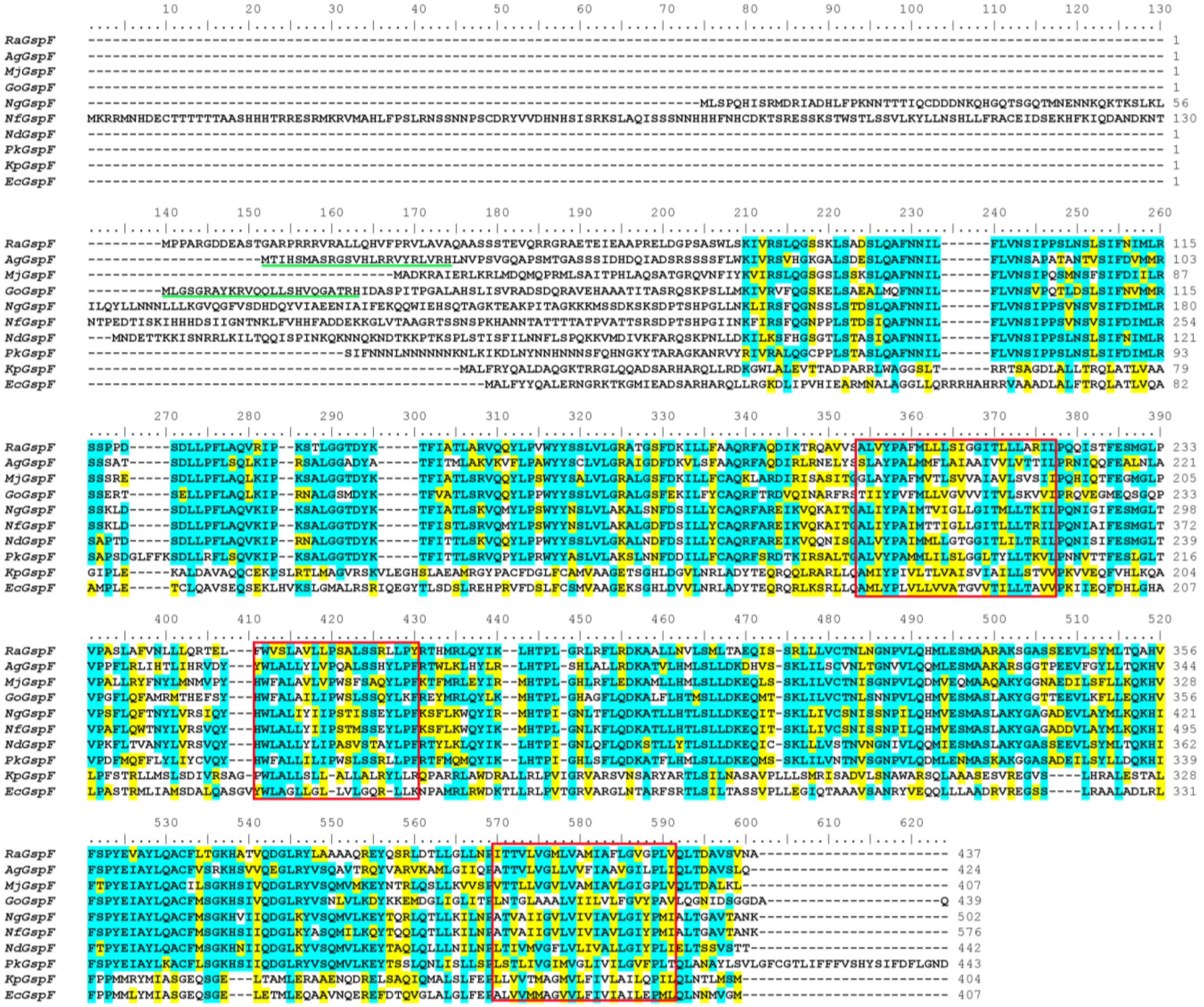

**Supplementary Fig. 4I** Protein sequence alignments of eukaryotic and reference bacterial GspF proteins. For further details see page 23.

(J) GspG

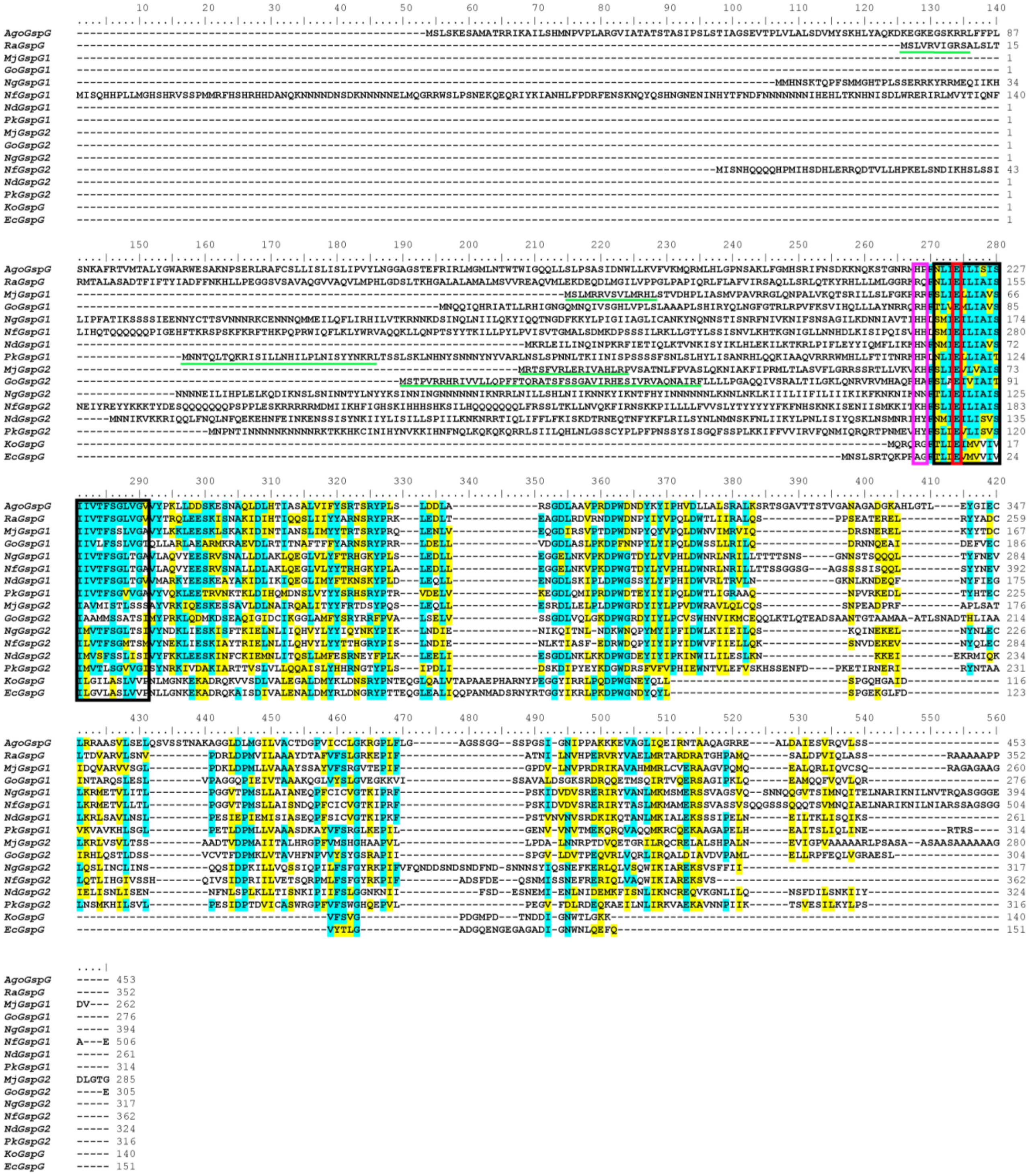


**Supplementary Fig. 4J** Protein sequence alignments of eukaryotic and reference bacterial GspG proteins. For further details see page 23.

**Supplementary Fig. 4** (pg. 8-22) Multiple sequence alignments of eukaryotic GspD (panel A-D, pg. 8-16), GspE (panel E-H, pg. 17-20), GspF (panel I, pg. 21) and GspG (panel J, pg. 22) proteins with their prokaryotic homologues. The alignments illustrate the presence of N-terminal sequence extensions in the eukaryotic proteins, with targeting sequences recognized by MitoFates predictor[2] underlined in green. Identical and similar residues are highlighted in turquoise and yellow, respectively (using 50% conservation of the position as the threshold). **A.** In addition to the initially identified eukaryotic GspD proteins, representative sequences of GspDL, a more divergent paralogue identified by phylogenetic profiling, are included in the alignment together with reference bacterial GspD sequences. **B-D.** Three eukaryotic proteins homologous to the N-domain of bacterial secretins. While GspDN1 corresponds to a full single N-domain, GspDN2 and GspDN3 relate to its C-terminal and the N-terminal halves.
**E-H.** In addition to GspE proteins, three additional eukaryotic proteins GspEL, GspEN2A and GspEN2B were identified. While the bacterial GspE proteins contain four domains referred to as N1E, N2E, C1E and C2E, all four eukaryotic GspE proteins (GspE, GspEL, GspN2A and GspN2B) contain different domain variants. GspE proteins (**E**) carry C1E and C2E domains, GspEL proteins (**F**) contain C1E domain and GspEN2A and GspEN2B (**G** and **H**) contain N-domains. Only eukaryotic GspE proteins carry GxxxGK[ST] motif (also called the Walker A motif or the P-loop) functionally critical for the ATPase function of the protein. **I.** Transmembrane domains of GspF are highlighted by red rectangles. **J.** A conserved polar anchor of GspG is highlighted by the magenta rectangle and the transmembrane domain by a black rectangle, while the absolutely conserved glutamic acid residue at the +5 position relative to the pseudopilin processing site is highlighted in red.
Species abbreviations: *Mj – Malawimonas jakobiformis*, *Go – Gefionella okellyi*, *Ra – Reclinomonas americana*, *Ag – Andalucia godoyi*, *Nd – Neovahlkampfia damariscottae*, *Pk – Pharyngomonas kirbyi*, *Ng – Naegleria gruberi*, *Nf – Naegleria fowleri*, *Ko* **-** *Klebsiella oxytoca, Kp – Klebsiella pneumoniae, Ec – Escherichia coli.*

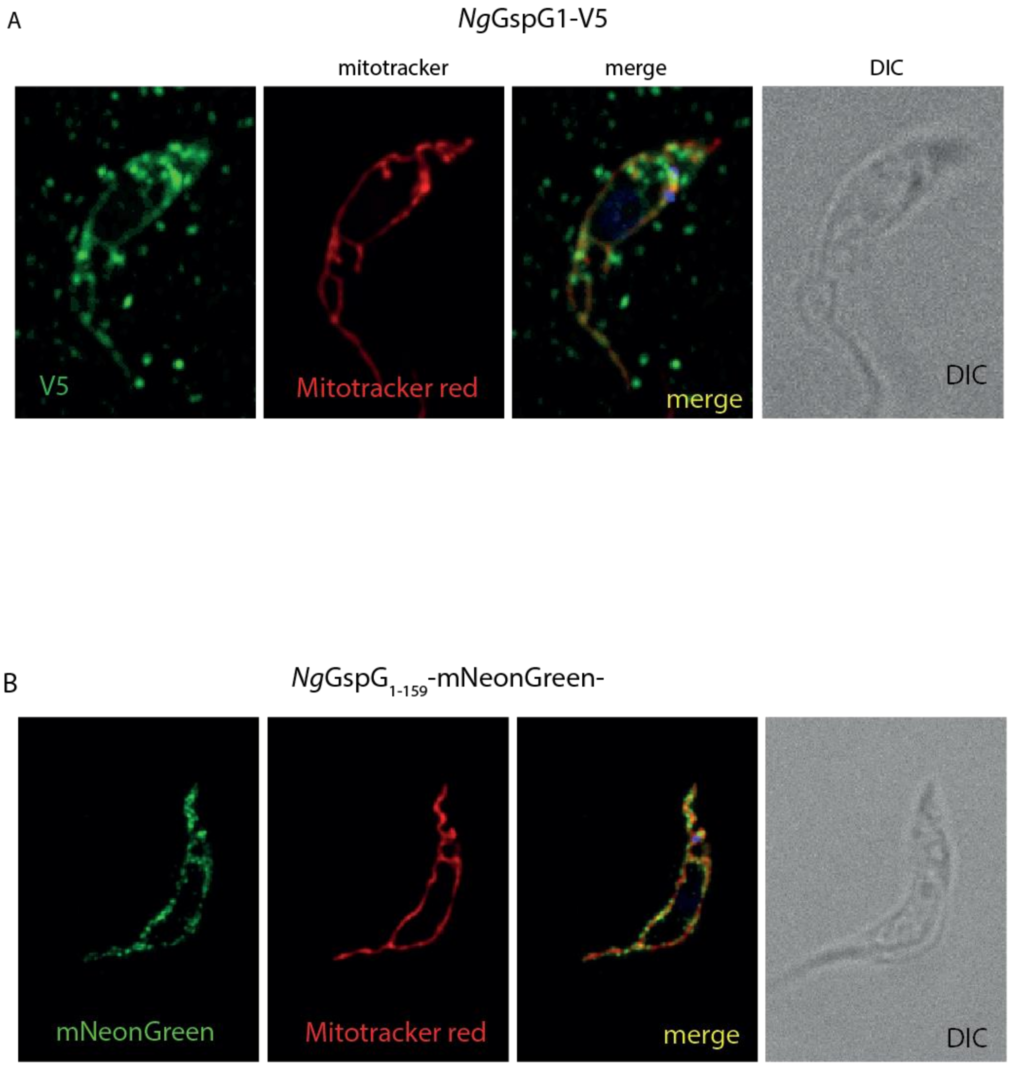


**Supplementary Figure 5. *Ng*GspG1 expressed in *T. brucei*.** The expression of *Ng*GspG1 (A) and the N-terminal part of the protein (residues 1-159) fused with mNeonGreen shows the mitochondrial localization of the constructs. In the case of full-length *Ng*GspG1 with the C-terminal V5 tag, only very weak expression upon high contrasting could be detected.

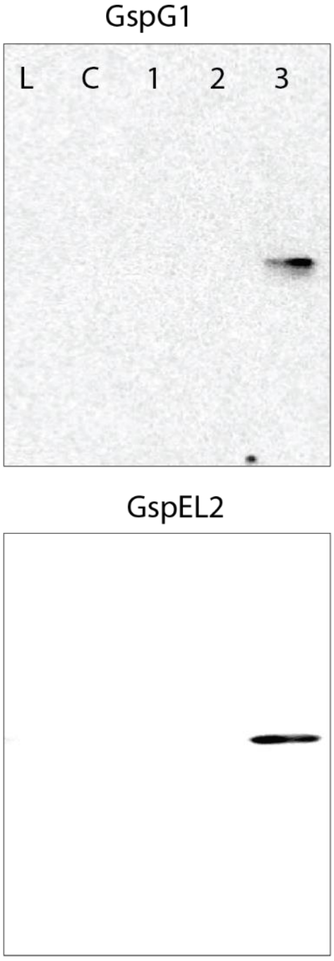


**Supplementary Fig. 6.** Example of western blot analysis using specific polyclonal antibody raised against GspG1and GspEL2 of *N. gruberi.* L - lysate, C – cytosol, 1,2,3 – sub-fraction of high-speed pellet fraction obtained by gradient centrifugation.

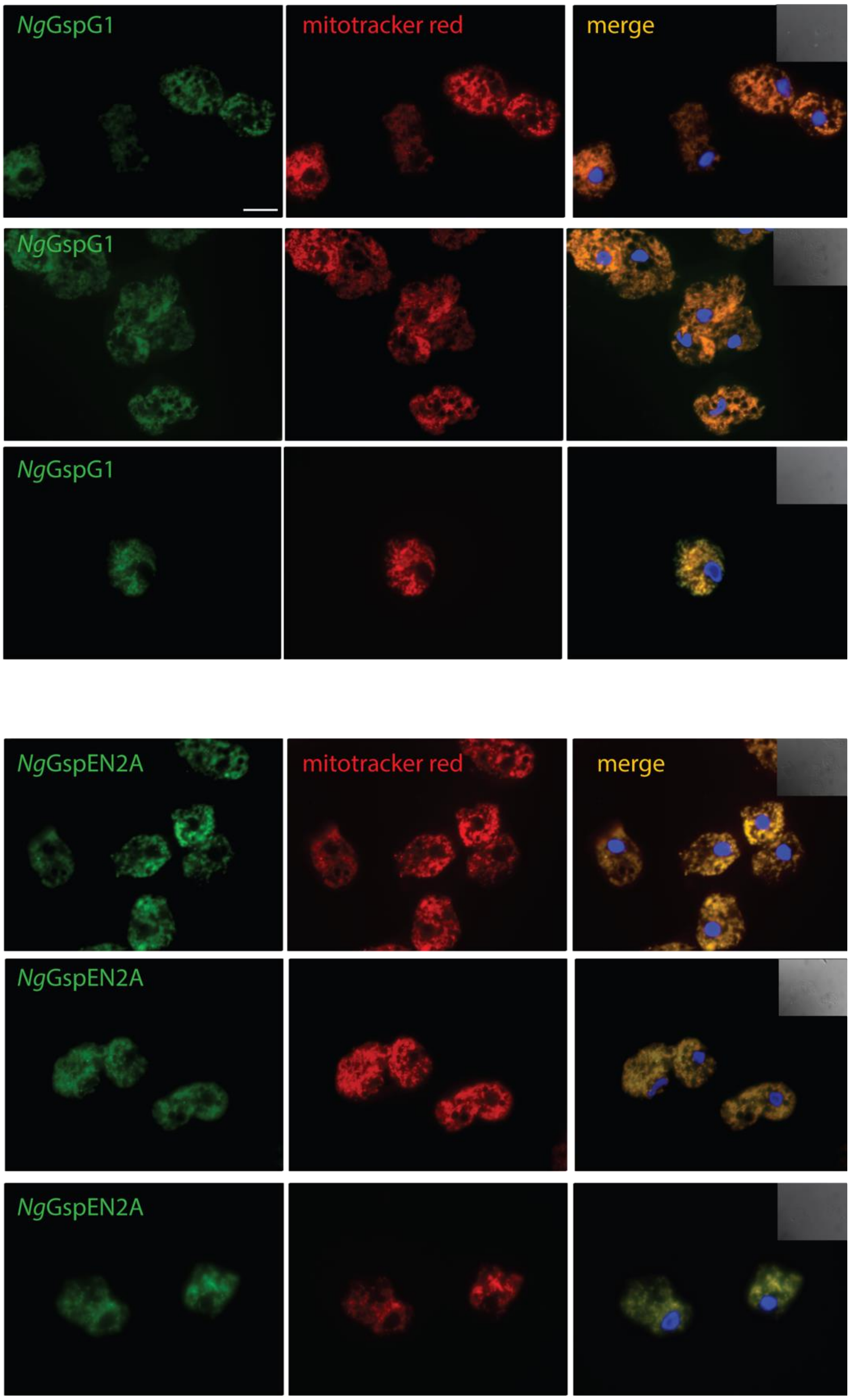


**Supplementary Fig. 7. Additional images of immunofluorescence microscopy detection of *Ng*GspG1 and *Ng*GspEL2.** Scale bar 10μm.

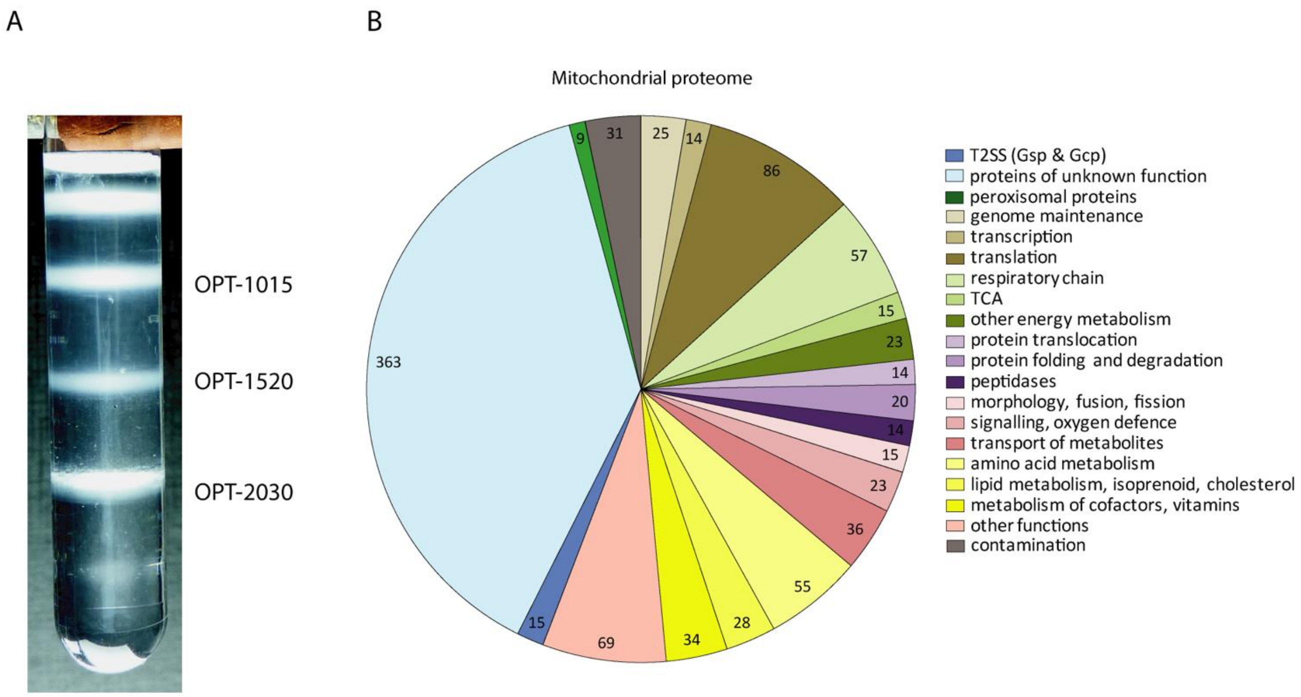


**Supplementary Fig. 8** Functional annotation of the putative mitochondrial proteome of *N. gruberi*. **A.** *N. gruberi* cell lysate was further separated into three fractions on Optiprep gradient and proteins extracted from these fractions were then digested with trypsin and peptides were separated by nanoflow liquid chromatography and analyzed by tandem mass spectrometry. **B.** In total 946 putative mitochondrial proteins were identified, which were sorted into functional categories using BLAST and HHpred. The number of proteins in each category is indicated within the respective segment of the pie chart.

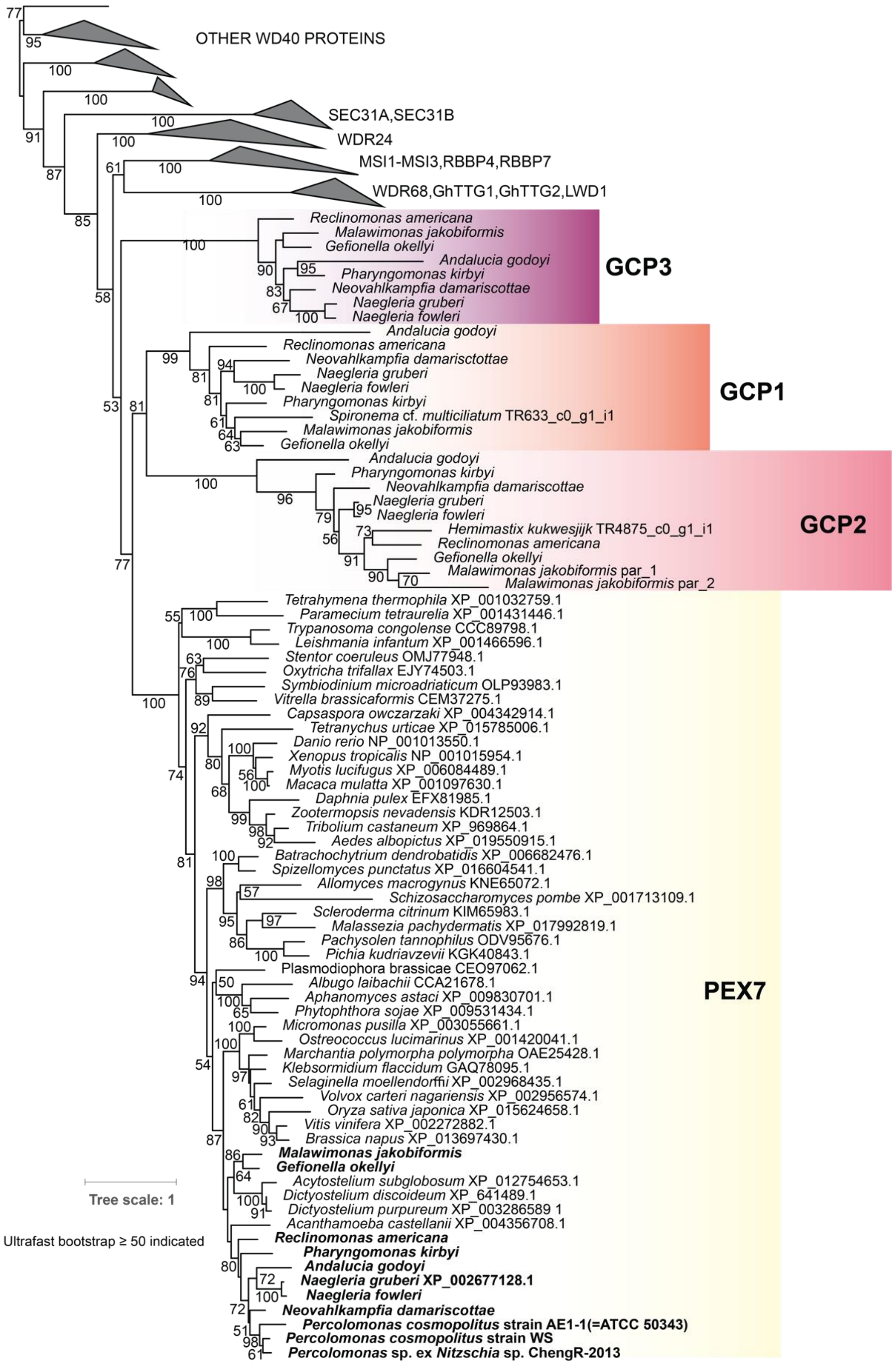


**Supplementary Fig. 9** Phylogenetic analysis of the WD40 superfamily including the novel members Gcp1, Gcp2, and Gcp3. The ML tree (IQ-TREE, LG+C60+G, 1000 ultrafast bootstraps, bnni) demonstrates the monophyly of each novel paralogue and suggests that they are specifically related to the peroxisome import protein Pex7. Note that the species possessing Gcp1 to Gcp3 contain also Pex7 itself (in bold).

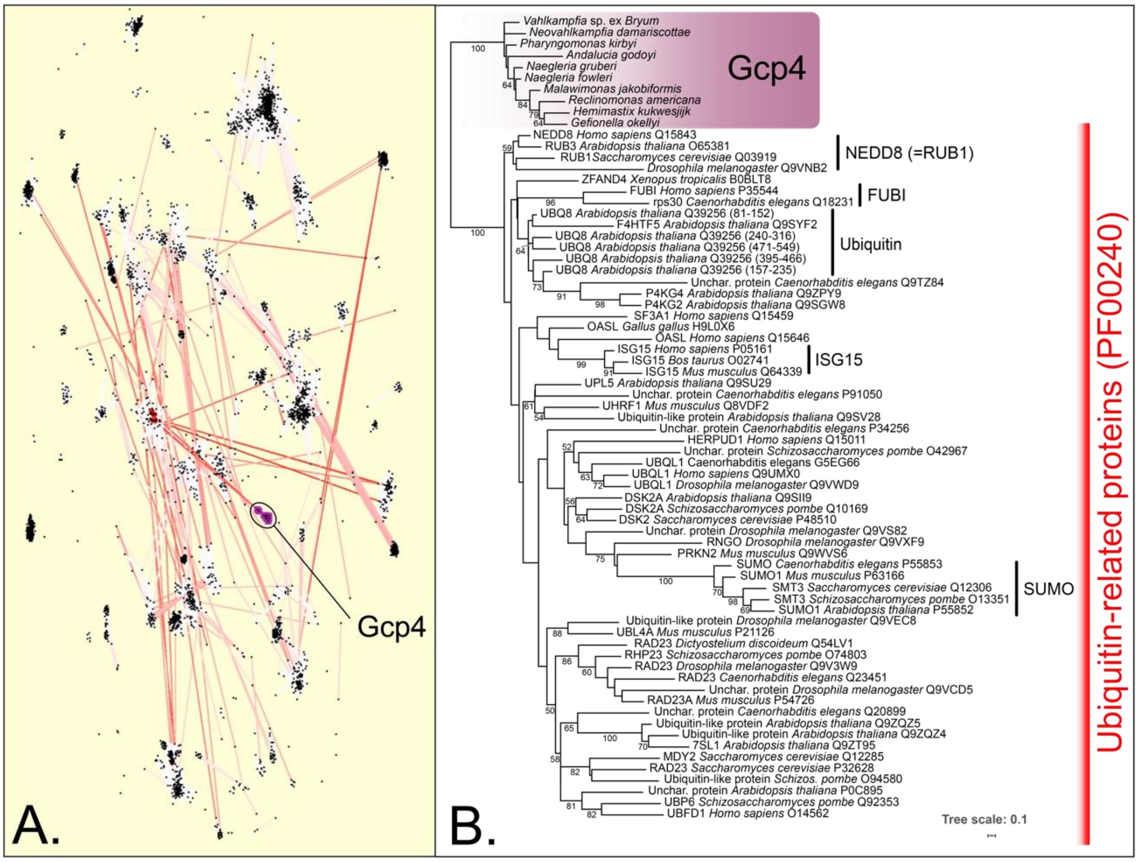


**Supplementary Fig. 10** Gcp4 proteins and their relationship to other members of the clan Ubiquitin (CL0072) as defined in Pfam v.31 database (https://pfam.xfam.org). **A.** Cluster analysis of ubiquitin fold of 4764 representative sequences (seed alignments) of the clan Ubiquitin containing members of all 59 defined families. This analysis was computed using the cluster analysis implemented in the CLANS (CLuster ANalysis of Sequences) program with following parameters: BLOSUM62 scoring matrix; extract BLAST HSPs up to E-value of 1e-4; 10,000 rounds. The analysis shows that Gcp4 sequences constitute a novel separate group with a weak affinity to the cloud containing proteins from ubiquitin family (PF00240) and Rad60 SUMO-like family (PF11976). Gcp4 proteins are marked in purple, four proteins directly connected to the Gcp4 cluster and situated in a cloud composed from ubiquitin and Rad60 SUMO-like families are marked in red (R5RDZ3_9PROT, NEDD8_HUMAN, RUB3_ARATH, A8BJ08_GIAIC). Connections between sequences are coloured by edge "frustration" (red=too long; blue=too short). **B.** Phylogenetic analysis of Gcp4 and ubiquitin-related proteins. Gcp4 sequences were added to the seed alignment of the ubiquitin domain as defined in the Pfam v.31 database (PF00240) and the ML tree was calculated using IQ-TREE multicore version 1.5.5 under the LG+G4 model with 10,000 ultrafast bootstraps. The tree is arbitrarily rooted between Gcp4 proteins and the remaining sequences included in the analysis. Sequences assigned to named clades correspond to different type I ubiquitin-like proteins (i.e. those that are conjugated to substrate proteins), the remaining sequences correspond to Type II ubiquitin-like proteins (i.e., larger proteins including a ubiquitin-like domain within a more complex domain architecture.

Gcp5

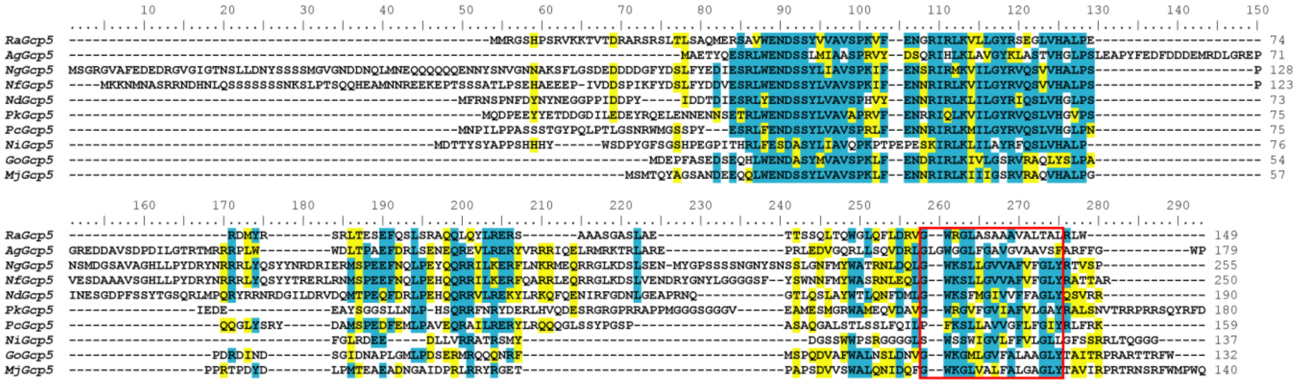

Gcp7

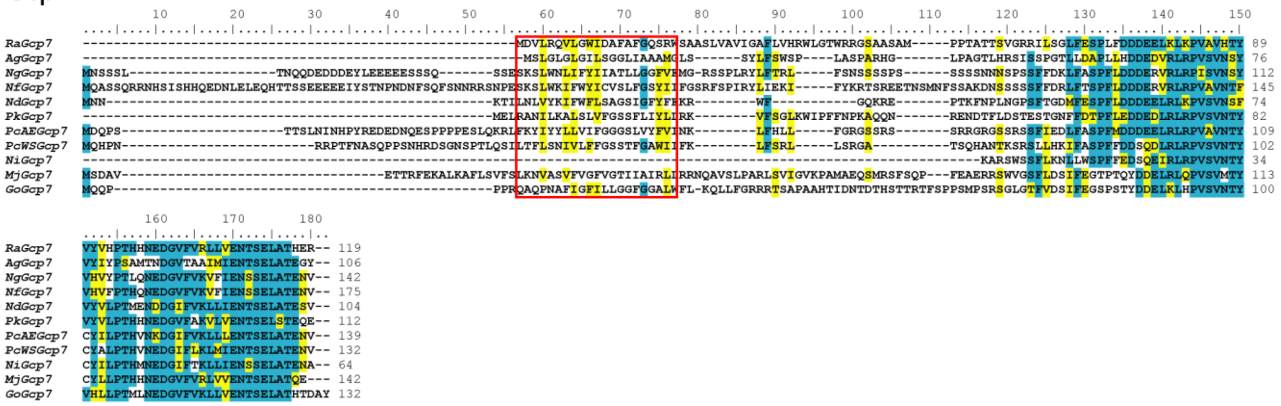

Gcp11

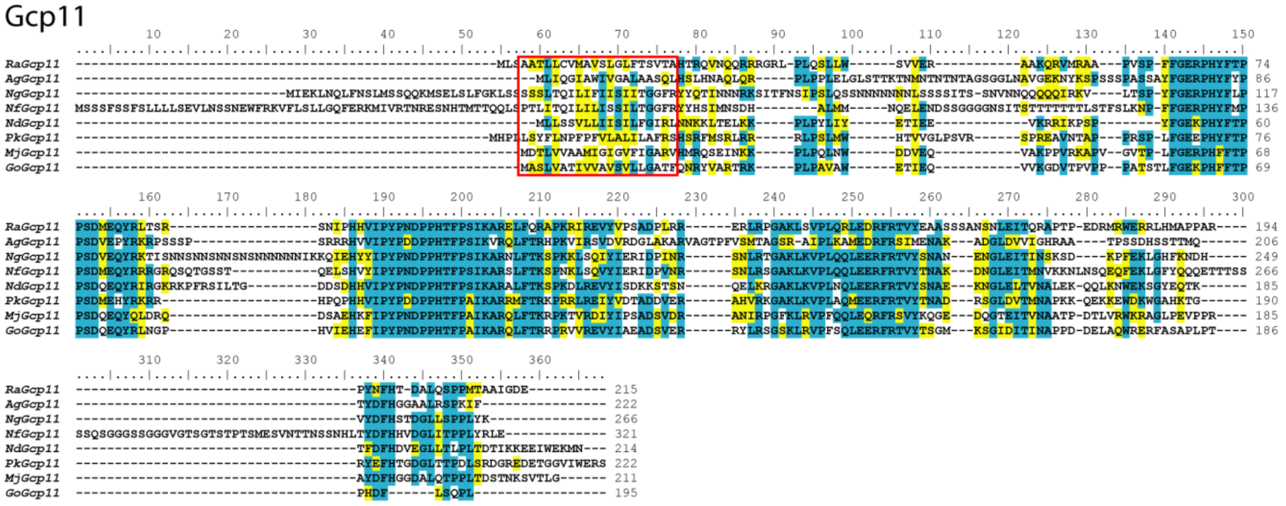

**Supplementary Fig. 11** (pg. 30 and 31) Multiple sequence alignments of Gcp5, Gcp7, Gcp11, and Gcp15 proteins. Regions including predicted transmembrane domains are framed by red rectangles. Identical and similar residues are highlighted in turquoise and yellow, respectively (using 50% conservation of the position as the threshold). Species abbreviations: *Mj – Malawimonas jakobiformis*, *Go – Gefionella okellyi*, *Ra – Reclinomonas americana*, *Ag – Andalucia godoyi*, *Nd – Neovahlkampfia damariscottae*, *Pk – Pharyngomonas kirbyi*, *Ng – Naegleria gruberi*, *Nf – Naegleria fowleri*, *PcWS – Percolomonas cosmopolitus* strain WS, *PcAE – Percolomonas cosmopolitus* strain AE-1, *Ni – Percolomonas* sp. contaminating a transcriptome assembly from *Nitzschia* sp. ChengR-2013. *Trit – Naegleria* sp. contaminating a transcriptome assembly from *Triticum polonicum*

Gcp15

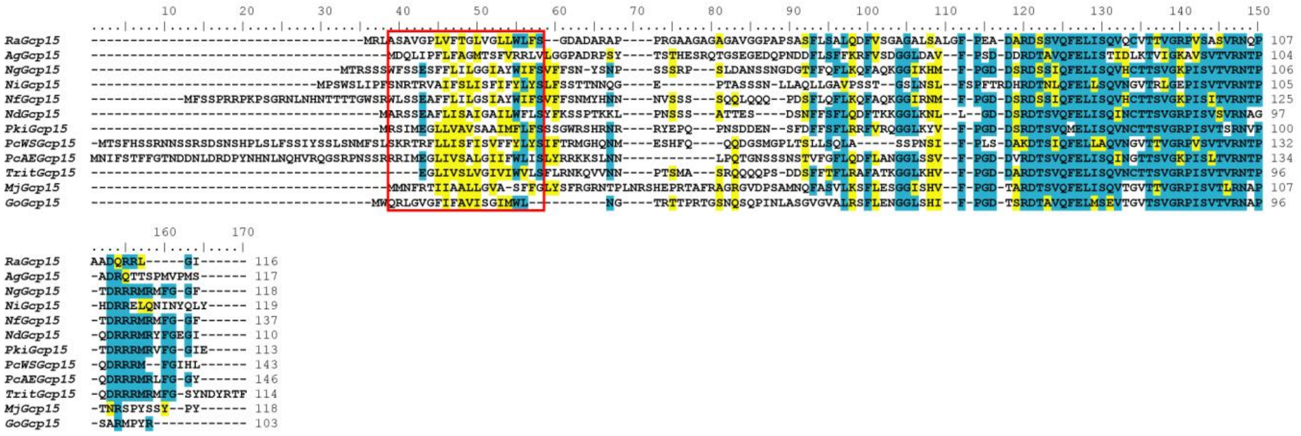


**Supplementary Fig. 11 cont.** For further details see page 30.

Gcp6

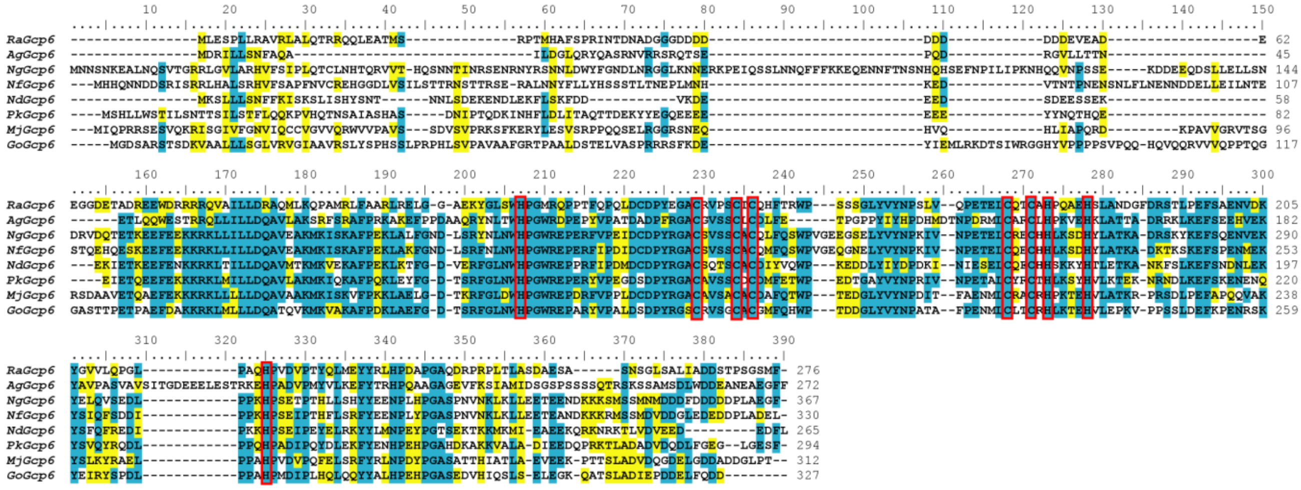

Gcp12

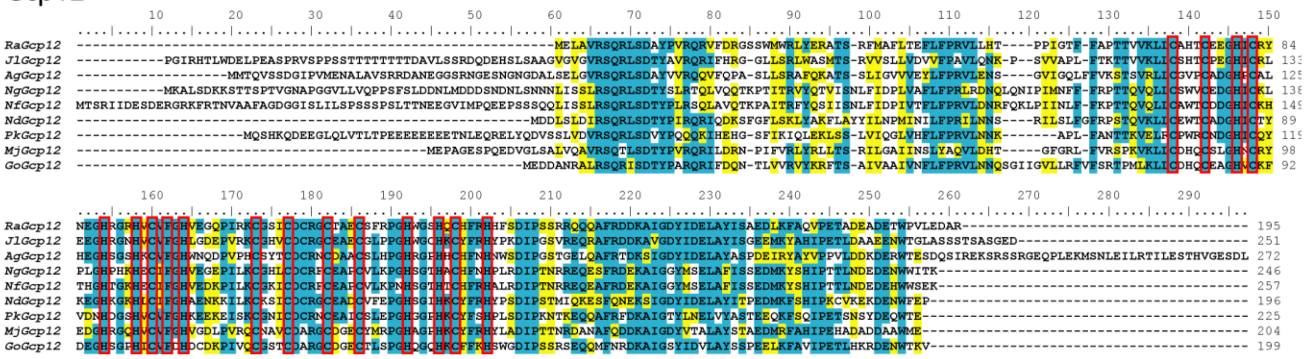

**Supplementary Fig. 12** Multiple sequence alignments of Gcp6 and Gcp12 proteins. Absolutely conserved histidine and cysteine residues (indicative of possible binding of a prosthetic group by the proteins) are framed by red rectangles. Identical and similar residues are highlighted in turquoise and yellow, respectively (using 50% conservation of the position as the threshold). Species abbreviations: *Mj – Malawimonas jakobiformis*, *Go – Gefionella okellyi*, *Ra – Reclinomonas americana*, *Ag – Andalucia godoyi*, *Nd – Neovahlkampfia damariscottae*, *Pk – Pharyngomonas kirbyi*, *Ng – Naegleria gruberi*, *Nf – Naegleria fowleri*, *Jl* – Jakoba libera.

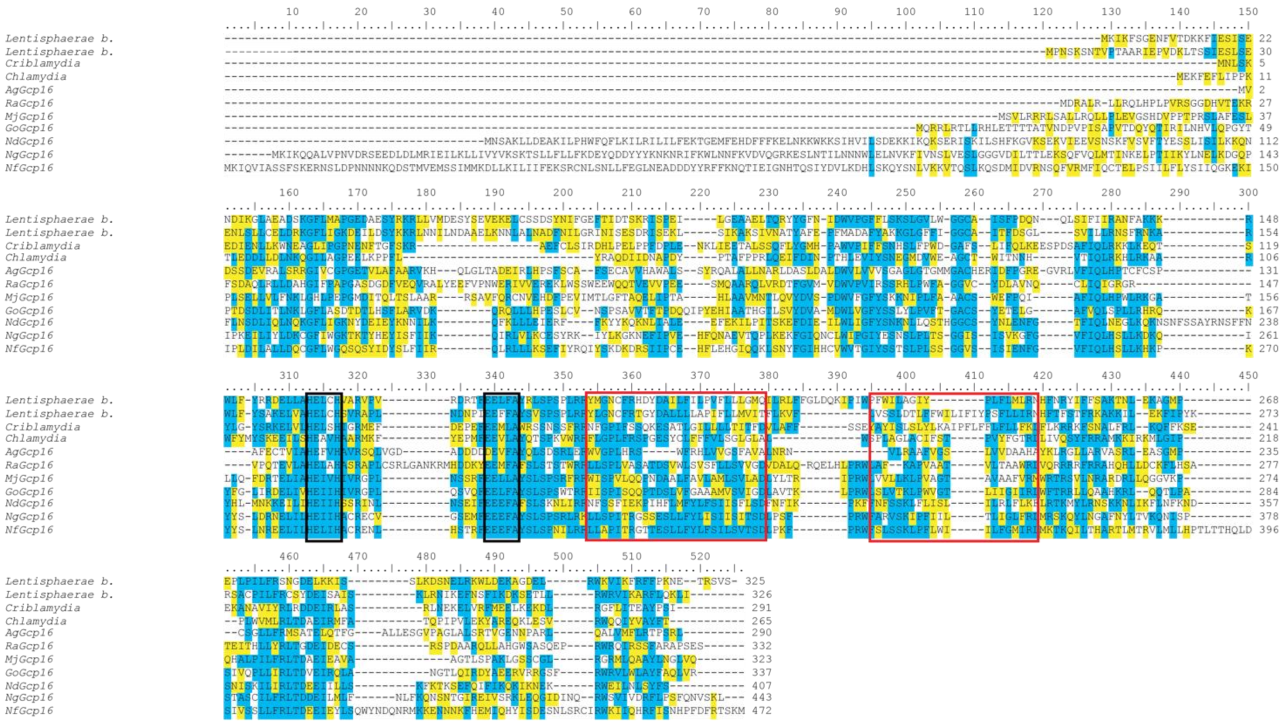

**Supplementary Fig. 13** Multiple sequence alignment of the eukaryotic Gcp16 proteins and selected bacterial homologues. Regions including predicted transmembrane domains are framed by red rectangles. Metallopeptidase motifs HEXXH and EXXA are highlighted by the black rectangle. Putative mitochondrial targeting presequences predicted by MitoFates are underlined. Identical and similar residues are highlighted in turquoise and yellow, respectively (using 50% conservation of the position as the threshold). Species abbreviations: *Mj – Malawimonas jakobiformis*, *Go – Gefionella okellyi*, *Ra – Reclinomonas americana*, *Ag – Andalucia godoyi*, *Nd – Neovahlkampfia damariscottae*, *Pk – Pharyngomonas kirbyi*, *Ng – Naegleria gruberi*, *Nf – Naegleria fowleri*.

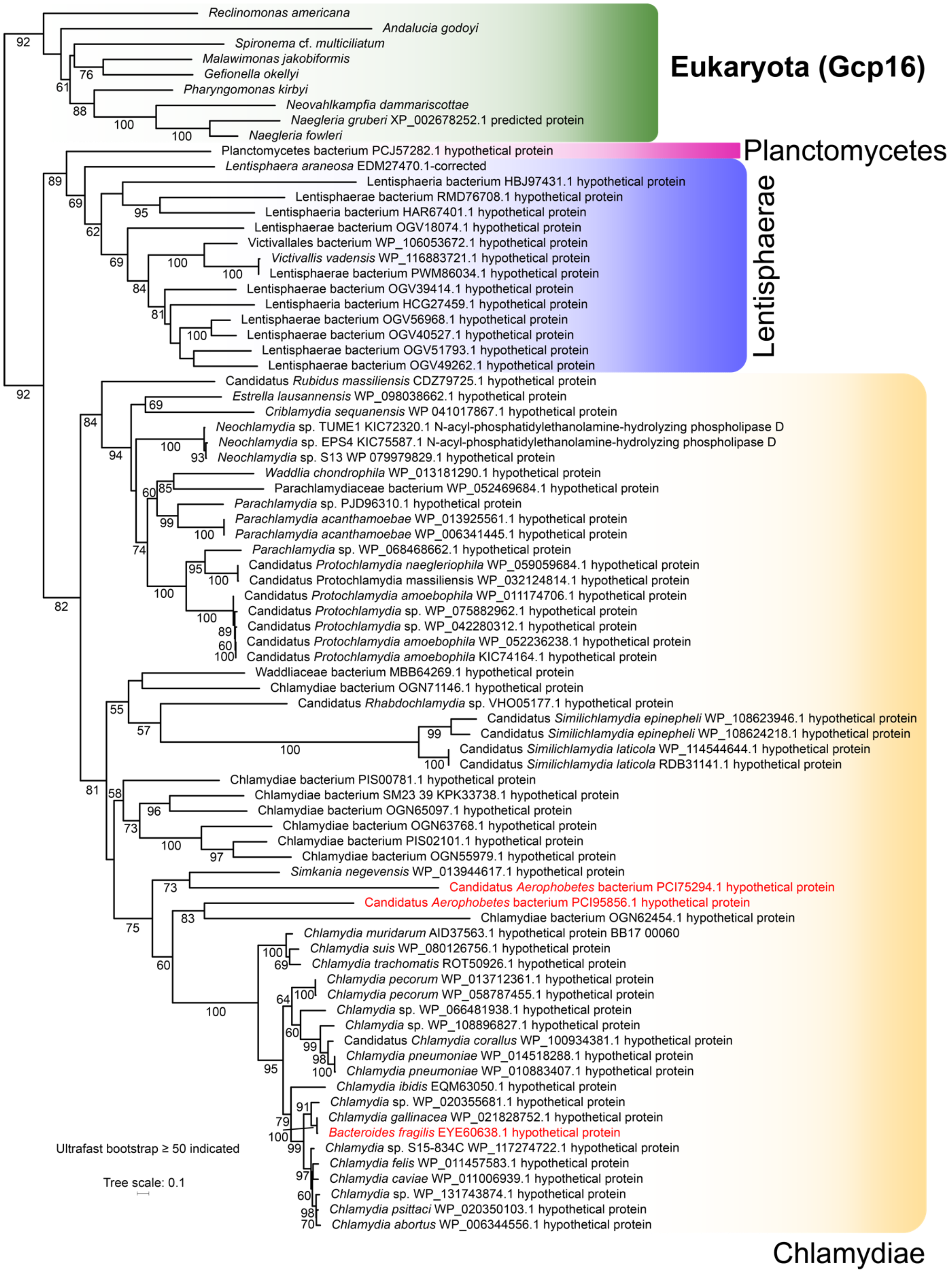


**Supplementary Fig. 14** Phylogenetic analysis of the Gcp16 protein. The ML tree (IQ-TREE, substitution model LG+R4, 1000 ultrafast bootstraps) includes all Gcp16 homologues identified in databases (using an exhaustive psi-blast search of the NCBI nr protein database and targeted blast searches of genomic and transcriptomic data from other repositories). The three sequences highlighted in red come from apparently misidentified taxa or contaminated genome assemblies. The sequence EYE60638.1 is encoded by a gene on a >800 kbp contig attributed to *Bacteroides fragilis* str. S6L5, i.e. a member of the phylum Bacteroidetes, but blast searches with proteins encoded by randomly picked genes from different regions of the contig all give as identical hits sequences from *Chlamydia gallinacea*. The two sequences attributed to two different isolates of an “Candidatus Aerophobetes bacterium” (i.e. members of the phylum Candidatus Aerophobetes) are located on contigs encoding proteins that exclusively (in case of PCI75294.1) or predominantly (PCI95856.1) retrieve sequences from the phylum Chlamydiae as best non-self hits, hence are most likely derived from unidentified representatives of the latter phylum.
