## Supplementary methods for "Analysis of diverse eukaryotes suggests the existence of an ancestral mitochondrial apparatus derived from the bacterial type II secretion system"

**Sequencing and assembly of *Neovahlkampfia damariscottae* and "*Malawimonas californiana*" genomes**

*Neovahlkampfia damariscottae* CCAP 1588/7 (obtained from the Culture Collection of Algae and Protozoa; https://www.ccap.ac.uk/) was grown for 5 days in 20 Corning® cell culture flasks (surface area 25 cm2) with 10 ml of ATCC 1525 medium. Since the culture contained various bacteria, we incubated cells overnight in 10 ml of fresh ATCC 1525 medium with 166 µl of antibiotics prepared as a mixture of ampicillin (100 µl), streptomycin (500 µl), penicillin (250 µl), and kanamycin (50 µl). Genome DNA was extracted using Qiagen DNeasy Blood & Tissue Kit following the manufacturer´s instructions. Genomic DNA was sequenced on Illumina Miseq 2x250bp platform. The total of 27,972,096 reads were trimmed by Trimmomatic (Bolger et al. 2014). An initial assembly was built with Spades 3.10.1 (Bankevich et al. 2012) and the scaffolds were clustered into different bins by MaxBin (Wu et al. 2014). Because the average GC content of the *N. damariscottae* genome is low (27.4% in the final assembly), bins with a relatively high GC content (> 40%) were inspected manually and after confirmation of their bacterial origin, they were removed from the assembly. Trimmed reads were mapped back to the cleaned assembly with Bowtie2 (--very-sensitive-local settings; Langmead and Salzberg 2012) and 14,663,323 mapped reads were used for the final assembly, which was built with Spades 3.10.1 using the "careful" option. The final assembly consists of 1,865 scaffolds with the cumulative length of 21.5 Mb, and the N50 value of 95,418 bp. It was deposited at GenBank with the accession number JABLTG000000000.

"*Malawimonas californiana*" (ATTC 50740) is a formally undescribed malawimonad that is cultivated with live *Enterobacter aerogenes* (ATCC 13048) as a food source (detailed recipes for the media are described at http://megasun.bch.umontreal.ca/People/lang/FMGP/methods.html). The large variety of food bacteria that was present in the original strain was reduced by repeated dilutions in 100 mL growth medium plus live *E. aerogenes*, so as to retain only a few malawimonad cells, and grown to the early stationary phase. The final isolate with less bacterial contaminants (after five growth cycles) is being kept for long-term storage under liquid nitrogen. For DNA purification, aliquots of the stock culture were added to fresh medium (500 mL, 2.5 L Erlenmeyer flasks) containing ~200 mg pre-cultured live *E. aerogenes* cells. Cultures were gently shaken at 22°C and daily supplemented with live bacteria, provided that most bacterial cells were consumed. Cells were harvested by centrifugation in the early stationary growth phase (after 2-5 days), at a point when most food bacteria were consumed. Harvested cells were lysed in a Tris-EDTA buffer containing 0.2 % SDS plus 100 µg/ml proteinase K, dialyzed for 24 hr against the same buffer, and then further purified by CsCl-*bis*benzimide equilibrium gradient centrifugation (Lang and Burger 2007). GS FLX Library Preparation Method protocols (Roche) were used to prepare shotgun and paired-end (3 Kb and 8 Kb) libraries for sequencing with the 454 method on the Titanium platform. Altogether, 2,955,688 shotgun reads, 1,082,625 reads from the 3 Kb paired-end libraries, and 1,100,119 reads from the 8 Kb paired-end libraries were generated and assembled using Newbler 2.8 (Roche), yielding a draft genome assembly consisting of 1,123 scaffolds with the cumulative length of 51.4 Mb, and the N50 value of 399,050 bp. The assembly was annotated by using Augustus (Hoff and Stanke 2013), yielding 13,534 predicted protein sequences. The assembly and the predicted protein sequence set are available at http://megasun.bch.umontreal.ca/Malawimonas_californiana. Note that the assembly is not bacterial contamination-free and is expected to be to a certain degree inaccurate in homopolymeric regions, owing to the known limitation of the 454 sequencing method. Work on a more accurate assembly and annotation is underway and the outcomes will be published elsewhere.

**Sequence curation**

Existing Gsp and Gcp gene models (public for the *N. gruberi* genome and private for *A. godoyi* and the two malawimonads) were evaluated by considering transcriptome data and sequence conservation within the respective gene families, and modified if necessary. One of the *N. gruberi* genes proved to be affected by an assembly error in the existing genome assembly; this was corrected by consulting the original Sanger sequencing reads. For genes without existing gene models (i.e. genes missed by automated annotation programs or residing in as-yet unannotated genome assemblies), the respective gene models were built manually *de novo* using the same source of supporting information. For *R. americana* only three alternative fragmented genome assemblies, often with redundant highly similar contigs presumably representing alternative allele variants, were available; hence, transcriptome data were used to link separate contigs containing different parts (exons) of the genes. The resulting gene assemblies may thus be chimeras of different alleles.

Only transcriptomic data were available for *P. kirbyi* and the three *Percolomonas* spp., so the Gsp and Gcp protein sequences were deduced by conceptual translation from the appropriate reading frames of the relevant transcript contigs. In several cases the original contigs from transcriptome assemblies proved to include a truncated coding sequence (CDS); in most of these cases a complete CDS could be obtained by manual iterative extension of the truncated ends by using raw Illumina reads identified by blastn searches and considering linking information provided by paired-end reads. Note that for one of the *Percolomonas* strains, the transcriptomic data are available only as contamination in the transcriptome assembly attributed to the diatom *Nitzschia* sp. ChengR-2013 (GenBank accession number GBCF00000000.1). The taxonomic assignment of the respective Gsp/Gcp sequences to *Percolomonas* rather than *Nitzschia* is based on the presence in the assembly of a 18S rRNA sequence phylogenetically close to a sequence from *Percolomonas cosmopolitus* strain WS (Supplementary fig. 1) and considering the fact that homologues are absent from all other diatom (even stramenopile) transcriptomic and genomic assemblies.

In case of Gcp4 a homologue was found in a transcriptome assembly attributed to the moss *Bryum argenteum* (GenBank accession number GCZP00000000.1), i.e. an organism lacking other Gsp and Gcp homologues. The sequence is most likely a contaminant coming from a heterolobosean, specifically a V*ahlkampfia* sp., based on the presence in the assembly of a partial 18S rRNA sequence (accession number GCZP01036684.1) exhibiting 99% identity to the 18S rRNA sequence from *Vahlkampfia avara* strain 4171L. It is likely that *Vahlkampfia*, like many other heteroloboseans, exhibits a full set of Gsp/Gcp homologues, which are not represented in the assembly GCZP00000000.1 due to an insufficient coverage of the contaminating organism.

**Defining orthogroups for phylogenetic profiling**

In order to classify eukaryotic genes into putative groups of orthologous genes (orthogroups) for the purpose of identification of genes co-occurring with initially identified Gsp homologues in eukaryotes (phylogenetic profiling), two slightly different rounds analyses were performed. They differed in taxon sampling (Supplementary Table 6; the first round lacking protein sequences from *G. okellyi* and *Monocercomonoides exilis*), sequence similarity thresholds (see step p1 below) and the way how week connections between sequence clusters were dealt with (see step n3 below).

The analysis included two main phases. In the first, orthologous relationships of each protein to all others were inferred using the following custom bioinformatics pipeline (steps p1 to p4 were repeated for each protein):

_ (p1) the protein was blasted (BlastP) against the entire set of proteins using a threshold e-value ≤ 1e-8 and the BlastP alignment overlapping at least 35 % of the query (first analysis) or minimal e-value threshold of 1e-5 and a the BlastP alignment overlapping at least 25 % of the query (second analysis).

_ (p2) the seven best BlastP hits of each proteome were retrieved, aligned with Clustal Omega (Sievers et al. 2011) under default parameters, and the resulting alignment was trimmed using trimAl (Capella-Gutiérrez et al. 2009) by removing columns with gaps in more than 50% of sequences.

_ (p3) a phylogenetic tree was built from the trimmed alignment using FastTree (Price et al. 2010) under default parameters.

_ (p4) orthologous relationships were inferred from the mid-point rooted phylogenetic tree using the specie-overlap method implemented in ETE (Huerta-Cepas et al. 2010).

All orthologous relationships obtained from these phylogenetic analyses were combined to build an undirected graph in which vertices represent proteins, edges represent observed orthologous relationships between the two connected proteins, and connected components represent orthologous groups. This complex and dense network was then simplified using these successive steps:

_ (n1) edges connecting two proteins that have been found orthologous only once were discarded.

_ (n2) for each edge, we calculated the ratio “number of times the two proteins have been found orthologous) / (number of times the two proteins have been found paralogous”. Edges with the ratio lower than 0.5 were discarded.

_ (n3) bridges, identified as edges connecting two vertices that have at least four neighbours and that do not share a connected vertex, were discarded. This step was applied only in the second analysis.

_ (n4) within each orthologous group, gene-fusions were identified and the orthologous group was corrected as follows:  if the removal of a vertex led to the formation of two isolated connected components of at least three vertices, the vertex was considered as a gene fusion (genuine or artificial) and was added to each of the two newly formed connected components. To validate the approach, we then inspected 30 randomly chosen orthologous groups (including some belonging to complex gene families) and found rare false positive and false negative cases. Additionally, 20 randomly chosen proteins identified by our pipeline as gene fusions (an average of 31 gene fusions were inferred per proteome) were all confirmed by manual BlastP analyses.

**References to Supplementary Methods**

Bankevich, A., Nurk, S., Antipov, D., Gurevich, A. A., Dvorkin, M., Kulikov, A. S., Lesin, V. M., Nikolenko, S. I., Pham, S., Prjibelski, A. D. and Pyshkin, A. V. SPAdes: a new genome assembly algorithm and its applications to single-cell sequencing. *Journal of Computational Biology* **19,** 455–77 (2012).

Bolger, A. M., Lohse, M., Usadel, B. Trimmomatic: A flexible trimmer for Illumina sequence data. *Bioinformatics* **30,** 2114–20 (2014).

Capella-Gutiérrez, S., Silla-Martínez, J. M. & Gabaldón, T. trimAl: a tool for automated alignment trimming in large-scale phylogenetic analyses. *Bioinformatics* **25,** 1972–3 (2009).

Hoff, K. J. & Stanke, M. WebAUGUSTUS--a web service for training AUGUSTUS and predicting genes in eukaryotes. *Nucleic Acids Research* **41**, W123–8 (2013).

Huerta-Cepas, J., Dopazo, J. & Gabaldón, T. ETE: a python Environment for Tree Exploration. *BMC Bioinformatics* **11,** 24 (2010).

Lang, B. F. & Burger, G. Purification of mitochondrial and plastid DNA. *Nature Protocols* **2,** 652–60 (2007).

Langmead, B. & Salzberg, S. L. Fast gapped-read alignment with Bowtie 2. *Nature Methods* **9**, 357-359 (2012).

Price, M. N., Dehal, P. S. & Arkin, A. P. FastTree 2 – approximately maximum-likelihood trees for large alignments. *PLoS One* **5,** e9490 (2010).

Sievers, F. *et al.* Fast, scalable generation of high-quality protein multiple sequence alignments using Clustal Omega. *Molecular Systems Biolology* **7,** 539 (2011).

Wu, Y. W., Tang, Y. H., Tringe, S. G., Simmons, B. A. & Singer, S. W. MaxBin: an automated binning method to recover individual genomes from metagenomes using an expectation-maximization algorithm. *Microbiome* **2,** 26 (2014).
